# A Suite of Designed Protein Cages Using Machine Learning Algorithms and Protein Fragment-Based Protocols

**DOI:** 10.1101/2023.10.09.561468

**Authors:** Kyle Meador, Roger Castells-Graells, Roman Aguirre, Michael R. Sawaya, Mark A. Arbing, Trent Sherman, Chethaka Senarathne, Todd O. Yeates

**Affiliations:** Department of Chemistry and Biochemistry, University of California, Los Angeles, CA, USA 90095; UCLA-DOE Institute for Genomics and Proteomics, Los Angeles, CA, USA 90095

**Keywords:** symmetry, self-assembly, protein cages, protein-protein docking, *de novo* interface design, machine learning, cryoEM

## Abstract

Designed protein cages and related materials provide unique opportunities for applications in biotechnology and medicine, while methods for their creation remain challenging and unpredictable. In the present study, we apply new computational approaches to design a suite of new tetrahedrally symmetric, self-assembling protein cages. For the generation of docked poses, we emphasize a protein fragment-based approach, while for *de novo* interface design, a comparison of computational protocols highlights the power and increased experimental success achieved using the machine learning program ProteinMPNN. In relating information from docking and design, we observe that agreement between fragment-based sequence preferences and ProteinMPNN sequence inference correlates with experimental success. Additional insights for designing polar interactions are highlighted by experimentally testing larger and more polar interfaces. In all, using X-ray crystallography and cryo-EM, we report five structures for seven protein cages, with atomic resolution in the best case reaching 2.0 Å. We also report structures of two incompletely assembled protein cages, providing unique insights into one type of assembly failure. The new set of designed cages and their structures add substantially to the body of available protein nanoparticles, and to methodologies for their creation.

## Introduction

Recent advances in protein design are making it possible to engineer self-assembling protein architectures of high complexity ^1–3^. The seminal work by Padilla et al. in 2001 ^4^ laid the foundation for creating novel protein cages and other extended materials by exploiting principles of symmetry. The key idea was that bringing two different symmetry elements together in a precisely defined arrangement is sufficient to dictate formation of surprisingly diverse and complex assembly outcomes. Such symmetry combination materials (SCM) are ideally suited to benefit from recent advances in computational protein design. Considering just two-component types, 124 different architectural forms have been articulated mathematically ^5^. And for each such symmetry form, thousands of different pairs of protein oligomers comprise potential building blocks, leading to an extraordinarily deep space for design; the introduction of *de novo* protein components further expands the space. Importantly, only a sliver of this design space has been explored experimentally to date. Large machine learning models are enabling far more efficient search of protein conformational space ^3,6–9^ and effectively applying these tools to model protein materials will be key in the ultimate realization of their vast potential.

Achievements over the last decade have demonstrated the potential value of engineered protein cages and related types of SCMs for applications across the areas of nanotechnology and medicine. Their biocompatibility as well as their size, topology, and multivalency have enabled applications such as the localization of target substrates ^10^, molecular delivery ^11^ or sequestration of payloads ^12^, and scaffolding of antigens ^13–15^, enzymes ^16,17^, or binders for high resolution imaging ^18,19^. Notwithstanding these promising demonstrations, successful designs have only scratched the surface with respect to the rich functional complexity and dynamics that are possible with protein assemblies, as exemplified by naturally evolved systems ^20–25^. Indeed, emerging efforts to mimic the complex behaviors of natural systems are leading to exciting new design prospects ^26,27^.

Crucially, the predictable design of novel protein cages and similar materials remains a challenge ^28^. While a first approach for designing protein cages used genetic fusion ^4,29^, subsequent work (beginning with King et al. 2012) has increasingly relied on the computational design of *de novo* non-covalent interfaces between protein subunits. A critical factor therefore in successful material design is the specification of a geometrically precise interface that is accessible and cooperative with the surrounding energy landscape ^30^. The interface must encode sufficient information to drive the emergence of quaternary structure ^31^, while not corrupting the energetics governing tertiary structure ^32^. Importantly, diverse SCM’s including cubic and icosahedral protein cages, can be created through the installation of a single interface type between candidate building blocks. Of course, only a minute fraction of possible protein pairs and orientations constitute suitable starting points for designing a new interface through amino acid substitutions, so identifying plausible protein backbone poses is an important algorithmic consideration for SCM approaches.

In the present study, we evaluate two new elements as part of an improved protocol for designing protein cages. Addressing the challenge of generating protein interfaces resembling those in native complexes, we employ a protein fragment-based method that identifies design poses with backbones exhibiting common modes of association in nature ^5^. With regard to amino acid sequence design at the new interface, we first designed cages using knowledge-based scoring functions. Spurred by those results, we pursued a new approach based on recently developed machine learning methods to achieve greater success, implementing the ProteinMPNN graph neural network in a unified workflow. Through a combination of biochemical and detailed structural characterization, we validate a suite of designed two-component tetrahedral cages to substantially expand the set of engineered protein nanomaterials. A comparison of design parameters with patterns of success or failure (including in past efforts) offers insights into challenges at the frontier of protein design. The new symmetric materials, design descriptions, and accompanying software serve as a foundation to explore the synthesis of a growing universe of protein materials.

## Results and Discussion

### Defining the target assembly

As a design target, we chose a tetrahedral architecture denoted by T:{C3}{C3}. In materials of this form, two different trimeric protein components with C_3_ symmetry come together, with four copies of each trimer giving an overall stoichiometry of A_12_B_12_. Each trimer is oriented so its 3-fold axis is coincident with a body diagonal of a cube (Figure 1b). As the degrees of freedom available to form the tetrahedron are rather strict, key to producing bonafide T:{C3}{C3} assemblies is the accurate modeling of a *de novo* protein-protein interface between the surfaces of trimeric building blocks to enforce a strict intersecting angle of the two trimeric axes. If the interface between the pair of trimers is accurately modeled — *i*.*e*. if they associate strongly and with the correct geometry — four copies of each trimer will assemble to form a mid-nanometer scale, symmetric protein cage (Figure 1a).

**Figure 1.**
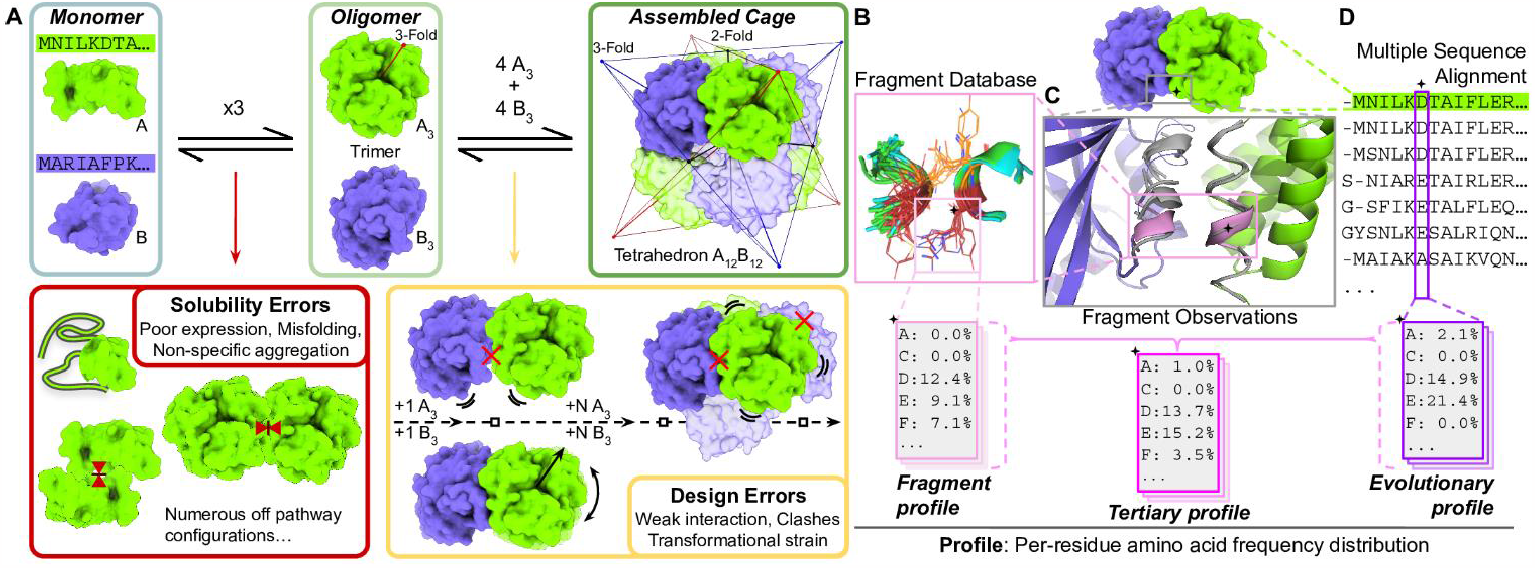
A framework for interface design and amino acid profile constraints for creating protein cages. **a)** A diagram of assembly states and errors leading to failure. The two-component tetrahedral assembly is represented by the multi-colored stellar octahedron, where each color constitutes one tetrahedron. **b-d)** For the pose, fragment observations featurize the nascent interface, both for interaction potential during docking and for sequence design (panel c). Fragment Observations are depicted as gray ribbons which overlay ribbons colored according to each component. A fragment observation of interest is indicated in pink. The corresponding cluster of structurally similar fragments from the Fragment Database is utilized to calculate amino acid frequencies for all positions of the cluster (panel b). At the starred position, all fragment observations contribute to an amino acid frequency distribution, which is analyzed at all residues to cumulatively produce a Fragment Profile. **d)** The initial sequence is used to search for aligned sequences. At the starred position, a frequency distribution of evolutionarily observed amino acids is calculated. The collection of frequency distributions at every residue constitutes the Evolutionary Profile. The fit of fragment observations at each position in the pose modulates the extent to which a Tertiary Profile reflects the Fragment Profile or the Evolutionary Profile.

In generating candidate docked positions (*i*.*e*. poses) consistent with tetrahedral designs, we restricted the interface search space to native like conformations, adopting the fragment-based docking approach implemented in the program Nanohedra ^5^. This approach analyzes the surface-exposed segments of two protein components and systematically identifies poses that bring segments from the two components into arrangements commonly observed in known protein-protein interfaces (Figure 1c). We performed all-to-all pairwise docking between 84 different trimeric structures from the PDB with ∼377,000 total poses output from 3,584 pairs of trimers (see Methods). To ensure the prioritization of interfaces composed of more extensive tertiary motifs, we filtered the complete set of 377,000 poses, utilizing fragment observations from three or more discrete secondary structure elements in the nascent interface, resulting in ∼45,000 qualifying poses. We gathered coarse energy and area specific metrics (see scouting protocol Methods) and retained poses with greater than 1,200 Å buried surface area (BSA) in the complex, a negative calculated interface free energy, and less than 3 Å backbone root mean squared deviation (RMSD) between the initially docked pose and an energy minimized model. Finally, we used overall shape-based features (see Methods) as an additional filter to arrive at 590 candidate poses.

### Interface design using fragments and knowledge-based hydrogen bond networks

We frame our interface design ideas around a three state model of protein folding and association ^33^ (Figure 1a). The first state consists of protein monomers. In the second state, multiple protein chains are associated to form one oligomer. Importantly, designs that reach the discrete oligomeric state avoid multiple off-pathway states that together constitute *solubility errors*. Well-formed oligomers serve as the precursors for the third state, the full protein-protein complex, where sets of oligomers are further associated in a specific geometry to form a higher-level architecture. Herein, the complex state constitutes a full tetrahedral protein cage. When a complex is not formed successfully from viable oligomers, the result constitutes some type of *design error*, reflecting a failure to establish the intended interface. To ensure the experiment would provide feedback on the interface design task, which is critical for improving *de novo* interface design techniques ^34,35^, we aimed in our design strategies to most strongly limit solubility errors, *e*.*g*. by seeking to avoid undue hydrophobicity.

Our fragment-based docking (Nanohedra program) includes statistical information on amino acid preferences for tertiary structure (fragment-fragment) motifs ^36^. We therefore sought to utilize this information to bias amino acid selections during sequence design. We calculated position-specific amino acid frequencies for every fragment residue in the nascent interface, collectively referred to as the fragment profile (Figure 1b, see Methods). Similarly, an evolutionary profile was calculated from position-specific amino acid frequencies observed from multiple sequence alignments (MSA) of homologous proteins (Figure 1d, see Methods). These two profiles were combined into the tertiary profile, which represents an amino acid distribution conditioned on the tertiary structure of the nascent interface and the underlying protein folds.

Early in method development, we observed that hydrophobic amino acids tended to be favored during design owing to Lennard-Jones score terms ^37^ and the default FastDesign ramping protocol ^38^. As the fragment database utilized for pose identification contains motifs found in interfaces from native complexes, and those complexes typically demonstrate greater polar characteristics ^39^, we hypothesized that explicitly sampling polar amino acids could reduce the prevalence of hydrophobic atomic interactions while retaining well-packed and complementarity tertiary motifs ^40^. To utilize fragment information for design and sampling, we developed a modified HBNet protocol ^41^, using fragment information to guide the HBNet search. We refer to this protocol as FragmentHBNet (see Methods). Briefly, interface residues with fragment observations initiate the HBNet search for low energy hydrogen bond networks. In a subsequent step, the fragment profile is used to constrain amino acid sampling during packing and minimization at fragment residues around the hydrogen bond network residues. These dual searches effectively prioritize the most well packed cores that support low energy hydrogen bonding networks. Finally, the tertiary profile is used to guide sampling of the entire interface. Interface design trajectories with FragmentHBNet affected several key interface design metrics, including lowering contributions from hydrophobic buried surface area (BSA) while increasing the number of hydrogen bonds.

The 590 candidate poses noted above were each subjected to the FragmentHBNet design protocol, producing a designed structure for the top 20 trajectories. All designs were then filtered for shape complementarity > 0.68, measured BSA > 1,000 Å, fraction of hydrophobic BSA < 65%, buried unsatisfied hydrogen bonds < 3/1,000 Å^2^ of BSA, and no residues with highly unusual atomic contact patterns (as judged by an error score greater than 2 standard deviations in the program Errat ^42^). One design from each pose was prioritized and a mutation reversion protocol was performed (see Methods). From all reverted and original designs, we again prioritized one design per pose and finally manually inspected each design for missing segments near interfaces; poorly suited termini or unmodeled regions in the proximity of the interface were discarded. Lastly, poses were clustered for similarity according to iAlign and 41 designs were selected for testing. We refer to this design set using the FragmentHBNet protocol as T33-fn.

### Characterization of T33-fn Designs

For biochemical characterization, we added a polyhistidine affinity tag (‘His Tag’) onto an exposed terminus of one of the two components and generated codon-optimized genes for simultaneous expression of both components in *Escherichia coli* (see Methods). With the His Tag attached to only one component, we expected to purify both components if the interface design was successful. After gene synthesis, *E. coli* cells were transformed, and the designs were expressed and purified by immobilized metal affinity chromatography (IMAC). SDS-PAGE analysis of the lysates revealed 35 cases wherein both genes were well-expressed. Out of these 35, both components were in the soluble fraction after clarifying the lysate by centrifugation in 10 cases. Eight of the designs demonstrated the expected co-elution of both subunit types, though with some variation in relative abundance, and a tendency toward higher quantities of the His-Tagged trimer (Figure S1a). For all eight designs, we concentrated the elution fractions with both components present and isolated the assemblies using size exclusion chromatography (SEC). The chromatograms indicated significant assembly for a single design (Figure 2), while the other seven revelaed component trimers, some larger fractions containing both components, and limited amounts of assembly-sized species (Figure S1b).

**Figure 2.**
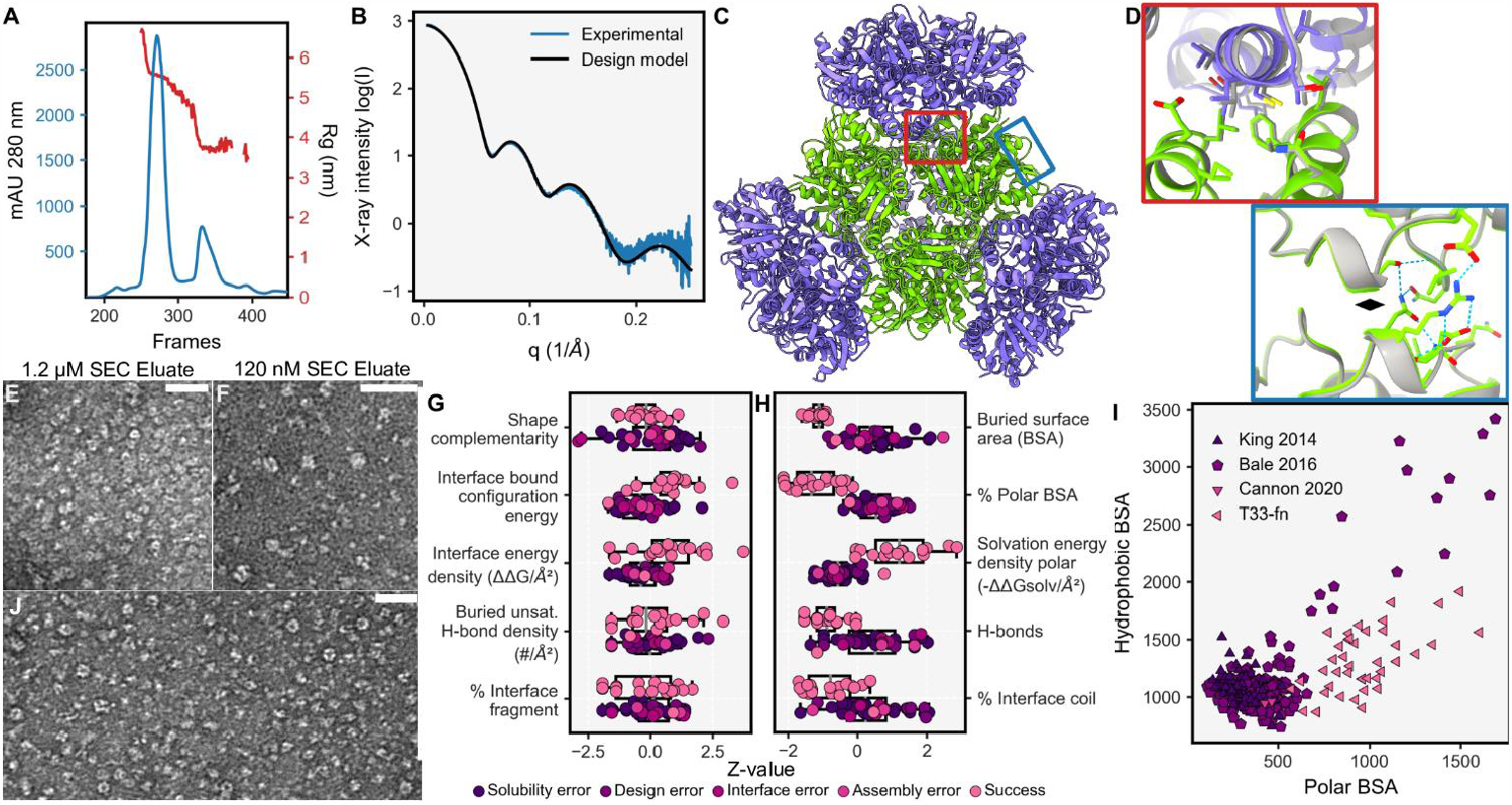
Biophysical and structural analysis of T33-fn10, a cage designed using knowledge-based algorithms. **a)** SEC-SAXS of T33-fn10 giving a measured Rg of 5.6 nm for the assembled cage. **b)** A SAXS X-ray scattering plot averaged over the fractions corresponding to cage species (blue) and a theoretical plot from the designed model (black). **c)** A crystal structure of T33-fn10 colored according to component trimer type. **d)** A close-up view of the designed interface, split into two regions. The black diamond represents the location of a two-fold axis of symmetry between separate green trimers. **e-f)** Micrographs of T33-fn10 at 1.2 μM and 120 nM respectively, showing disassembly upon dilution of particles. Scale bars are 50 nm. **g-h)** A comparison of T33-fn design metrics (top group) with prior 2-component SCM successful designs (bottom group) for important design filters (panel g); and protocol specific differences (panel h). Individual designs are colored according to design outcome. **i)** Distribution of buried surface area (BSA) according to atomic polarity for various 2-component SCM designs colored by publication. Markers indicate the design symmetry (triangle - Tetrahedral, pentagon - Icosahedral). **j)** A micrograph of T33-fn40 demonstrating cages after purification by SEC. The scale bar is 50 nm.

For the most well-behaved design, T33-fn10, biochemical characterization revealed properties expected from the design model (Figure 2). SEC coupled to small angle X-ray scattering (SAXS) demonstrated excellent fit to the design with a measured radius of gyration (Rg) of 5.55 nm compared to the designed value of 5.53 nm (Figure 2a,b). Additionally, we obtained crystals that diffracted to modest resolution, with the best dataset reaching 6 Å. A crystal structure was solved by molecular replacement using the design model (see Methods) revealing eight copies of each monomer in the asymmetric unit (ASU). Upon application of crystal symmetry operators, two separate tetrahedral assemblies are recapitulated, confirming the designed interface (Figure 2c,d). Both instances of the assembly show excellent agreement to the design model with mean values of 0.8 Å RMSD over all C-alpha atoms and a value of 0.99 for the assembly local distance difference test (LDDT) (Table 1). Validation by negative stain transmission electron microscopy (TEM) revealed that specimens prepared at 1.2 μM of assembled complex appeared crowded but mostly homogenous, while specimens prepared at 120 nM lack clear evidence of cage-like assemblies (Figure 2e,f). The designed interface results in apparently intact assemblies at concentrations in excess of 1 μM, but with limited assembly at lower concentrations.

**Table 1.** Structurally validated assembly statistics:

| Design Name | Resolution (Å) | Assembly RMSD (Å) | ASU RMSD (Å) | Assembly LDDT |
| --- | --- | --- | --- | --- |
| T33-fn10 | 6.0 | 0.75 | 0.45 | 0.99 |
| T33-ml23 | 2.0 | 2.37 | 1.26 | 0.78 |
| T33-ml28 | 2.7 | 1.73 | 1.08 | 0.86 |
| T33-ml30 | 4.2 | 3.62 | 1.88 | 0.69 |
| T33-ml35 | 2.9 | 1.55 | 1.11 | 0.85 |
| T33-ml23<br>A<sub>12</sub>B<sub>9</sub> | 3.9 | 2.60 | 1.45 | 0.73 |
| T33-ml35<br>A<sub>9</sub>B<sub>12</sub> | 4.4 | 1.55 | 1.42 | 0.85 |

Agreement between experimental structures and their respective design models. All structures are from cryo-EM except T33-fn10, which was from X-ray crystallography. ASU - Asymmetric unit, RMSD - root mean squared deviation, LDDT - local distance difference test.

Designs with unequal stoichiometry were more difficult to characterize. For one design, T33-fn40, TEM captured assemblies resembling the intended model in projection (Figure 2j), though heterogeneity was noted in SDS-PAGE of SEC fractions (Figure S1b). For the remaining six co-eluting designs, generally low yields of the non-His-Tagged components limited purification of fully assembled complexes.

### T33-fn interface design analysis

Notwithstanding the successful creation of a new protein cage, T33-fn10, the low success rate in the experimental trials overall, despite promising indications of soluble oligomers and evidence for their association (*i*.*e*. co-elution), indicated that most designs failed to assemble faithfully. To understand how design indications contribute to biochemical outcomes, we grouped the designs according to the classifications of fig 1a, – *i*.*e. solubility errors, success*, and *design errors* – and compared these outcomes to metrics calculated for the design models after Rosetta refinement (see Methods). We further separated design errors into three types of error classification: *design errors*, where both components are soluble yet don’t interact, *interface errors*, where the components coelute without higher order assembly, and *assembly errors*, where higher order species are observed, but no tetrahedral complex was observable. Finally, we included the successfully characterized two-component cage design models from King *et al*. 2014, Bale *et al*. 2016, and Cannon *et al*. 2020 ^40,43,44^, to provide further understanding of design requirements.

Important interface metrics, including buried unsatisfied hydrogen bond density ^35^, shape complementarity ^45^, interface bound configuration energy ^46^, and calculated free interface energy density ^47^ indicated all design were as favorable, if not more favorable than prior successful designs (Figure 2g). Areas of deviation highlight particular choices made in our design protocol, such as the number of hydrogen bonds, fraction of polar BSA, and calculated solvation free energy density (Figure 2h). A notable observation concerns the total BSA in the T33-fn design set. Our initial solvent accessible surface area (SASA) measurement using Rosetta underestimated BSA by ∼2-fold (see supplement). As a result, T33-fn had relatively large interfaces on average (2,287 Å^2^), which were substantially polar in character, exceeding total BSA and polar BSA values of previously designed protein cages (Figure 2i). Interestingly, T33-fn interfaces utilized extensive loop/coil secondary structure elements, for which it appeared difficult to design adequate atomic interactions (Figure 2h).

Our design choices in this experiment were guided in substantial measure by a desire to minimize outcomes where unduly hydrophobic interfaces might lead to solubility errors. While designs were generally soluble, retrospective analysis suggests that T33-fn10 was the single design to possess a solvation free energy in the range of prior successful designs. Though prior successes also had negative solvation energy values, where negative indicates that aqueous solvation (*i*.*e*. disassociation) is favored, T33-fn selection factors manifested in more negative values for interface solvation free energies overall (Figure S2a,b). Though numerous other factors are undoubtedly important, many of our designs likely lacked crucial contributions of hydrophobicity towards interface formation. The challenges in optimizing parameter choices and protocols, clearly highlighted by this exercise, led us to incorporate machine learning techniques into a new protocol for designing protein cages.

### Fragment-guided protein cage design using machine learning

In a second set of designs, we revisited both docking and design methodologies, augmenting each step with recent developments in machine learning, where available. For docking, we improved upon techniques for building block input and search of the docking space. For sequence design, we implemented the MPNN machine learning algorithm ^48^, based on graph neural networks. Finally, in filtering, we prioritized agreement between fragment observations and neural network outputs to prioritize sequences that agreed with native interface constraints and were predicted computationally to fold into the intended oligomeric components.

As a first improvement, structure prediction using AlphaFold was integrated into the design pipeline ^6^. In the early stages, we performed predictions to fill in unmodeled regions of the input oligomers; their absence in our earlier protocol led to uncertainty and tedious assessment of potential collisions. In a second step, we assessed whether designed sequences are predicted to fold into the oligomeric state. Both assessments improved confidence that selected designs would meet prerequisites for oligomeric and complex formation.

To improve upon identification of high-quality docked poses, we implemented three search heuristics in Nanohedra (see Methods). First, we searched for poses where the resulting interfaces had continuous fragment overlaps across multiple residues. This option locates fragment pairs only if they participate in additional paired observations, which embeds each qualifying residue into higher order relationships, *i*.*e*. a network of coordinating residues. Next, we clustered coarsely sampled poses into a smaller number of transformational groups, which reduced the search space while maintaining top docking candidates. Finally, each cluster was finely sampled along the rigid body degrees of freedom consistent within the symmetric architecture in order to identify poses with optimal docking metrics.

Again, we docked poses in the T:{C3}{C3} architecture resulting in 55,611 pairwise combinations of 334 AlphaFold-curated trimers. Poses that implicated 40 or more interface residues and three discrete secondary structures using fragments were selected for finer sampling using Nanohedra score optimization, resulting in 151,597 poses. After docking, we further filtered these poses by increasing the required number of discrete fragment secondary structures to include two secondary structure elements from each trimer in the interface (> four fragment secondary structures total). The remaining 72,419 poses represented plausible backbone models for a novel two-component protein cage.

We then utilized the deep graph based neural network implemented in ProteinMPNN ^7^ for the task of sequence design. Exploiting ProteinMPNN’s computational efficiency, we deeply explored the generative sequence space conditioned on each of the candidate poses, producing ∼3.7 million sequences. We performed 40 inference trajectories on the backbone coordinates of each pose in the fully symmetric assembly form, using three separate protocols for residue selection (see Methods). In the *interface* protocol, we specify that ProteinMPNN infers the amino acid sequence for residues at the new interface between components (comprising 10 trajectories). In the *interface + neighbors* protocol, ProteinMPNN infers the sequence of both interface residues and their neighbors (10 trajectories). In the *all* protocol, ProteinMPNN infers the amino acid at every possible residue (20 trajectories). Across the protocols, we observed the ProteinMPNN score, *i*.*e* the ProteinMPNN inference profile negative log loss, was dependent on the number of designed residues (Figure 3e), while increasing the number of interface residues had little effect (Figure S2c).

**Figure 3.**
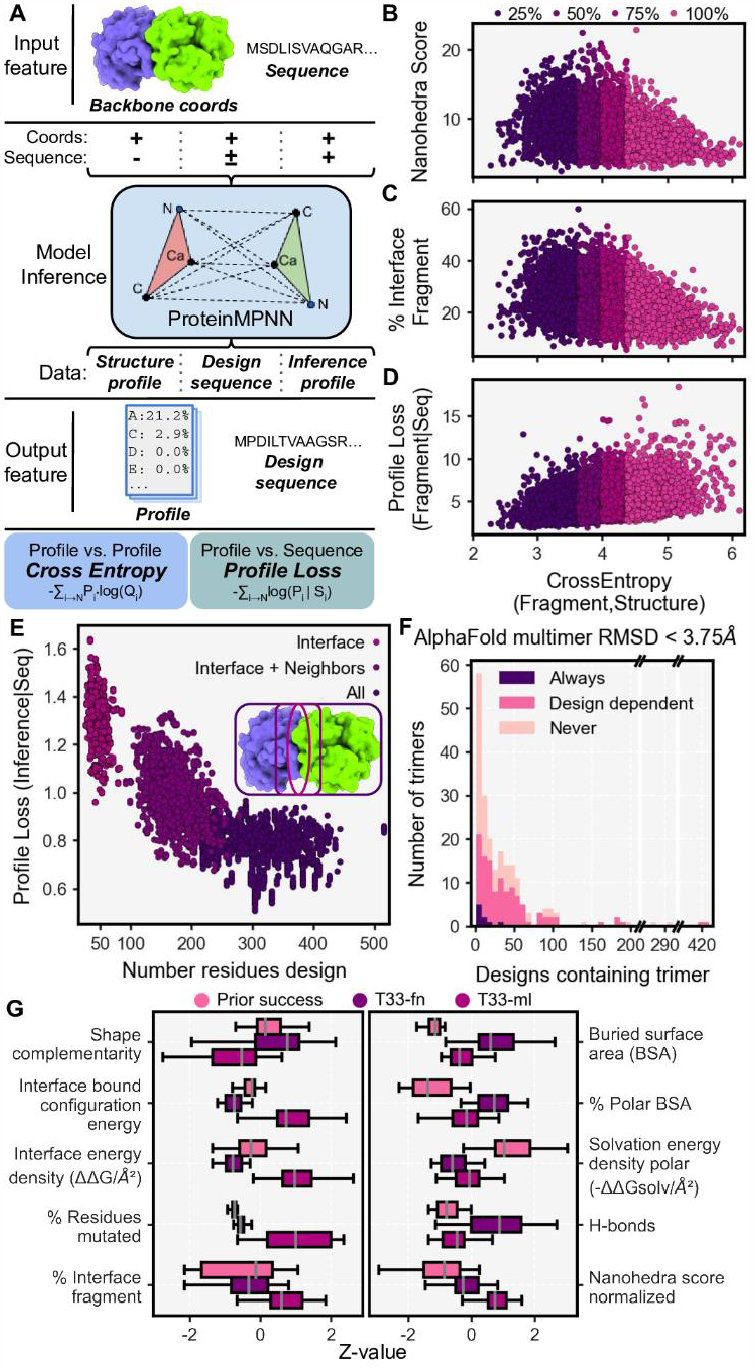
Characterization of sequences and poses for T33 machine learning (T33-ml) design models. **a)** ProteinMPNN model inference informs on different features of the sequence-structure relationship. Coordinates and sequence serve as input features to vary inference methods and therefore output features. Output features are used to measure cross entropy and profile loss to inform on the fit of inference to other distribution profiles. **b-d)** Comparison of the ProteinMPNN structure profile and fragment profile cross entropy vs Nanohedra score (panel b); percent of interface with fragment observations (panel c); and fragment profile loss given the designed sequence (panel d). **e)** The distribution of ProteinMPNN scores for designed sequences compared with the number of residues chosen for design based on three selection protocols. **f)** A stacked histogram of ensemble AlphaFoldInitialGuess folding outcome for designed sequences grouped according to each input trimer PDB identifier. **g)** Box plots showing the distribution of T33-ml design metrics scaled according to z-value compared to T33-fn and prior successful protein cages.

To explore how ProteinMPNN inference could be used to prioritize docked poses for experimental testing, we investigated the ProteinMPNN structure profile, *i*.*e*. the amino acid probability distribution inferred only from the pose backbone coordinates (Figure 3a). We compared this amino acid profile to the fragment amino acid profile for the pose (described above) calculating the cross entropy (CE) value between the two, with lower CE values equating to closer agreement between probability distributions. Lower CE correlated with a larger Nanohedra score (Figure 3b) and a higher percentage of the total interface represented by fragments (Figure 3c). Importantly, lower CE values correlated with lower fragment profile negative likelihood loss given the sequence, referred to as the fragment profile loss (Figure 3d). A lower value for the profile loss maximizes the likelihood that, for a single designed sequence, the sequence will encode the sequence-structure relationship embedded by fragment-based data. The agreement between fragment information and ProteinMPNN inference identifies poses for which sequences are more likely to encode structures that are compatible with the intended interface.

Given these trends, we filtered for poses with high quality fragment observations whose individual sequences exhibited a fragment profile loss less than five. Additionally, we filtered for sequences where *all* residues were designed, the ProteinMPNN score was less than one, and the recovery or retention of native sequence was at least 45%. We also implemented filters to ensure sequences designed in the complex state remain compatible with the oligomeric, unbound state. We filtered for total ProteinMPNN score in the unbound state less than 880 and an evolution profile loss (analogous to prior profile loss, but utilizing the evolutionary profile) less than 2.5 to avoid aberrant placement of hydrophobic residues.

For each of the 4,241 poses that passed these filters, we chose the single best designed sequence according to ProteinMPNN score and subjected it to predictive (computational) structural validation (see Methods). On one hand, we took the design sequence and threaded it onto the docked pose, mutating every residue to the designed amino acid and then evaluating structural features (see Methods: Threading sequences). On the other hand, using AlphaFold multimer, we performed folding predictions in the trimeric state to assess how strongly each sequence specified the proposed oligomer. We performed folding without MSA features and instead used the AlphaFoldInitialGuess variant to bias folding on the pose coordinates (see Methods, adopted from Bennett et al 2023) ^49^. We found many of the AlphaFold predictions recapitulate the intended trimeric forms in a sequence dependent manner. When grouped by building block identity, the majority of trimers (n=107) gave predicted oligomeric structures within 3.75 Å (C-alpha RMSD) of the known structure (Figure 3f). Further, we found many trimers failed to satisfy this threshold for any designed sequence (n=93), while few trimers folded correctly for every sequence analyzed (n=9).

Following predictive structural validation, we selected designs with shape complementarity >= 0.65, BSA >= 1,500 Å^2^, buried unsatisfied hydrogen bond density <= 2/1,000 Å^2^, and interface solvation free energy density >= -0.01/Å^2^. Finally, we applied two different filters using folding calculations. In the most selective case, we included designs where both trimers successfully folded to within 3.75 Å RMSD (n=248), resulting in 17 designs. In a more permissive case, we selected designs where only one component folded satisfactorily (n=1,438), yielding 16 designs. To increase design diversity, we added three designs where both trimers folded while we relaxed shape complementarity to >= 0.6 and tightened BSA to >= 1,800 Å^2^. We additionally selected an equal number of designs that were created either using the *interface* or *interface + neighbors* residue selection protocols. For the final 65 high quality designs, we subjected each structurally threaded model to thorough refinement in Rosetta (see methods) and removed 26 designs based on deviating shape complementarity and solvation free energy density.

The resulting 39 T33-ml (<u>m</u>achine learning) designs constitute a completely different set of sequences compared to prior work, including those from the T33-fn designs above (Figure 3g). Despite less stringent sequence design filters, in most cases the T33-ml metrics fell in ranges between those calculated for prior success and those from the T33-fn set (which were more polar). In most cases, the number (or fraction) of amino acids mutated was an order of magnitude higher than in past designs (Figure 3g). Surprisingly, for calculated interface free energies, most designs had positive values, while negative energies indicate more favorable binding or subunit association (Figure S2d).

### Experimental characterization of T33-ml designs

As before, we appended a single His Tag to a surface accessible terminus and synthesized 38 bicistronic genes for expression in *E. coli* (one design failed during gene synthesis). After expression and IMAC purification, 35 out of 38 showed both components expressed, while both components were present in the soluble fraction in 24 cases. From these, both components co-eluted from IMAC (Figure S3) in 17 cases. These were subjected to SEC to test for intact assembly. In SEC, 4 designs out of 17 demonstrated an unambiguous peak at the expected elution volume for the full assembly (Figure 4). For the other 13, more complex elution patterns showed intermediate assemblies and individual trimeric species (or monomers), among minor populations of intact assemblies (Figure S3b,4). Further, SEC-SAXS validated these observations for the best behaving designs (Figure S5a). Investigation by TEM showed particles possessing the predicted size and features of complete assemblies for 6 of the designs, with 6 out of 38 representing an experimental success rate of 16%.

**Figure 4.**
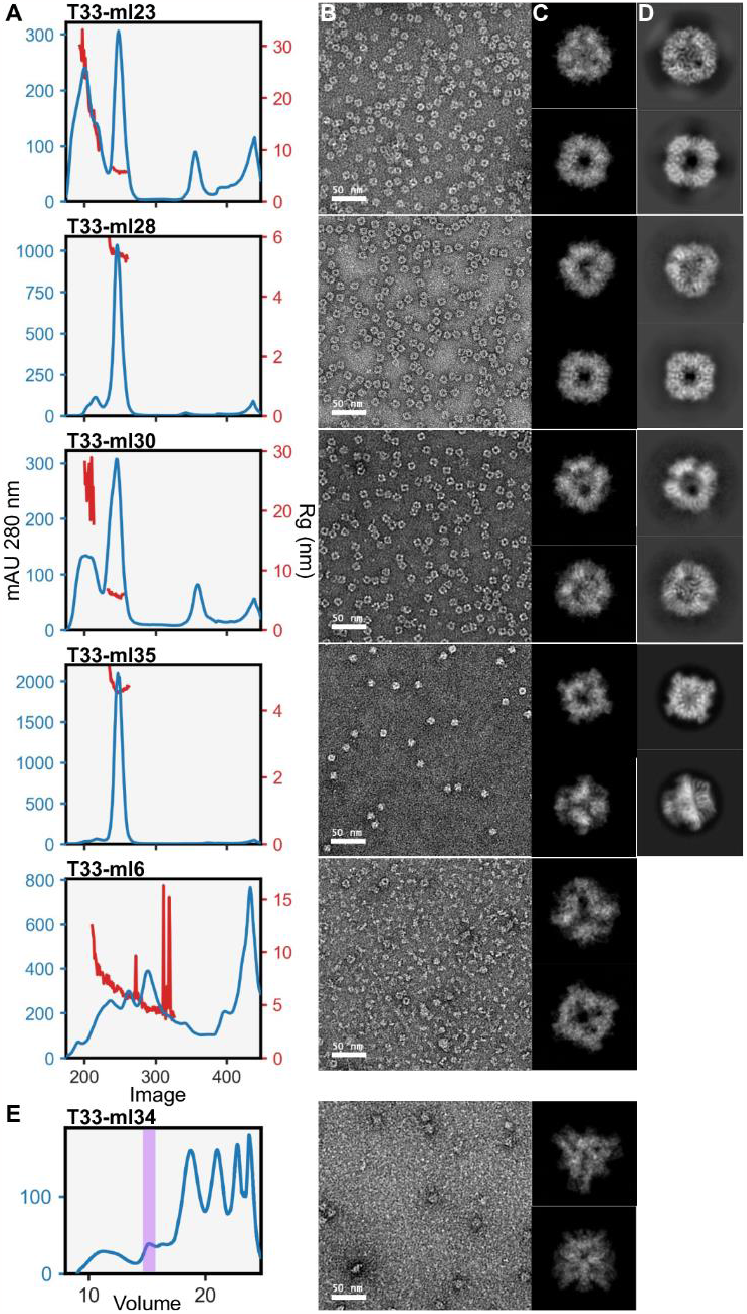
Biochemical characterization of six tetrahedral cages produced using machine learning protocols (T33-ml). **a-d)** Biophysical analysis on cages: SEC curves in blue, with SAXS measured radius of gyration (where available), in red (column a); negative stain micrographs (column b); 2D projections of the design (column c); and the 2D class where cryo-EM data collection was possible (column d). **e)** T33-ml34 demonstrates incomplete assembly by SEC, however, the IMAC elution visibly assembles under negative stain TEM. The proposed assembly elution volume is shaded in purple.

We undertook atomic structure determination by cryo-electron microscopy (cryo-EM) for four designs that behaved robustly in solution in order to assess their accuracy with respect to the intended models. For four designs, we acquired datasets where images demonstrated 2D classes matching model projections (Figure 4), and with sufficient orientational diversity to perform high-resolution 3D reconstructions. In all cases, the reconstructions were consistent with the expected positioning of trimers in the tetrahedral designs. Beginning with computationally designed models, we performed refinement into the resulting density maps (see Methods). We evaluated the LDDT of the resulting structures in the context of the design models as well as the RMSD over the entire assembly and over one A-B heteromer from the cage (Table 1). In figure 5, we present the various cryo-EM structures alongside their respective design models to emphasize the close agreement between design and experiment.

**Figure 5.**
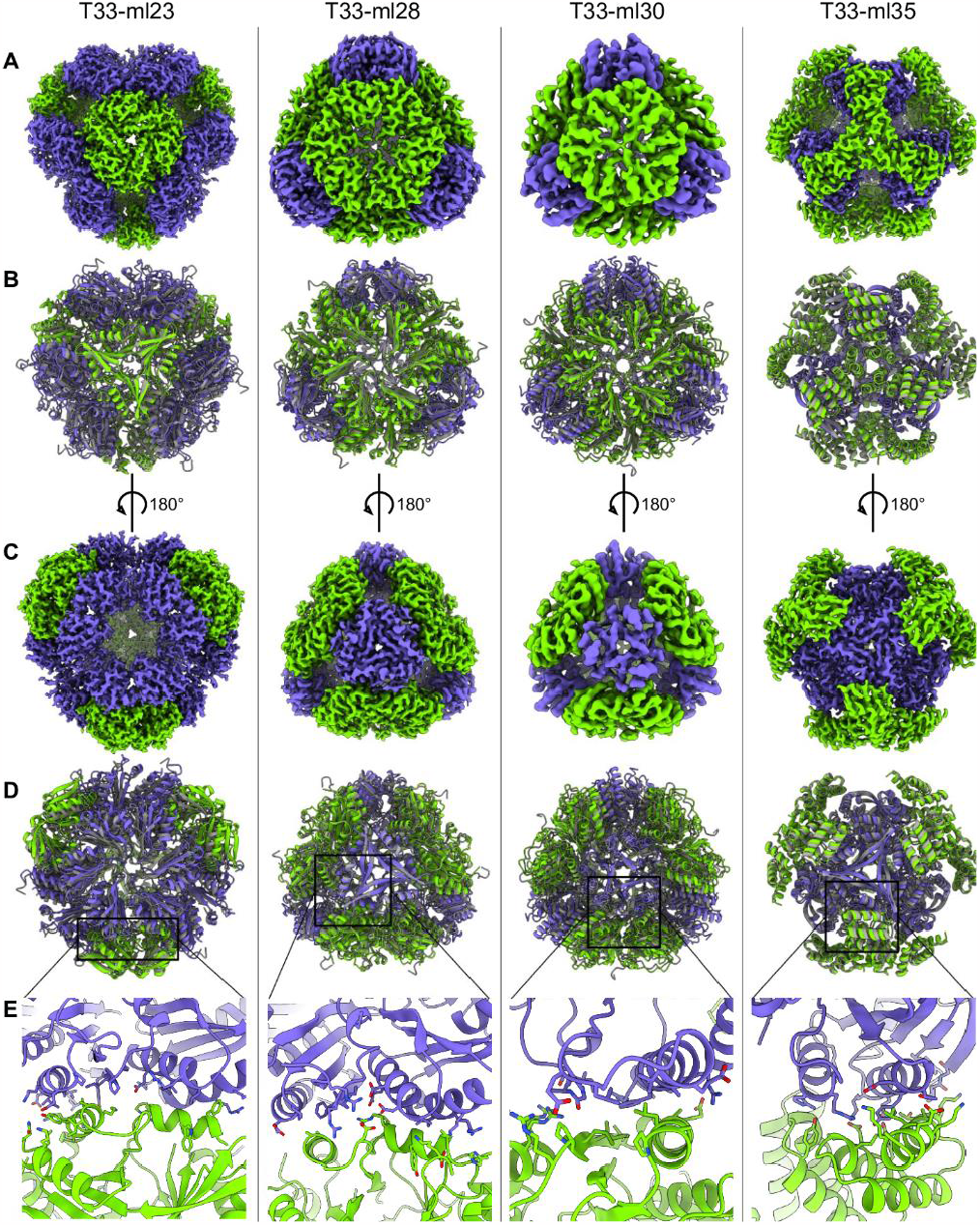
Structural characterization of assembling two-component cages by cryo-EM. **A-B)** The reconstructed cryo-EM map for the designs T33-ml23, T33-ml28, T33-ml30, and T33-ml35 are centered on the 3-fold axis of the first component (green) with the second component colored purple (row A); and superimposed on the design model represented in gray (row B). **C-D)** The reference frame is rotated 180° to center the cryo-EM map on the 3-fold axis of the second component (row C); while the superimposed design model in gray is also rotated 180°. **e)** A close-up of the residues for each structure that support formation of the *de novo* interface.

The best resolved design, T33-ml23, reached 2 Å resolution, revealing atomic details such as holes in aromatic side chains (Figure S5b,c). Although the full assembly deviates from the design by 2.4 Å RMSD, the cage demonstrated high stability and rigidity, which enabled outstanding imaging resolution. In contrast, for the worst resolved design, T33-ml30, refinement reached a resolution of 4.2 Å with a 3.6 Å RMSD for the whole assembly. Interestingly, in all cases minor rigid body adjustments occurred along the allowed degrees of freedom. Overall, this tended to result in slightly larger structures than designed (3% on average corresponding to an increased radius of gyration of 1.5 Å).

### T33-ml interface design analysis

Cryo-EM structures of our T33-ml cages showed overall deviations on the order of 1.5 Å from the designed models (Figure 6a). This motivated a comparison of calculated interface metrics for the experimentally realized structures vs. the designed models. Interestingly, we found that the calculated interface energies were more favorable for the actual experimental structures. We also noted that, even for the experimental structures, the values for calculated interface energies were mostly positive (tending to indicate dissociation). Among other limitations associated with calculated energy values, this serves as a cautionary reminder that energy calculations of this type do not adequately capture the importance of protein concentration in understanding association behaviors, especially in situations where many components come together in a complex.

**Figure 6.**
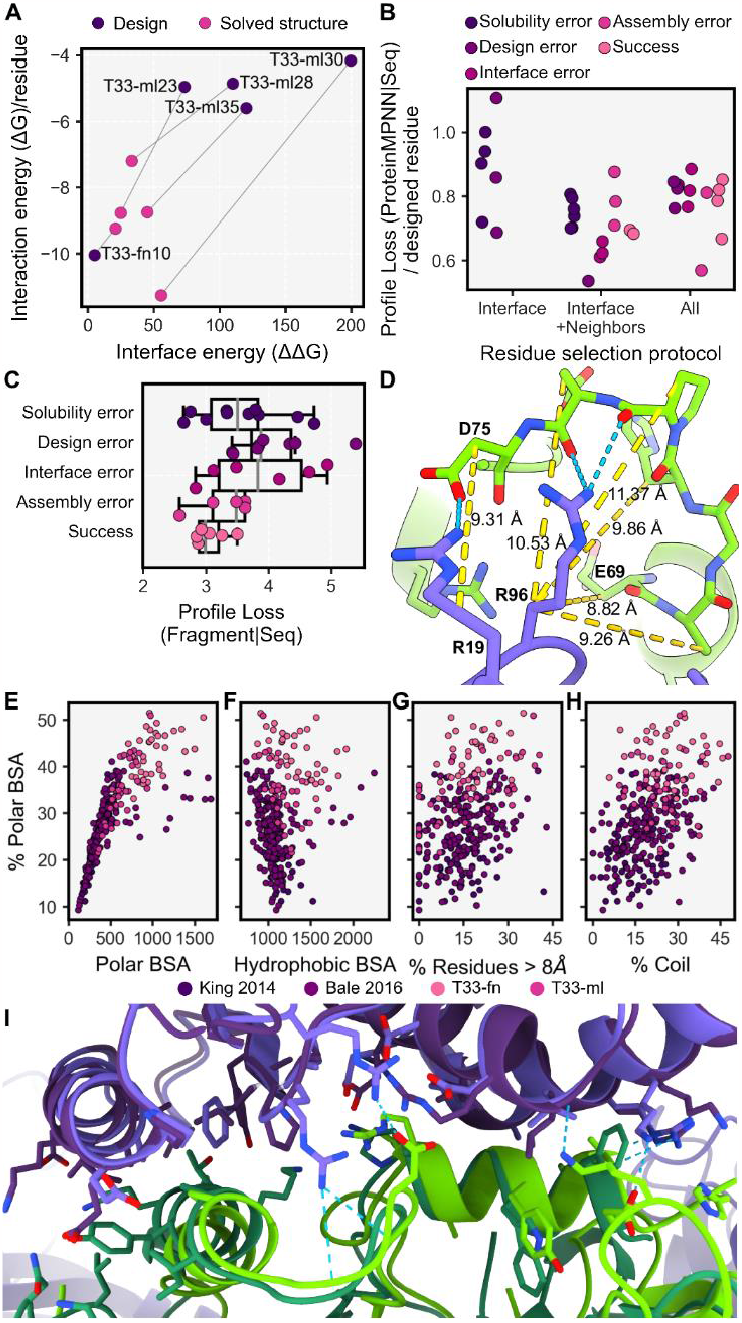
Understanding design outcomes. **a)** Rosetta interaction energy versus interface energy for the design models and solved structures presented in this work **b)** ProteinMPNN score per designed residue positions for biochemically characterized T33-ml designs grouped by design selection scheme and colored according to design outcome. **c)** Soluble T33-ml design outcomes correlate with fragment profile loss. Each point is grouped and colored according to design outcome. **d)** Interface comparison of T33-ml23 (dark purple/dark green) to T33-ml28 (purple/green). Cross-interface hydrogen bonds and salt bridges are indicated in blue for T33-ml28. **e-h)** For all two component cages from King *et al*. 2012, Bale *et al*. 2016, and this work, the fraction of polar buried surface area (BSA) as a function of polar BSA (panel e); hydrophobic BSA (panel f); percentage of loop/coil secondary structure (panel g); and percentage of residues in the interface with C-beta distances greater than 8Å (Coloring according to publication or T33-fn/T33-ml)(panel h); **i)** Example of a hydrogen bonding interaction between residues (A:19 and A:96 and B:69:75 in T33-ml28) with C-beta distances greater than 8Å.

As for T33-fn designs, we classified the T33-ml designs according to biochemical outcome (classifications discussed in T33-fn interface design analysis) to understand predictors of interface design success. We found that the ProteinMPNN score was not a strong predictor of design outcome (Figure 6b). When accounting only for soluble designs, the strongest predictor of biochemical outcome was a minimal fragment profile loss given the designed sequence (r^2^=-0.59, fig 6c). That is, successful designs tended to be those for which the amino acid sequence inferred by ProteinMPNN exhibited good agreement with the tabulated amino acid profiles for fragments at the interface. This finding highlights the benefit of using fragment-based sequence-structure information to guide pose selection and improve experimental outcomes for *de novo* designed interfaces.

### Variations and plasticity at the interfaces of T33-ml cages

From an analysis of our cage structures, we noted several cases where the observed structures revealed conformational adjustments that could be interpreted, *post-facto*, in atomic terms. For T33-ml30, a pair of helices interact across the interface, while peripheral strand and loop/coil sections participate in polar interactions. Though ProteinMPNN selected arginine residues to satisfy these hydrogen bond donor/acceptor pairs at the periphery, the structure reveals the polar groups were ultimately unsatisfied by interactions with other protein atoms. Instead, their burial was avoided through a rigid body rearrangement around the helical fragments (Figure 5e) which maintain the highest LDDT values for the pose (Figure S6). The T33-ml35 cage provides another example of this phenomenon. There, a rigid body shift compared to the design allows space for W90 and R94 of component B to fit in the interface, while fragment residues provide tight hydrophobic packing and an extensive network of hydrogen bonding (Figure 5e).

Two cages, T33-ml23 and T33-ml28, offer a particularly unique situation for comparison. The trimeric components that formed the basis for these two cages happen to be homologous and therefore structurally similar (Figure 6h). For both cases, one trimer is a tandem-BMC microcompartment protein while the other trimer is a protein from the CutA protein family; the trimers are also arranged similarly. Despite overall structural similarity (2.9 Å RMSD overall and 2.0 Å for the A-B subunit pair), the two cages have rather dissimilar sequences. Both cages were the result of design protocols where all residues were available for sequence redesign, which resulted in 58% overall sequence identity between the two cages, and only 44% over residues shared in the designed interfacial region. The interfaces also differ in overall character. For T33-ml23, the interface is more hydrophobic (75%), while the T33-ml28 interface (60% hydrophobic) presents over 10 polar interactions between the two trimeric components. This comparison highlights a high degree of sequence/structure degeneracy within the designed assembly space, an observation consistent with patterns of natural evolution in protein-protein interfaces ^50,51^.

### Increasing polar interactions in designed interfaces

We analyzed interface metrics for T33-fn, T33-ml, and all 2-component SCM designs that were docked in King *et al*. 2014 and Bale *et al*. 2016 ^43,44^. Unsurprisingly, we found that the amount of polar BSA positively correlates with the fraction of polar BSA (r^2^=0.783), but hydrophobic BSA was independent of the fraction of polar BSA (r^2^=0.095) (Figure 6e,f). For the surveyed designs, these relationships indicate that increased polar contributions are primarily obtained by increasing the total BSA rather than replacing hydrophobic atoms below some necessary minimum.

We next examined conformational features to understand how designs managed to bury polar atoms at their interfaces. The fraction of polar BSA positively correlated with the percent of interface residues whose C-beta atoms are separated by greater than 8 Å (Figure 6g). In those cases, polar interactions come from side-chain to side-chain contacts (enriched at the increasingly distant side-chain positions, Figure S7a), or through side-chain to backbone polar interactions, which can include C-beta distances up to 12 Å (Figure 6d). We next analyzed the extent of loop/coil segments in designs, as they make up a significant proportion of interface secondary structure in natural complexes ^52^ and inherently have more backbone polar atoms available for hydrogen bonding. The percent of interface loop/coil segments also positively correlated with the fraction of polar BSA (Figure 6h), while no correlation was observed with the percent of interface residues separated by greater than 8 Å (Figure S7b). The ProteinMPNN design approach is notable for its ability to accommodate backbone variation or uncertainty, and this appears to be an advantage for generating designs where more polar interactions are possible between loop/coil regions and their side chains. These observations and trends should guide future studies.

### Structural analysis of partial assembly states

During processing of cryo-EM data for the T33-ml23 and T33-ml35 cages, ∼20% of particles were classified in conformations representing partial assemblies along the route to a full A_12_B_12_ cage. We performed 3D reconstruction using C1 symmetry and built models into 3.9 Å and 4.4 Å density maps for T33-ml23 and T33-ml35 respectively (Figure 7a,b). For each, the partial assembly structures were very similar to the intact cage, but with a single trimer absent from the reconstruction (Table 2).

**Table 2.** Comparison of intermediate assemblies to complete assembly structures.

| Design Name | Assembly RMSD (Å) | ASU RMSD (Å) | Assembly LDDT |
| --- | --- | --- | --- |
| T33-ml23<br>A<sub>12</sub>B<sub>9</sub> | 1.27 | 0.93 | 0.76 |
| T33-ml35A<sub>9</sub>B<sub>1</sub><br>2 | 1.92 | 1.08 | 0.64 |

ASU - Asymmetric unit, RMSD - root mean squared deviation, LDDT - local distance difference test

**Figure 7.**
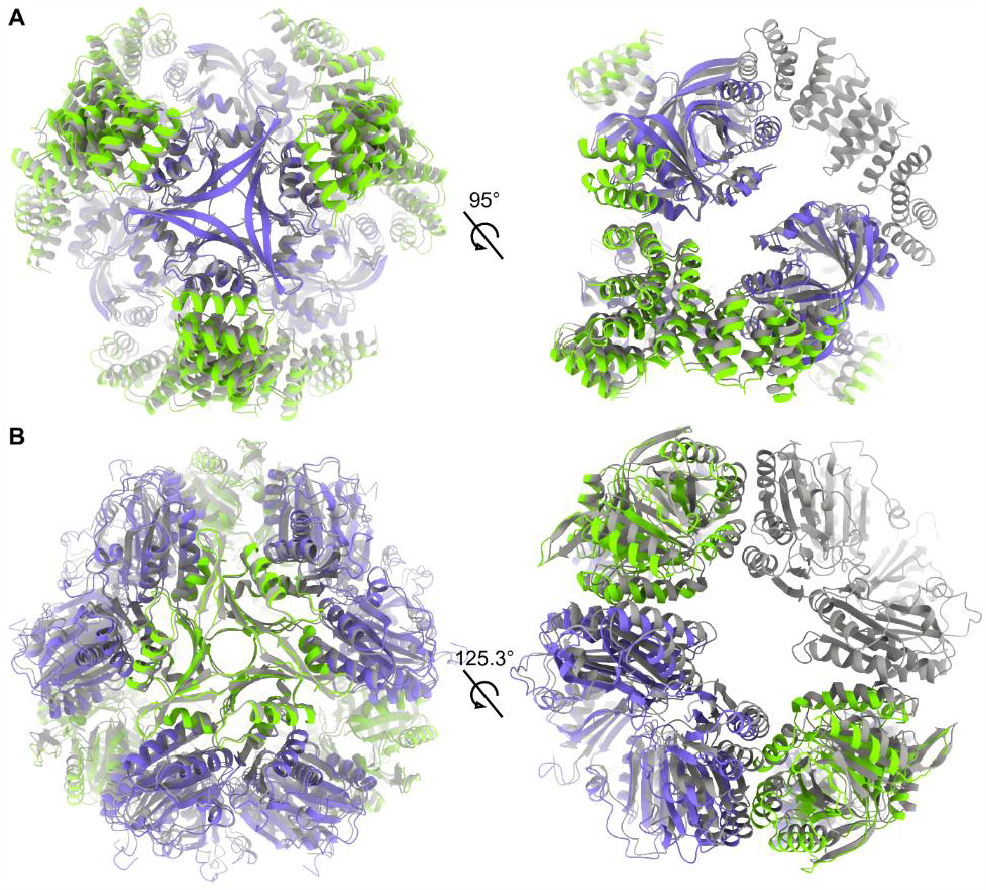
Assembly intermediates and structural comparisons to full assemblies. **a)** T33-ml35 A_9_B_12_ (green/purple) model superimposed on the A_12_B_12_ assembly (gray). A side view shows B components from A_9_B_12_ protrude into the site of the missing A_3_ trimer. **b)** T33-ml23 A_12_B_9_ (green/purple) model superimposed on the A_12_B_12_ assembly (gray). A side view shows A components from the A_12_B_9_ intermediate relaxing away from the missing B_3_ trimer.

A notable difference in the partial assemblies was the distance between equivalent backbone atoms across the missing trimer. For both intermediate structures, this distance increased by 1.0 Å compared to the full assembly. We take these displacements to reflect the magnitudes of the atomic rearrangements that occur during the stepwise or hierarchical assembly processes. We note that in T33-ml23, the missing component is a BMC trimer. Several studies have highlighted the flexibility of BMC proteins ^53,54^, and the observation of structural variation in partially assembled forms could reflect underlying flexibility of the components.

At present, it is not possible to ascribe an underlying cause to the formation of partial structures with certainty. Such cases could arise when minor design defects (*e*.*g*. slight deviations from ideal angles between components) propagate, making addition of the last puzzle piece dependent on substantial atomic movements (Figure 1c). Alternatively, partial assembly forms could reflect kinetic traps, as has been discussed in the context of viral capsid assemblies ^55^. Finally, it is notable that cages built from two (or more) distinct subunit types bring up the likelihood that one assembly component could be depleted before the other during assembly. To this point, it is notable that many viral capsids that assemble from multiple subunit types employ a proteolytic mechanism to produce the distinct subunit types from a longer polyprotein, which assures equal abundance of components^56^.

## Conclusions

The present study demonstrates the successful design and structural validation of a new suite of nanoscale symmetric protein cages using protocols that prioritize fragment-based sequence-structure relationships at *de novo* interfaces. These results elaborate on prior demonstrations of symmetric cages ^4,28,29,40,43,44^ and fragment-based design ^57,58^ by utilizing higher order positional relationships present in both fragments and graph-based neural networks to capture native protein properties.

For five of the new cages, we were able to determine atomic structures by X-ray crystallography or cryo-EM. Analysis of these cases generally confirmed the formation of intended tertiary motifs. Though non-helical motifs were employed in some cases, the designed interfaces were rich in alpha helical interactions overall. Differences between designed models and observed structures were generally small, but sometimes consequential in leading to unexpected, favorable atomic interactions.

From our studies, a comparison of different computational protocols – knowledge-based vs machine-learning – is complicated by the introduction of multiple different heuristic choices. Nonetheless, the high success rate (approximately 16%) that we achieved using the latest graph neural network approaches is notable. Introducing complex, native-like, structural features by intentional design is a major computational challenge, and this is particularly true with regard to polar interactions such as hydrogen bonds, which rely on greater atomic precision than hydrophobic interactions. Our successful results, as well as contemporary reports ^59^, indicate that machine learning methods are favorably suited for such complex tasks. Also notable for our design work was the allowance in machine learning for considerable backbone variation (*e*.*g*. by networks trained with backbone noise). In our study, this manifested in cases where important interfacial interactions were made between side chains in loop regions, which, in the absence of backbone variations in modeling, would not have been possible; *i*.*e*. they would not have been predicted using more conventional design methods based on fixed backbones. The consideration of backbone variation – including in insertions and deletions – is an area of key importance in ongoing work on antibody design ^60,61^. The machine learning approach also allowed a great deal of sequence novelty (with improved success rate) when the design protocol was not confined to the new interfacial regions.

As a last point of interest, our studies also illuminated the structures of partially assembled cages, which are likely representative of intermediates in the multi-step assembly process for cage formation. Indeed, partial structures appeared to dominate in a number of our failed designs. Understanding partial structures and the events leading to them could be impactful in improving design successes in future work.

## Acknowledgements

This work was supported by NIH grant R01GM129854 (TOY) and USPHS National Research Service Award T32GM008496 (KM). Additional resources for sample preparation and electron microscopy screening were supported by grant DE-FC02-02ER63421 from the Department of Energy Office of Science.

We thank Duilio Cascio for administration and support of structural techniques performed at UCLA, Genesis Falcon for assistance with crystallography, and Tom Holton and Alex Lisker for computing support. We thank Justin Miller for helpful discussions regarding biochemical investigation. We thank David Strugatsky for cryo-EM data collection acquired at the Electron Imaging Center for Nanomachines at the California NanoSystems Institute at the University of California, Los Angeles. We thank Patrick Mitchell for support in data collection and computation at Stanford-SLAC Cryo-EM Center (S2C2) under User Proposal CA151. We thank Omar Davulcu for support in data collection and computation at the Pacific Northwest Center for Cryo-EM (PNCC) under User Proposal 160163.

Some of this work was performed at the Stanford-SLAC Cryo-EM Center (S2C2), which is supported by the National Institutes of Health (NIH) Common Fund Transformative High-Resolution Cryo-Electron Microscopy program (U24 GM129541). The authors would like to thank Lisa Dunn and Patrick Michell for invaluable support and assistance with research performed at S2C2. A portion of this research was supported by NIH grant U24GM129547 and performed at the PNCC at OHSU and accessed through EMSL (grid.436923.9), a Department of Energy (DOE) Office of Science User Facility sponsored by the Office of Biological and Environmental Research.

We thank Tsutomu Matsui for critical assistance with SAXS data collection at the Stanford Synchrotron Radiation Lightsource (SSRL). Use of SSRL, SLAC National Accelerator Laboratory, is supported by the DOE, Office of Science, Office of Basic Energy Sciences under Contract No. DE-AC02-76SF00515. The SSRL Structural Molecular Biology Program is supported by the DOE Office of Biological and Environmental Research, and by the NIH, National Institute of General Medical Sciences (NIGMS) (P30GM133894). The Pilatus detector at beamline 4-2 at SSRL was funded under NIH Grant S10OD021512.

The contents of this publication are solely the responsibility of the authors and do not necessarily represent the official views of NIGMS or NIH.

## Author contributions

Conceptualization, KM and TOY; Methodology, KM, RCG, RA, MS, and TOY; Software, KM; Formal Analysis, KM; Investigation, KM, RCG, RA, MS, MA, TS, and CS; Resources, TOY; Data Curation, KM, RCG, and MS;Writing - Original Draft, KM and TOY; Writing - Review & Editing, KM and TOY; Visualization, KM and RCG; Supervision, KM and TOY; Funding Acquisition, TOY.

## Code availability

All code is freely available and hosted at https://github.com/kylemeador/symdesign. To perform design as described in this publication, a Google colab document is available at https://bit.ly/symdesign-colab.

## Declaration of interest

KM, RCG, MA, and TOY are inventors on a patent application by UCLA on materials described.

## Methods

All descriptions of computational processes are implemented in the symdesign repository https://github.com/kylemeador/symdesign. To perform design as described in this publication, a Google colab document is available at https://bit.ly/symdesign-colab. Where applicable, functions or scripts that are indicated to perform the described procedures are indicated in italic and relative to the repository’s main directory, i.e. *path/to/symdesign*. Flags and arguments for program operation are indicated in courier font.

### Structural preprocessing and docking

#### Docking round 1 inputs

Each of the round 1 trimers (see Supplement) was retrieved from the Protein Data Bank (PDB) in July 2020 using the following filters: global symmetry symbol equal to C3, X-ray diffraction dataset with resolution <= 2.5 Å, helical content > 30%, and *E. coli* as the protein expression organism. Entries with more than one protein entity or that contained nucleic acid entities were removed. Entries with a rcsb_polymer_entity_annotation.type containing the words OPM, mpstruc, MemProtMD, PDBTM, or MEMBRANE PROTEIN were removed. All structures were clustered using the 70% sequence identity threshold available from the PDB. The clustered representatives were then cross referenced against QSBio to select only high or very high confidence biological assembly predictions ^62^, and the corresponding biological assembly was downloaded. 84 candidate oligomeric building blocks were selected for docking trials. Before docking, a monomer from the assembly was chosen and subjected to symmetric refinement into the REF2015 score function ^37^ using the FastRelax mover and the suggested flags for pareto optimal refinement ^63^.

#### Docking round 2 inputs

Each of the round 2 trimers (see Supplement) was retrieved from the PDB in March 2023 using the following filters: global symmetry symbol equal to C3, but not symmetry type dihedral, X-ray diffraction dataset with resolution <= 3 Å, *E. coli* as the protein expression organism, and 80 <= number of residues <= 300. Entries with more than one protein entity or that contained nucleic acid entities were removed. Entries with a rcsb_polymer_entity_annotation.type containing the words OPM, mpstruc, MemProtMD, PDBTM, or MEMBRANE PROTEIN were removed. Additionally, if the entry title contained the keywords tail, fibre, shaft, head, spike, glycoprotein, ectodomain, or “receptor binding protein”, it was removed. All entries that satisfied these selection criteria were input into the thermophilic prioritization, assembly confirmation, and sequence clustering protocol as described subsequently.

First, the list of all organism taxonomy IDs were collected from ThermoBase_ver_1.0_2022 ^64^. These taxonomic ids were used to filter the selection criteria described above for only entries in which the source organism is one of the thermophilic ids. Each matching thermophilic entry ID was then subjected to the 30% sequence clustering service provided at the PDB. The sequence cluster groups returned entry ID’s sorted according to resolution. Each subsequent ID was iteratively tested for a high or very high confidence biological assembly annotation from QSbio ^62^. If a matching biological assembly entry wasn’t located within the cluster group in QSBio, the PDB was iteratively queried until an assembly with both “author_defined_assembly” and “author_and_software_defined_assembly” annotations was identified. This procedure was repeated in order of decreasing resolution until a match was found or all cluster group members were exhausted. For each cluster group where a confirmed assembly was located, the group was removed from further selection and the entry identifier was saved.

After selection of proteins from thermophilic taxonomic ids proteins, all remaining entry IDs from the initial selection criteria that weren’t found via thermophilic prioritization were again clustered according to 30% sequence identity from the PDB. Again, these non-thermophilic sequence clusters were subjected to the resolution sorting and assembly confirmation procedure in the prior paragraph until all 30% clustered groups were either represented by a confirmed assembly or discarded when all members were exhausted with no confirmed assembly representative. Every entry that passed thermophilic prioritization, assembly confirmation, and sequence clustering had the corresponding biological assembly downloaded for docking.

Building blocks may be retrieved using this procedure from the PDB API using the script present at:*symdesign/tools/retrieve_oligomers*.*py*

#### Modeling missing density

Each trimeric PDB entry was preprocessed to model missing density in internal loops and termini using AlphaFold. The reference sequence associated with the PDB entry was queried using hhblits (see Running hhblits) and the resulting multiple sequence alignment was parsed into a format corresponding to AlphaFold msa feature arrays. All subsequent inference occurred as described in AlphaFoldInitialGuess inference using the reference sequence as the sequence input and the biological assembly coordinates as coordinate input. The only exception was that the AlphaFold msa features processed here were provided to the AlphaFold feature dictionary instead of using a blank feature dictionary.

Of the AlphaFold predicted structures, the structure with the lowest RMSD to the input asymmetric unit (measured over all originally present C-beta coords from one protomer of the trimer) was structurally aligned to the asymmetric unit and selected as the disorder modeled trimeric representative. Each trimeric assembly was examined by eye for successful recapitulation of the input biological assemblies by AlphaFold. If the extent of modeled disorder was minimal, *i*.*e*. only few residues at the termini were added, the original trimeric biological assembly was used in place of the AlphaFold model. In rare cases, there were larger deviations present when comparing the biological assembly to the AlphaFold model by eye which may have indicated erroneous alignment of the prediction to the biological assembly given alternatively modeled residues or a lack of evolutionary evidence for the trimer resulting in weak predictions around the oligomeric interface. In all these cases the biological assembly was chosen if there was no unmodeled density, otherwise the trimer was discarded from consideration. Finally, all trimers were visually examined for large protruding features which caused prolonged instead of compact structures. These were additionally removed.

Whether a building block was provided in a file or retrieved from the PDB, the flag `--loop-model-input` will specify loop modeling should occur while performing preprocessing for the `nanohedra` module.

#### Generation of fragment observations

Fragment observations are identified as in Laniado *et al*. 2021 ^5^. Observation proceeds by identifying interface residues for the pose, examining each residue for surface accessibility, and then matching surface exposed residues to a particular fragment type, where the fragment type is determined by structural match between neighboring residues of a specified interval (here +/-2 residues) from the residue of interest (for a total of 5 residues) and corresponding fragments of protein structure from the fragment database (totaling 5 residues in length). The fragment type is similar to a secondary structure classification, but utilizes neighboring atoms for classification. Next, those surface accessible interface residues that match a fragment type are examined for potential fragment interaction by querying the identified fragment type against all unique fragment clusters from all fragment types. Fragment clusters constitute a spatially distinct observation of one fragment type with another and are clustered into dense, orientationally dependent groups by RMSD measurements. Fragments that do not clash with backbone or C-beta atoms constitute “ghost fragments” and represent each of the possible orientations between two fragment types that can potentially interact for the identified residues. Finally, ghost fragments are compared against another group of residues to match the other residue’s fragment type to the ghost fragment, fragment type. A match is identified if the RMSD between the other fragment and the ghost fragment is below a threshold, in this case 1 Å. For the case of interface residue fragment observations, the grouping of other residues belong to the surface of another oligomer, which may be of the same or a different entity type.

#### Nanohedra docking

All docking procedures can be performed using the `nanohedra` module from the provided git repository.

Nanohedra fragment-based docking routines were used as implemented in Laniado et al. 2021 ^5^. For T33-fn designs, docking was performed with the following parameters `--minimum-matched` 3 `--initial-z-value` 1 `--match-value` 0.5. In lieu of providing a desired fragment type for docking search, a modification was made to automatically calculate the majority secondary structure type of all surface exposed residues. This secondary structure type was used as the source of initial fragment overlap searching.

For T33-ml docking, the following modifications were used. Input building blocks had their termini trimmed back to remove extended loop/coil segments that mainly arise as the result of preprocessing with AlphaFold. This trimming occurs through the flag `--trim-termini`. Fragment potentials, *i*.*e*. ghost fragments, were subjected to an additional search constraint that they must also initially match with another ghost fragment from the same protein component. We refer to this search technique and implement its usage through the flag `-- continuous-ghosts` due to the use of multiple overlapping ghost fragments that overlap for a continuous region of the identified fragment potential. For instance, the residue i and i+4 of component 1 have continuous ghosts if their ghost fragments occupy the same location. Such a scenario is more likely if they are members of the same secondary structure, and if that secondary structure’s potential contacts consist of regularly spaced contacts.

#### Transformational clustering, fine grained search of docked space

As our ideally docked protein-protein interfaces contain multiple fragment observations, the pose created by ideal overlap of any single fragment observation has the effect of creating multiple redundant docked observations with subtle variation in transformational parameters. In docking, typically an advantage of redundant observations can be realized through clustering. By grouping each similar pose into an ensemble of potential positions, a global analysis of ensemble density revealed positions with the highest potential. To both prioritize fine grained search within clustered ensembles and reduce the roughness produced by discrete sampling of the available degrees of freedom, we implemented a grid optimization procedure into the final stage of docking. Importantly, optimization proceeds within the available degrees of freedom (DOF), so each available transformational DOF is placed on a grid and combinatorially sampled. After any single round of optimization, those positions with the highest scores are selected and a new grid is sampled from the DOF until the optimization target function is achieved. In this work, we chose Nanohedra score as our optimization target function and sampled until the change in score from one round of optimization to the next fell below %5 improvement or sampling new grid positions fell below the resolution of adequately describing a unique transformation. Unique transformations were binned into a six dimensional transformation hash, consisting of three translations and three Euler angles, to describe the unique parameters that specify a rigid body transformation between two bodies.

## Metric calculation

### Residue types

Given that a docked pose will have new amino acids specified at interacting residue positions, *interface residues* were defined as residues based on C-beta to C-beta atom distances. For docking measurements, the distance to identify residues was 9 Å, while for design, 8 Å was used. For all interfacial C-beta contact search procedures, residues are queried from a heterotypic globular component. This criterion also includes residues from a symmetrically related, *i*.*e*. non-self oligomer, of the same protein entity. For measurements on designed structures, *interface residues* were identified by their contribution of BSA to the interface. In all cases where the residue is modeled with the amino acid glycine, the C-alpha atom is used to measure C-beta distances. *Neighbor residues* are defined as residues with a C-beta atom within 8 Å of another residue’s C-beta atom. *Interface fragment residues* are defined as residue positions containing a fragment observation as identified by a fragment potential search. The interface *core, rim*, and *support* residues are defined according to ^31^.

### Position specific profile calculation

Each profile constitutes a per-residue amino acid frequency distribution, which is tabulated over all positions of the pose in question. All distributions are limited to the 20 canonical amino acids, and positions where no information is present are discarded.

#### Fragment profile

Every fragment observation is an association between a tertiary motif observed during modeling and a database of structurally clustered fragment observations. To represent the sequence preferences associated with a single fragment cluster, the amino acids observed from each member of the cluster are tallied to create a cumulative amino acid frequency distribution for each residue position in the fragment cluster. The resulting set of distributions represents the amino acid probabilities associating observed sequence preferences with the structural motif.

When fragment observations are identified for a pose, multiple fragments may be present at each residue. To capture the full sequence-structure probability from multiple fragment observations, each observation references the respective amino acid distribution at the corresponding structural position from the cluster. After collecting all participating distributions, they are combined to reflect the contribution of each fragment observation to the total fragment potential. This is accomplished through scaling every fragment observation distribution by both a match score and an interaction weight to enable summation to a single distribution.

For the match score, m, the structural match of the pose fragment observation to the representative fragment from the respective cluster is calculated according to equation 1.

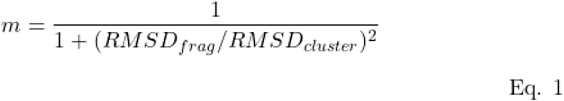

Where RMSD_frag_ is the RMSD between the fragment observation in the pose and the RMSD of the representative fragment from the cluster, and RMSD_cluster_ is the average RMSD measured for all member fragments which belong to the cluster.

For the interaction weight, t – the interaction importance – each structurally aligned residue in a fragment cluster is measured for atomic contacts with the opposite fragment pair. In this measurement, the number of times that side-chain atoms from the residue of interest interact with atoms from the opposite fragment is normalized by the number of side-chain atoms in the residue. This normalization creates a relative interaction weight for this residue in this member fragment from the cluster. Finally, for every member of the cluster, the mean interaction weight of the same residue position is taken as the residue specific fragment interaction weight, t_r_. Calculation of t_r_ is defined by equation 2.

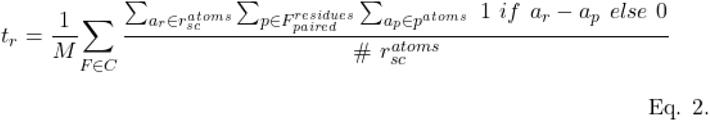

The variable f is the fragment observation from the set of all fragment observations from the fragment cluster, C, whereas p is a residue from the set of residues belonging to the paired fragment structure, F^residues^_paired_. The variable a_r_ is an atom from the side-chain (sc) atoms of residue r, a_p_ is an atom from residue p, and M is the number of members in the fragment cluster, C.

Equation 3 defines the process for creating an amino acid distribution for a single residue.

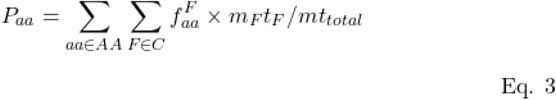

The set AA is the set of all amino acids and aa is a single amino acid. F represents a single observation from the set of all fragment observations, C. f ^F^_aa_ is the frequency of amino acid aa from fragment observation F. The quantity m_F_t_F_ is the match to the fragment cluster from observation i multiplied by the interaction potential for the particular fragment index associated with the fragment cluster from observation F. Finally, mt_total_ is the sum of all m_F_t_F_ observations for the residue and creates the denominator for the scaling factor for each F observation by its overall contribution importance.

The total fragment profile for a pose reflects all amino acid frequency distributions, P_aa_, calculated for each residue in the pose with fragment observations.

#### Evolutionary profile

The hidden markov model written to the .hhm file extension was parsed and the model values for each amino acid at each residue were converted to amino acid frequencies. These values constituted the per-residue amino acid frequencies for evolutionary profile based calculations. All positions of an evolutionary profile that were generated based on a reference sequence, but were used in subsequent structural analysis, were removed from the structure representation to keep the index of the reference residue aligned with the structure. This is important in adjusting disordered internal or terminal residues to have the proper profile alignment.

#### Tertiary profile

The tertiary profile is the per-residue amino acid distribution calculated from a combination of the fragment profile and the evolutionary profile. The tertiary profile assumes the amino acid frequencies of the evolutionary profile for residue positions where there are no fragment observations, and therefore, no fragment profile. For positions where there is fragment profile information, the resulting distribution is weighted by the fragment profile distribution with a maximum weight of alpha (here 0.5) with the inverse, (1 - alpha), contributed by the evolutionary profile. If the fragment observations are deemed weak, according to the fragment match, m and the fragment interaction, t, then alpha is reduced by a modifier which reflects the weakness. Such a modifier reduces alpha, and thus the fragment profile’s contribution to the tertiary profile. The m and t value modifiers are calculated by comparison to the fragment database and cluster references, respectively. If the values fail to meet quality thresholds, alpha is modified by the proportion of the discrepancy.

During Rosetta design protocols, the tertiary profile specified which amino acids were available for sampling at each position. Those amino acids that had a frequency greater than 0, through the flag –use_occurence_data, were available for design. For the HBNet protocol, those amino acids that had a frequency greater than 0 for the fragment profile were utilized for the design of fragment residues.

### Nanohedra score

Nanohedra score was calculated according to the procedure in Laniado *et al*. 2021 ^5^ except the score was only tabulated for central fragment residues, not every fragment residue. For normalization, the Nanohedra score was divided by the number of fragment residues. This gives a maximum normalized value of 2.

### Interface energy

All default energy terms from the REF2015 score function were calculated for each residue and residue neighbor in the interface and summed to yield various solvation energy measurement states. The interface energy complex is calculated on the full complex, while the bound version is calculated on the bound confirmation in the oligomeric state, and the unbound version is calculated on the minimized confirmation in the oligomeric state. To calculate the interface energy, the unbound is subtracted from the complex (complex - unbound), while to calculate the bound configuration energy, the bound is subtracted from the complex (complex-bound).

### Interface solvation energy

The energy terms, `lk_ball_wtd` and `fa_solv` were calculated for each residue and residue neighbor in the interface and summed to yield various solvation energy measurement states. The solvation energy complex is calculated on the full complex, while the bound version is calculated on the bound confirmation in the oligomeric state, and the unbound version is calculated on the minimized confirmation in the oligomeric state. To calculate the Interface solvation energy, the complex is subtracted from the unbound (unbound - complex), while to calculate the bound configuration energy, the bound is subtracted from the unbound (unbound - bound).

### Interface bound configuration energy

To calculate the energy required to attain the atomic conformations utilized during interface complexation from the uncomplexed state (not including the process of complexation with the interface partner), the energy at each residue was captured for both states, the complexed conformation form of the oligomer and the oligomer after four rounds of atomic packing and minimization. Next, the energy difference between the individual residues in the bound conformations and the unbound conformations were taken, with positive values indicating energy is needed to assume the bound conformation.

### BSA calculation

The atoms participating in the interface are measured for solvent accessible surface area (SASA) in both the complexed state and the uncomplexed state (no repacking). Next, the total SASA in the uncomplex state was subtracted from the SASA in the complex state to find the difference. For segregation of SASA by atomic polarity, only atoms that were deemed polar had SASA summed, while for hydrophobic BSA only non-polar atoms were summed. All calculations were performed using the program FreeSASA^65^.

### SS calculation

The program Stride ^66^ was used to calculate secondary structure for each residue. When secondary structure percentages were assigned to interface portions, the total number of interface residues was used as the denominator and the number of residues of a particular secondary structure type were the numerator.

### ProteinMPNN scores

ProteinMPNN score was calculated as in Dauparas et al. 2022 ^7^, where the score was the average over all residues in either the complexed state, *i*.*e*. bound interface, or the uncomplexed state, *i*.*e*. the interface was separated. For ProteinMPNN score for a particular subset of residues, only those residues were averaged. The total ProteinMPNN score reflects a total summation of the individual scores for all residues.

### Shape complementarity

Calculations were performed using the ShapeComplementarityFilter in Rosetta. During design calculations residues were selected based on 8 Å C-beta C-beta interface residue membership. For calculations performed during retrospective analysis, residues were included if they were interface residues according to BSA involvement.

### Buried unsatisfied hydrogen bonds

The number of hydrogen bonds participating in the complexed interface state and the uncomplexed interface state (no repacking) were summed. The number of unsatisfied hydrogen bonds in the uncomplexed state was subtracted from the number of unsatisfied hydrogen bonds in the complexed state to find the difference.

### Local distance difference test (LDDT)

The LDDT score ^67^ was utilized according to the original implementation to compare structures regardless of superposition technique. For cage assemblies, all reported values are the result of calculation of one chain that was perfectly symmetric.

### Root mean squared deviation (RMSD)

Calculation was performed using the Kabsch algorithm for finding an optimal overlap and calculating an RMSD.

### New hydrophobic collapse sites

The hydrophobic collapse index (HCI)^68^ was calculated with the following modifications. Instead of the amino acid types FILV being classified as collapsible residues, the amino acid types were expanded to include FMILYVW. Additionally, the HCI threshold was modified to 0.48 from the reported 0.43 to maintain consistent overlap with observations of collapse ^68^. The HCI was taken for both the designed sequence and the reference sequence, *i*.*e*. the sequence before sequence design occurred, and regions in the designed sequence that resulted in HCI larger than the HCI threshold, but were not larger than the HCI threshold for the reference sequence were determined to be new hydrophobic collapse sites.

### Interface composition similarity

The residue burial types in the designed interfaces were calculated according to classification as either core, rim, and support residues. To measure the similarity between designed interfaces and natural interfaces, the number of interface residues of each type were compared to the number of residues expected based on the size of the interface. To find the expected values of each residue burial type, lines of best fit reported in Levy 2010^31^, were used to calculate expectation, corresponding to: core = 0.01*BSA + 0.6, rim = 0.01*BSA - 2.5, and support = 0.006*BSA + 5. For the calculated BSA, the difference between the expected number of residues and actual number of residues in each classification, R, was found. Next the percentage of this difference from the expected value was subtracted from 1. After the expected difference is summed for each residue, the mean value was taken as the interface composition similarity as in equation 4. Values of 1 indicate exact similarity to the expected interface composition values and 0 indicates no similarity.

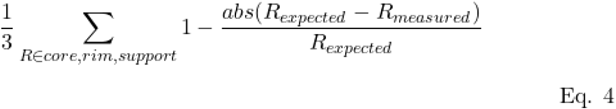

### Spike ratio

The spike ratio is defined here as 1 minus the ratio of how distant the centers of masses of the two trimeric components are from the center of mass of the full cage. In this comparison, the larger distance takes the denominator, so a ration of 1 indicates the two component are equidistant from the cage center, giving a spike ratio of 1 – 1 = 0, while a ratio of 0.5 would indicate one component is twice as far from the cage center as the other, giving a spike ratio of 1 – 0.5 = 0.5

## Pose selection

A weighting scheme was utilized to prioritize poses and designs based on evaluation of metrics calculated for each considered structure/sequence. Those poses or designs that demonstrated the largest weighted sum were selected, where the individual weighting terms are according to a provided weight and the normalized value of that pose/designs metric compared to all others considered.

### 590 candidate poses and prioritization of their designs for T33-fn characterized sequences

For selection of poses, the metrics, direction of prioritization, and weighting coefficient are as follows: shape complementarity of fragment residues, prioritize higher values, 0.3 weight; interface residue composition similarity, prioritize higher values, 0.4 weight; the percent of interface with fragment observations, prioritize higher values, 0.1 weight; and the percent of hydrophobic BSA, prioritize lower values, 0.2 weight.

For each set of designs from each filtered pose, the weighting scheme was applied similarly to select the best design, with the metrics, direction of prioritization, and weighting coefficient as follows: interface energy, prioritize lower values, 0.25 weight; interface bound configuration energy, prioritize lower values, 0.15 weight; atomic density of non-hydrogen interface atoms (following the local density metric ^39^), prioritize higher values,0.2 weight; the density of buried unsatisfied hydrogen bonds, prioritize lower values, 0.15 weight; shape complementarity of the interface, prioritize higher values, 0.1 weight; and the BSA to SASA ratio of the pose, prioritize lower values, 0.15 weight.

### 4,241 candidate poses for T33-ml

For T33-ml designs, 4,241 poses were investigated in depth. These were selected according to the following filters. The type of residue selection protocol was all residues, there were no new hydrophobic collapse sites, the fraction of residues that were different amino acids from the original sequence, *i*.*e*. mutated positions, were less than 55 %, the ratio of fragment observations at each residue site (multiple fragment ratio) was greater than 2.5, the number of fragment observations at the interface was greater than 20, the total ProteinMPNN score in the unbound state was less than 880, the ProteinMPNN score in the complex state over all designed residues was less than 1, the profile loss of the sequence given the evolutionary profile was less than 2.5 (average of all residue positions), the profile loss of the sequence given the fragment profile was less than 5 (average of all fragment residues), Finally, shape based features such as the spike ratio less than 0.5, and the distance between the cage center of mass and the minimal atomic distance for each component which is less than three times the distance for the component which is displaced further.

Finally, from the selected poses, one design was selected according to the lowest ProteinMPNN score and characterized through computational structure prediction and threading.

## Design protocols

### Symmetry

All design methodologies were undertaken in the presence of the entire symmetric system in question. For oligomeric state measurements, this was in the trimeric system, while for complex state measurements, either the entire tetrahedral complex was created, or, where applicable, the minimal contacting group of protein chains was constructed, which contained the minimal information necessary to fully model all possible interactions present in the complex. All modifications made to the sequence of oligomeric components were maintained at a single protomer representative (the captain) and propagated symmetrically to all identical, but spatially separate, symmetry mates.

### Running hhblits

Per-residue hidden markov models were created using amino acid sequences submitted to hhblits ^69^ against the UniRef30_2020_02 database (retrieved from https://gwdu111.gwdg.de/~compbiol/uniclust/2020_02)/ and run with the following parameters `-ohhm` *FILENAME*.*hhm* `-oa3m` *FILENAME*.*a3m* `-hide_cons -hide_pred-hide_dssp -E 1E-06`.

### iAlign clustering

Clustering according to interface alignment in iAlign ^70^ was performed. Where it was found that there were overlapping poses, the best pose was selected according to the weighting procedure described in 590 candidate poses and prioritization of their designs for T33-fn characterized sequences.

### Tertiary constrained FastDesign

Metropolis criteria Monte Carlo optimization was performed using the FastDesign mover in Rosetta. FastDesign is set up with a SequenceProfile TaskOperation, which limits the residues available for packing to those residues specified by the provided position-specific scoring matrix file ^71^. The profile of interest for this protocol was the tertiary profile. Additionally, design (defined as amino acid sampling) is only allowed at interface residue positions, while interface residue neighbors are allowed to pack (defined as sampling of residue conformations such as backbone and rotameric states). Five rounds of FastDesign sampling are performed in the REF2015 score function and the resulting asymmetric unit is written to a file.

The implementation of this protocol can be found at *dependencies/rosetta/interface_design/design_profile*.*xml*.

### Scouting FastDesign

Scouting constitutes a quick round of design where minimal residues are selected, few amino acid types are allowed for design, and all FastDesign is subjected to one cycle of design. The first set of residues designed constitutes the interface fragment residues and uses the FastDesign with protocol InterfaceDesign2019. The second set of residues set for design constitutes the remaining interface residues and uses the FastDesign protocol PolarDesign2019.

The implementation of this protocol can be found at *dependencies/rosetta/scout*.*xml*.

### FragHBNet

The combination of HBNet with fragment residue packing prioritizes extensive hydrogen bond networks for their ability to simultaneously support well packed, hydrophobic interactions. The method is a modification to the search procedure of the MCMC HBNet protocol (Maguire 2021). Importantly, the `start_selector` keyword in the HBNet Mover uses a `residue_selector` which includes all observed fragment residues identified for the pose. The residue positions that are available for HBNet inclusion include interface residues (of which fragment residues belong), and interface neighbors. During HBNet a search for interface residues utilizes the amino acid types available from the union of the fragment and evolutionary profiles, where a frequency above 0 (via the flag `--use_occurence_data true`) is included, while for interface neighbors, only the wild-type amino acid type is available. HBNet proceeds for 50,000 Monte Carlo runs and accepts all found networks less than -0.65 REU, containing three or more residues. After HBNet search, the top 250 networks are input to the MultiplePoseMover, which performs one cycle of FastDesign sampling (with REF2015 score function and PolarDesign2019 weights) at FragHBNet residues, defined as an interface residue, an identified hydrogen bond network residue, or a neighbor of a network residue. MCMC sampling at interface residues utilizes the intersection of amino acids available from the fragment and evolutionary profile (`--use_occurence_data true`), while those outside the interface are only allowed to utilize amino acids available from the evolutionary profile (`--use_occurence_data true`). After each candidate network is briefly designed, the entire group of candidate networks is ranked and the top N network candidates are selected. In this work, we used the top 20 network candidates.

Ranking proceeds for the highest N*3 (60 in this work) candidates measured according to FragHBNet residue shape complementarity and the lowest N*3 (60 in this work) measured heterotypic interface interaction energy per-FragHBNet residue. Finally, candidate poses that fall within both of these ranking criteria are ranked in descending order according to the number of residues included in the FragHBNet. The top N (20 in this work) candidates are subjected to deeper sampling using 5 rounds of constrained FastDesign with HBNet participating residues restricted to repacking with an additional AtomPairConstraintGenerator utilized with `sd`=0.4 to minimize undesired loss of the identified hydrogen bond networks. This constraint was set up to mimic the energetic contributions of hydrogen bonds in REF2015 to allow HBNet residues to flexibly readjust as the remainder of the interface was designed.

The scripts for these design protocols are found at: *dependencies/rosetta/hbnet_scout*.*xml, dependencies/bin/sort_hbnet_silent_file_results*.*sh*, nd*dependencies/rosetta/hbnet_design_profile*.*xml*.

### Reversion criteria

For each residue in a design, reversion utilized a fuzzy prioritization mechanism wherein mutations are made to the original amino acid and measured for their effects ^72^. The prioritization filter accepted mutations if they resulted in a decrease in shape complementarity less than 0.02 units and resulting in a negative value for calculated hydrogen bonding energy. Subsequently, all passing mutations were ranked according to highest shape complementarity, lowest hydrogen bonding energy, and lowest unsatisfied hydrogen bonds. The best scoring reversions were accepted followed by iterative testing of remaining reversions. Those remaining were combined with prior accepted reversions and again tested for their cumulative impact, accepting if the above filters passed.

The implementation of this protocol can be found at *dependencies/rosetta/optimize*.*xml*.

### ProteinMPNN

For sequence design, residues identified as designable by one of three protocols, all - every residue, interface - interface residues, or interface+neighbors - interface residues and their neighbors, were specified as positions available for ProteinMPNN sequence inference. All positions and coordinates were symmetrized including symmetrically tying positions and using the ProteinMPNN.tied_sample() method. In protocols where not all residues were used for designs, the remaining positions were input as their wild-type identities into the model and contributed to the encoding and decoding steps. The final sequence and probabilities were trimmed to only the minimal sequence representing the asymmetric unit. Resulting probabilities were transformed to represent an inference profile, representing the per-residue amino acid frequency distribution predicted as a result of inference.

For structure profile creation, symmetrized coordinates were used as inputs and the ProteinMPNN class was called. The resulting log probabilities were transformed to normal log probabilities to create the ProteinMPNN structure profile, *i*.*e*. the per-residue amino acid frequency distribution predicted for the protein coordinates alone.

### Rosetta refinement

To ensure that Rosetta energy calculations were carried out as accurately as possible, all designs were relaxed into Rosetta before metrics were acquired. Designs were input as an asymmetric unit, with symmetry set using symmetry definition files ^73^. Symmetric refinement was performed into the REF2015 score function ^37^ using the FastRelax mover. Additionally, the suggested flags for pareto-optimal refinement ^63^ were included for five rounds of refinement including the flags:

`-relax:ramp_constraints false`

`-no_optH false`

`-relax:coord_cst_stdev 0.5`

`-nblist_autoupdate true`

`-relax:bb_move false`

`-constrain_relax_to_start_coords`

`-use_input_sc`

`-relax:coord_constrain_sidechains`

`-flip_HNQ`

`-no_his_his_pairE`

The implementation of this protocol can be found at *dependencies/rosetta/refine*.*xml*.

### Threading of ProteinMPNN sequences to the designed structure

For structural metric measurement of ProteinMPNN designed sequences, the amino acid identities at each residue position were mutated to the designed ProteinMPNN sequence. Threaded designs were then refined in Rosetta following the Rosetta refinement procedure. For threading, however, refinement only used one cycle of refinement to sample the structural state.

The implementation of this protocol can be found at *dependencies/rosetta/refine*.*xml*.

### Refinement for structural analysis

For retrospective analysis of cage designs, all design models were subjected to an additional iteration of Rosetta refinement. In most cases, designs should be minimized after one round of Rosetta refinement. In T33-ml designs, however, a higher fraction of amino acids were mutated and thus backbone rearrangements were expected. As a consequence, this extra refinement ensured further convergence to minimum energies.

## Structure Prediction

### AlphaFoldInitialGuess inference

For all inference performed using the AlphaFoldInitialGuess model, the amino acid sequence was processed into arrays corresponding to AlphaFold sequence features, and structure coordinates were processed into arrays to be input as coordinate positions to the AlphaFoldInitialGuess class, which was derived from descriptions provided in Bennett et al. 2023 ^49^. AlphaFold msa features were provided empty to the AlphaFold feature dictionary as described in ^74^. All predictions were performed using AlphaFold multimer with the model parameters `multimer_v3`. Residues that were missing in their entirety or had side chain atoms missing had atomic coordinates initialized at the origin (i.e. 0,0,0). The structure coordinates were then input into the `prev_pos` feature to emulate a prior round of model inference, at which point AlphaFold was run having assumed these starting coordinates were a result of a prior round of prediction. Predictions preceded using for all five of the provided AlphaFold multimer model parameters unless satisfactory confidence metrics were achieved (pLDDT ≥ 85), at which point prediction was terminated. After model inference, the most confident model was refined into the Amber score function.

## Biochemical characterization

### Protein expression

Plasmids containing two genes of interest, but constituting a single design, were acquired from Twist Biosciences. Each plasmid contains a Kanamycin resistance gene for selection as well as a pET-Duet type interspatial expression cassette cloned between the genes of interest to enable bicistronic expression of the two proteins under the control of a LacO inducible T7 promoter. Plasmids were transformed into LOBSTR BL21(DE3)-RIL competent cells (Kerafast - EC1002) and always supplemented with 50 μg/ml of Kanamycin during growth. First, cells were grown to saturation overnight in 20 ml of PG media ^75^. The following day, 10 ml of saturated culture was used to inoculate 1 L ZYM-5052 cultures ^75^ which was left to grow for either ∼66 hours at 18°C, ∼42 hours at 25°C, or 16 hours at 37°C. Cells were harvested by centrifugation at 4,000 xg for 10-15 minutes and quickly moved to freezing conditions at approximately -80°C. Pellets were transferred to -20°C after 24 hours at -80°C.

### Immobilized metal affinity chromatography (IMAC)

Purification of proteins from cell pellets was performed with a 1:4 ratio of grams of pellet to milliliters of purification buffer (i.e. 10 g : 40 ml). Purification buffer contained 50 mM Tris-HCl, pH 7.5 at 4°C, 300 mM KCl, and 30 mM imidazole, while lysis buffer (utilized during cell lysis) was supplemented with 0.5 mM EDTA, 300 μg/ml lysozyme, 100 μg/ml benzonase nuclease, and one tablet of protease inhibitor cocktail EDTA-free (Thermo Scientific)/50 ml of cellular suspension. Pellets were resuspended in lysis buffer with end over end rotation for ∼30 minutes at 4°C until no visible chunks of cell pellet remained. Lysis of cellular resuspension proceeded with three passes of high pressure homogenization in a EmulsiFlex C3 homogenizer (Avestin) pressurized to > 15,000 psi. Lysate was then clarified using centrifugation at 20,000 xg for 30 minutes or 10,000 xg for 45 minutes at 4°C. The soluble supernatant fraction was collected and either incubated with 2 ml Ni-NTA resin/50 ml of soluble lysate (batch binding) or applied to a 5 ml HisTrap pre-equilibrated with purification buffer.

For batch binding, side over side rotation of the soluble lysate and Ni-NTA resin proceeded for 30 minutes at 4°C, after which the solution was allowed to stand for five minutes at 4°C, enough to separate the Ni-NTA resin and the supernatant by gravitational forces. Then, the supernatant was removed and the remaining Ni-NTA resin was collected and loaded into a filtered column where remaining soluble lysate was cleared from the Ni-NTA resin by gravity. The Ni-NTA column was washed by gravity flow with 10 column volumes (CV) of purification buffer supplemented to 60 mM imidazole (wash buffer). Finally, protein of interest was eluted from the resin with gravity flow using 9 CV of purification buffer supplemented with 250 mM imidazole (elution buffer).

For HisTrap binding, the soluble lysate was applied to the 5 ml column at 1 ml/min until all soluble lysate had passed through the column. Next, 10 CV of purification buffer were applied, followed by 2 CV of wash buffer and then a gradient from 60 mM imidazole to 300 mM imidazole (elution buffer concentration) was carried out for 3 CV. Finally, 10 CV of elution buffer was passed over the column.

Fractions corresponding to the lysate, the clarified lysate, the insoluble lysate, the unbound Ni-NTA supernatant (aka the flow through), the wash, and the elution were analyzed by SDS-PAGE for bands at the hypothesized size for the proteins of interest and classified according to one of insoluble, soluble, or co-eluting based on the presence of both bands.

### Size Exclusion Chromatography (SEC)

After IMAC purification, fractions containing proteins of interest were concentrated using Amicon Ultra 10 kDa MWCO concentrators until the sample reached approximately 500 μl volume or precipitation was visibly observed at which point concentration was immediately stopped. After concentration, the sample was clarified by centrifugation for 10 minutes at 16,900 xg to remove any precipitation from soluble protein. All chromatography was performed using the BioRad NGC HPLC system at 4°C. Analytical chromatography was performed using a 10/300 gl size column with either Superose6 Increase or Superdex200 Increase resin with flow at 0.3 ml/minute. For preparative scale chromatography, 16/60 scale columns with either Superose6 or Superdex200 residue was used while flowing at 0.9 ml/minute. Fractions corresponding to peaks in the chromatogram, observed by absorbance at 230/280 nm, were analyzed by SDS-PAGE for protein species of interest.

### CryoEM specimen preparation

Purified samples of the designed assemblies were removed directly from SEC fractions if the fractions were deemed sufficiently concentrated (>0.5 mg/ml) otherwise were concentrated to ∼1 mg/ml and stored at 4°C until specimen preparation. Upon freezing, QUANTIFOIL R 2/1 Cu 300 mesh grids (Cat # Q3100-CR1) were glow discharged for 30 seconds with a Pelco easiGlow glow discharger using 15 mA and a negative polarity. Immediately after glow discharge, grids were loaded into a Vitrobot Mark IV Thermo Fisher Scientific set to 100% humidity and blotted with 595 Filter Paper 55/20mm (Ted Pella Cat# 47000-100) using 1 total blot with a 0 second wait time, -4 blot force, 4 second blot time, and 0 second drain time and vitrified into liquid ethane. All specimens were prepared using a protective plastic facemask and a surgical mask to avoid contamination.

The sample for T33-ml30/T33-ml35 mixed were measured with absorbance of 0.583 as measured directly from SEC were frozen in SEC buffer consisting of 20 mM Tris-HCl pH 7.5 at 4°C, 150 mM KCl. The design T33-ml28 was diluted to 0.31 mg/ml using 20 mM Tris-HCl pH 8 at 214°C and 100 mM NaCl. The design T33-ml23 was diluted to 0.6 mg/ml using 20 mM Tris-HCl pH 8 at 21°C and 100 mM NaCl.

### Cryo-EM data acquisition and processing

Cryo-EM data were collected on Titan Krios cryo-electron microscopes equipped with a Gatan K3 Summit direct electron detector. Movies were recorded with SerialEM ^76^ at a nominal magnification of 81,000× (calibrated pixel size of 1.1 Å per pixel), 105,000× (calibrated pixel size of 0.86 Å per pixel) and 130,000x (calibrated pixel size of 0.65 Å per pixel), over a defocus range of −0.5 to −2.5 μm and a total dose of 40 e^−^/Å^2^.

Motion correction, CTF estimation, particle picking, particle extraction, 2D classification, and additional data processing were performed with cryoSPARC ^77^. An initial set of particles was automatically picked through the blob-picker method, extracted and 2D classified. Particles selected from 2D classes were used for ab initio reconstruction. This reconstruction was then used for the 3D refinements enforcing T symmetry. The 3D structure was used to generate 2D projections of the particles and then used to repick the particles from the images using a template picker. The re-picked particles were extracted from the micrographs, 2D classified and subjected to 3D refinements enforcing T symmetry. Particles were further classified using heterogeneous refinement, and the best classes were used for 3D refinements enforcing T symmetry. For the T33-ml23, we obtained an overall resolution of 2.0 Å,\ based on an FSC threshold of 0.143. For the T33-ml28, we obtained an overall resolution of 2.7 Å, for T33-ml30 the resolution was 4.2 Å and for T33-ml35, the resolution was 2.9 Å. Subsets of particles missing one trimer were isolated from the heterogeneous refinement step. 3D refinement with C1 symmetry of these particles missing a trimer resulted in cryo-EM maps for T33-ml23 at 3.9 Å and for T33-ml35, at 4.4 Å. Model building was performed using Coot ^78^, and automated refinement was performed using Phenix ^79^. Figures were prepared using ChimeraX ^80,81^.

## Data processing

### Molecular replacement and refinement of T330fn10 in phaser

Molecular replacement was needed to solve for the phases of the T33-fn10 crystal. A search was carried out in phaser ^82^ using the computationally designed A:B, asymmetric unit model of the cage. Initial hits suggested the A:B model was present, but the signal was marginal. Models with increasing copy numbers of A:B yielded improved results with the model containing eight copies of each of A and B. Upon further inspections, pores in the lattice and omit maps indicated that additional molecules were present in the lattice. Another copy of the tetrahedral model was manually placed in the lattice and oriented in a non-clashing orientation. A final round of molecular replacement with the new model improved the fit further as judged by difference maps without significant features.

Refinement of the initial molecular replacement model was carried out using phenix ^79^. The refinement parameters that yielded the best fit with the smallest deviation between R_work_ and R_free_ included one round of rigid body refinement with grouped ADP. Next, all B-factors were manually set to 200 and another round of rigid body refinement was performed using a reference model as a restraint, and subsequently refining by TLS and grouped ADP. This resulted in an R_work_ of 0.22 and R_free_ of 0.25.

### Cryo-EM map refinement

All maps were refined utilizing their corresponding design assembly model as the search model and the highest resolution map achieved from CryoEM data processing in phenix.real_space_refine ^83^. First, the refine method `run=rigid_body` was used with `resolution`=4.2, `weight=4, <monospace>ncs_constraints`=True, `ramachandran_restraints`=True, `c_beta_restraints`=True, `target_bonds_rmsd`=0.02, and `target_angles_rmsd`=2.0. Rigid body refinement was carried out for five macrocycles, and in some cases longer until the measured CC converged when the design and model had no large deviations. Next, `run=minimization_global+local_grid_search+morphing+adp` was used to refine individual atomic positions with the same flags as for `rigid_body`. After the first round of individual atomic parameters were refined, the model and map were opened for manual inspection in coot ^84^. Deviating side chains and backbone segments identified by visual inspection and validation tools were refined into the density with the Real Space Refine Zone, and model termini were built or removed depending on additional sequences added for affinity purification. Finally, edits in the captain NCS chain were propagated to all mate chains symmetrically and the file was saved for additional rounds of individual atomic refinement.

In most cases, after rigid body phenix refinement, individual chains demonstrated minor asymmetry when superimposed on symmetrically related copies, and were thus asymmetrically fit into the density. Extra measures were taken to find the best fitting individual chains, one for each protein entity, and then the representative chains were manually symmetrized to fit in the map. This procedure improved the overall fit and allowed individual atomic site modifications to reach the highest obtainable CC.

### SEC-SAXS

Initial processing of raw images proceeding according to the automated processing suite from SasTool which includes creating images with background subtraction, the calculation of 1D scattering intensity profiles, and radius of gyration measurements. Subsequent processing was performed to analyze the experimental results in the context of structural models. First, UV absorbance from SEC was used to identify peaks of interest for cage species. Next, SAXS images representing radial scattering intensities from the central fractions of the corresponding SEC peaks were analyzed using PRIMUS from the ATSAS Data analysis software (version 3.2.1) ^85^. The frames of interest were averaged together using the average tool to create average scattering intensities over the identified peak area. To calculate theoretical scattering profiles, the CRYOSOL model evaluation tool was used to convert .pdb files to scattering curves. Theoretical curves were superimposed on the scattering profile of the experimental frames and the scattering values were exported.

## Abbreviations

ASU: asymmetric unit
BSA: buried surface area
LDDT: local distance difference test
RMSD: root mean square deviation

