## Supplementary Materials for "A Suite of Designed Protein Cages Using Machine Learning Algorithms and Protein Fragment-Based Protocols"

#### Supplementary figure 1

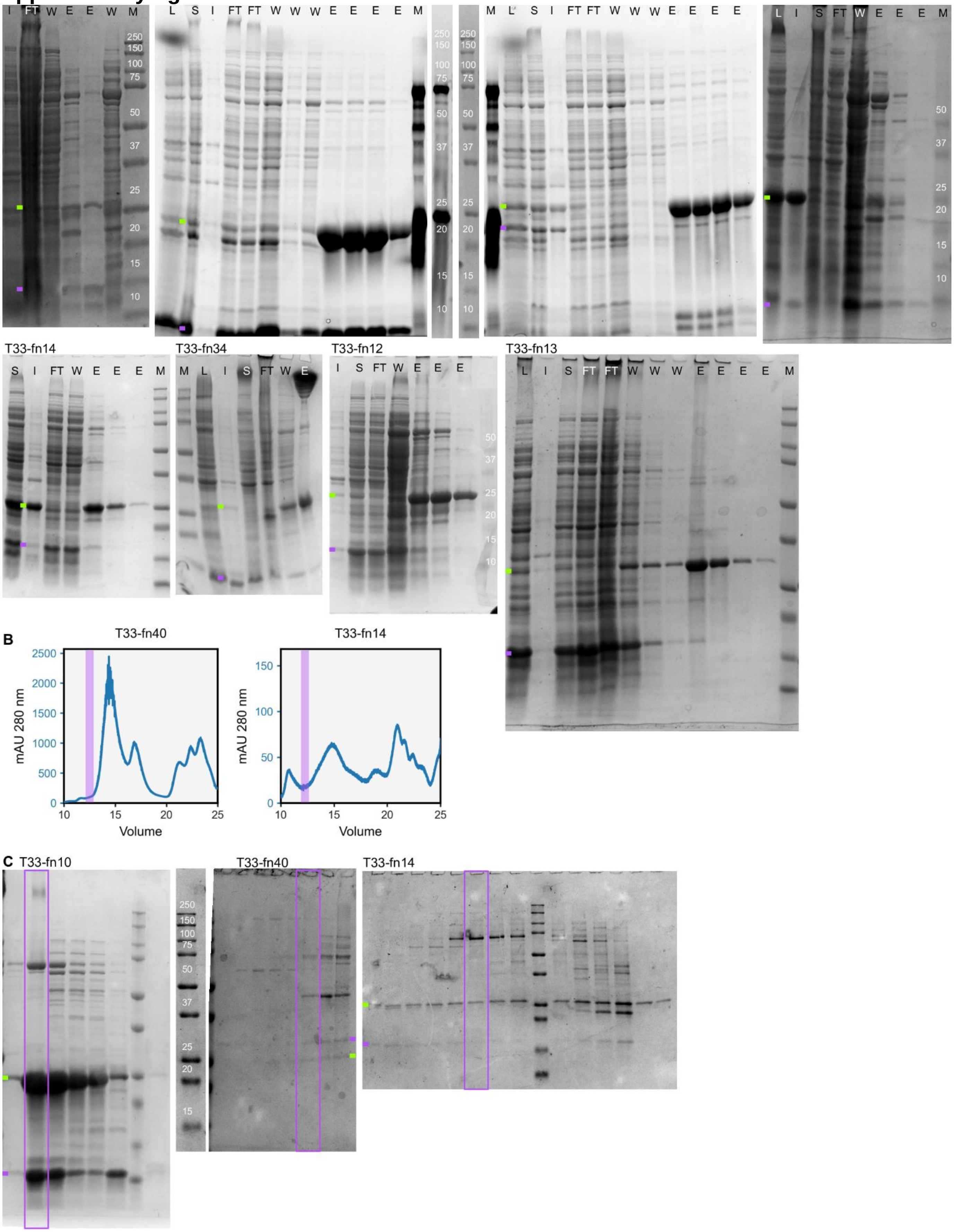

Purification of T33-fn designs.

**a)** IMAC purification gels for selected designs. Samples are presented to indicate the extent of solubility after centrifugation where clarification leaves insoluble material as precipitation. Further, purification reveals the extent to which species co-elute from IMAC. Each gel is labeled according to the following possible labels to indicate the time point in which it was taken. M - molecular weight marker, L - lysate, I - insoluble lysate fraction, S - soluble lysate fraction, FT - soluble lysate after flowing through a Ni-NTA column, W - wash of the Ni-NTA column with 60 mM imidazole, E - elution of the Ni-NTA column with 250 mM imidazole. Multiple fractions indicate time resolved fractionation of the indicated separation type. For each gel, the green bar indicates the experiment size of the component with a His Tag, while the purple bar indicates the experimental size of the second component. **b)** Chromatograms from SEC for selected T33-fn designs. The expected assembly size is overlaid in purple on top of the chromatogram. **c)** SDS-PAGE gels corresponding to SEC runs presented in figure 2. The design T33-fn40 indicates that multiple different species are present at fractions which are larger than trimers. For the designs T33-fn40 and T33-fn14, chromatograms and gels indicate species of assembled cages, intermediates, and trimeric species are present.

**Supplementary figure 2**

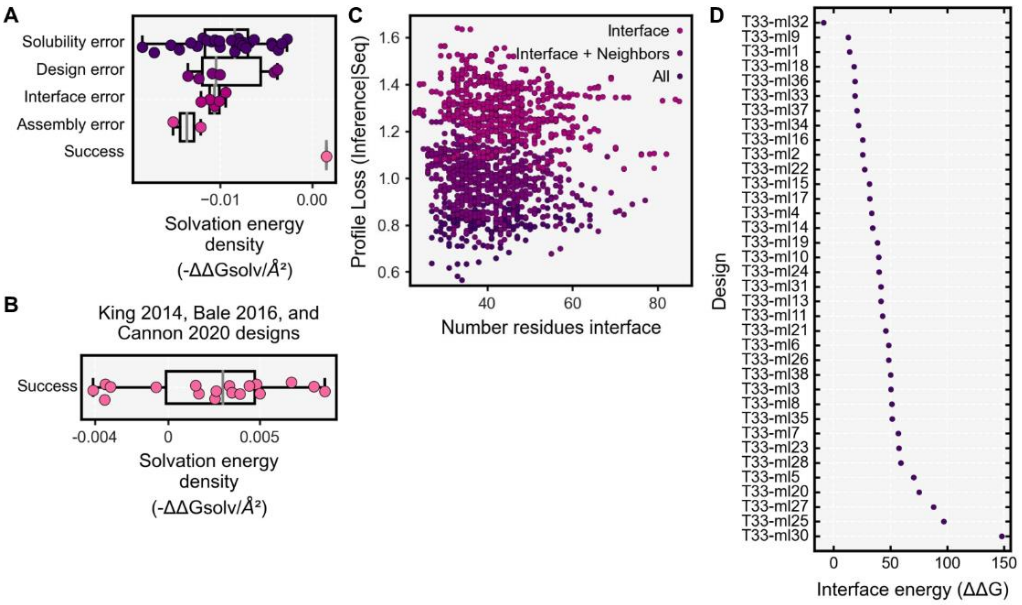


Design calculations of interface free energy and ProteinMPNN score for design models.

**a)** The calculated solvation free energy density values for T33-fn designs are mostly negative while the only design that demonstrates positive solvation energy, *i.e.* solvation is not favored, was T33-fn10, the best characterized design. **b)** For prior successful designs [1–3], calculated solvation free energy density tends to be positive, while designs successfully form with negative calculated values. **c)** The ProteinMPNN score is plotted versus the number of interface residues within 8 Å for each pose. Each datapoint for the ProteinMPNN score is indicated with the color corresponding to the design protocol that was used for the sequence inference. **d)** The interface energy was calculated for each T33-ml design and sorted according to increasing values. All designs have a positive calculated interface energy.

#### AlphaFoldInitialGuess trimeric predictions

Passing:

2zfh_1,3fwu_1,3k9a_1,3l39_1,5izs_1,6pnz_1,6vvw_1,7obj_1,8bro_1

Mixed:

1lw1_1,1m68_1,1mvn_1,1mww_1,1n2m_1,1nkv_1,1otg_1,1p32_1,1qwg_1,1sg4_1,1v4n_1,1v6h_1,1v9o_1,1wp8_1,1xx4_1,1yqf_1,2dtt_1,2dyy_1,2ekm_1,2flz_1,2fvh_1,2h6l_1,2i9d_1,2nmu_1,2p6h_1,2pd2_1,2q35_1,2qs7_1,2vx2_1,2y77_1,2yw4_1,3ahp_1,3b64_1,3c6v_1,3cyo_1,3dli_1,3dzv_1,3ef8_1,3ejv_1,3h5i_1,3hpd_1,3i2b_1,3i3f_1,3i3u_1,3i7t_1,3io0_1,3jv1_1,3ke5_1,3kjk_1,3mc3_1,3n79_1,3nhv_1,3nke_1,3quw_1,3qv0_1,3rf5_1,3tcr_1,3wfv_1,3x2y_1,4bfc_1,4c82_1,4g9q_1,4i61_1,4iyq_1,4jf3_1,4jgs_1,4jpr_1,4jqs_1,4ki3_1,4lho_1,4nsm_1,4usi_1,4wrb_1,4xcw_1,4y6i_1,5bmo_1,5cxd_1,5dii_1,5ds7_1,5eur_1,5ha6_1,5hrz_1,5joq_1,5ka5_1,5kvb_1,5uif_1,5z1q_1,6as5_1,6bj7_1,6cuq_1,6gdx_1,6j3m_1,6l8p_1,6ln3_1,6lr3_1,6mhh_1,6mmq_1,6qbw_1,6t76_1,6veh_1,6vvr_1,6vw4_1,6x7q_1,7m58_1,7ms9_1,7te3_1,8del_1

Failing:

1avq_1,1hl7_1,1j3l_1,1jlj_1,1jxz_1,1khx_1,1og6_1,1pf5_1,1rhy_1,1ui9_1,1viy_1,1vl0_1,1yox_1,1yx1_1,1zcl_1,2a7k_1,2ar3_1,2c5q_1,2ig8_1,2is8_1,2qlp_1,2ves_1,2vhe_1,3e99_1,3eby_1,3fsc_1,3gkb_1,3gmj_1,3hrx_1,3hyt_1,3irs_1,3k93_1,3kwe_1,3lao_1,3lke_1,3mae_1,3mc4_1,3o3w_1,3r8y_1,3soz_1,3tqf_1,3ub1_1,3vbp_1,3vnp_1,3wv7_1,3zjb_1,4b6r_1,4di1_1,4gdz_1,4isx_1,4kw2_1,4m17_1,4mej_1,4myo_1,4nrd_1,4r7t_1,4rfu_1,4uof_1,4wia_1,4wk3_1,5b2f_1,5fus_1,5jru_1,5m62_1,5o34_1,5ucq_1,5un0_1,5v13_1,5vjy_1,5wfg_1,5xum_1,5z81_1,6cv6_1,6its_1,6ive_1,6lnl_1,6ny9_1,6p7l_1,6p7o_1,6tj2_1,6ty6_1,6we5_1,6wmg_1,6zzm_1,7c1i_1,7cp2_1,7dsz_1,7l7w_1,7o45_1,7okc_1,7rgv_1,7std_1,7tbp_1

### Supplementary figure 3

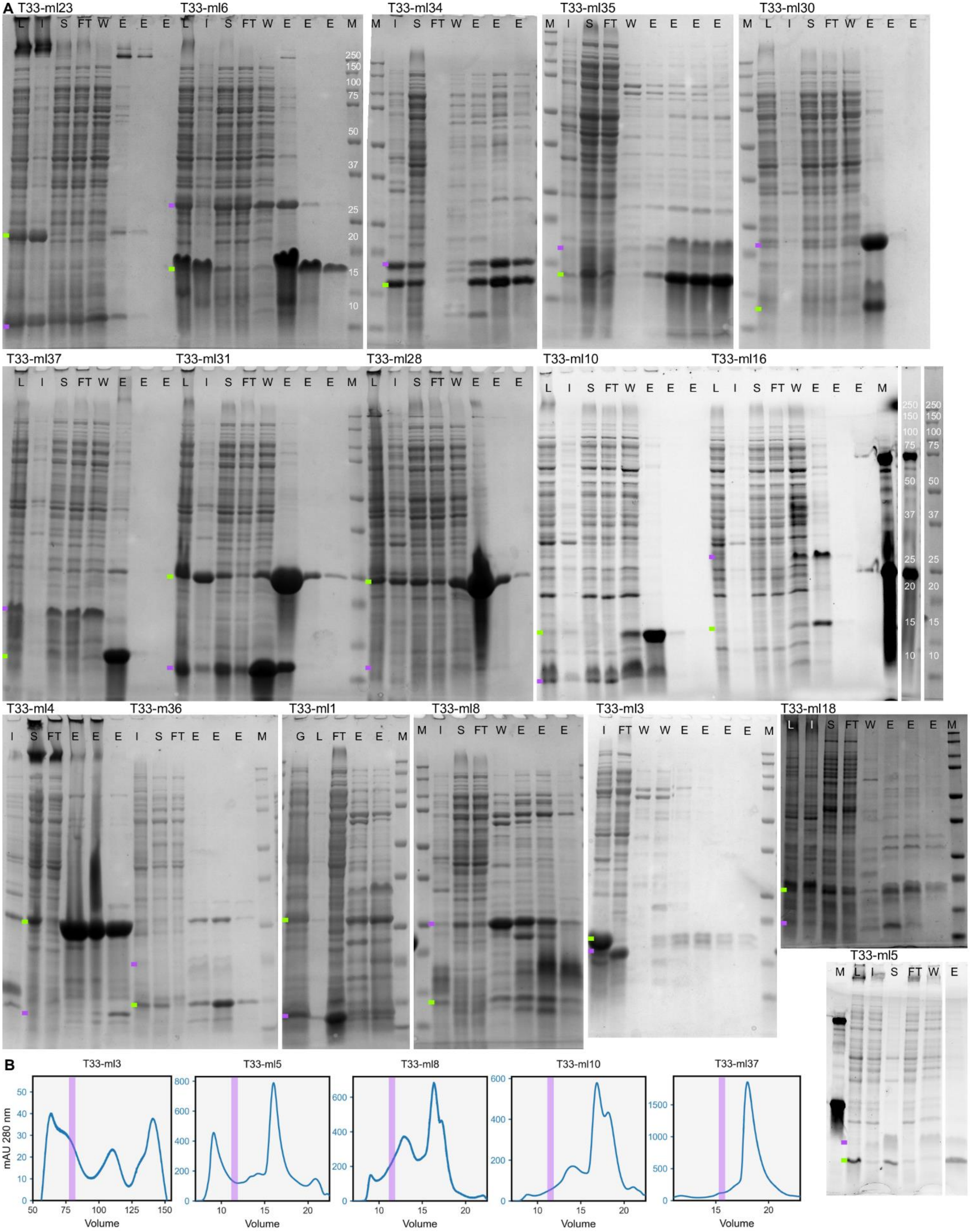

Immobilized metal affinity chromatography of T33-ml designs.

**a)** IMAC purification gels for selected designs. Samples are presented to indicate the extent of solubility after centrifugation where clarification leaves insoluble material as precipitation. Further, purification reveals the extent to which species co-elute from IMAC. Each gel is labeled according to the following possible labels to indicate the time point in which it was taken. M - molecular weight marker, L - lysate, I - insoluble lysate fraction, S - soluble lysate fraction, FT - soluble lysate after flowing through a Ni-NTA column, W - wash of the Ni-NTA column with 60 mM imidazole, E - elution of the Ni-NTA column with 250 mM imidazole. Multiple fractions indicate time resolved fractionation of the indicated separation type. For each gel, the green bar indicates the experiment size of the component with a His Tag, while the purple bar indicates the experimental size of the second component. **b)** Chromatograms from SEC for selected designs classified as an assembly error. The expected assembly size (~11-12 ml) is overlaid in purple on top of the chromatogram.

### Supplementary figure 4

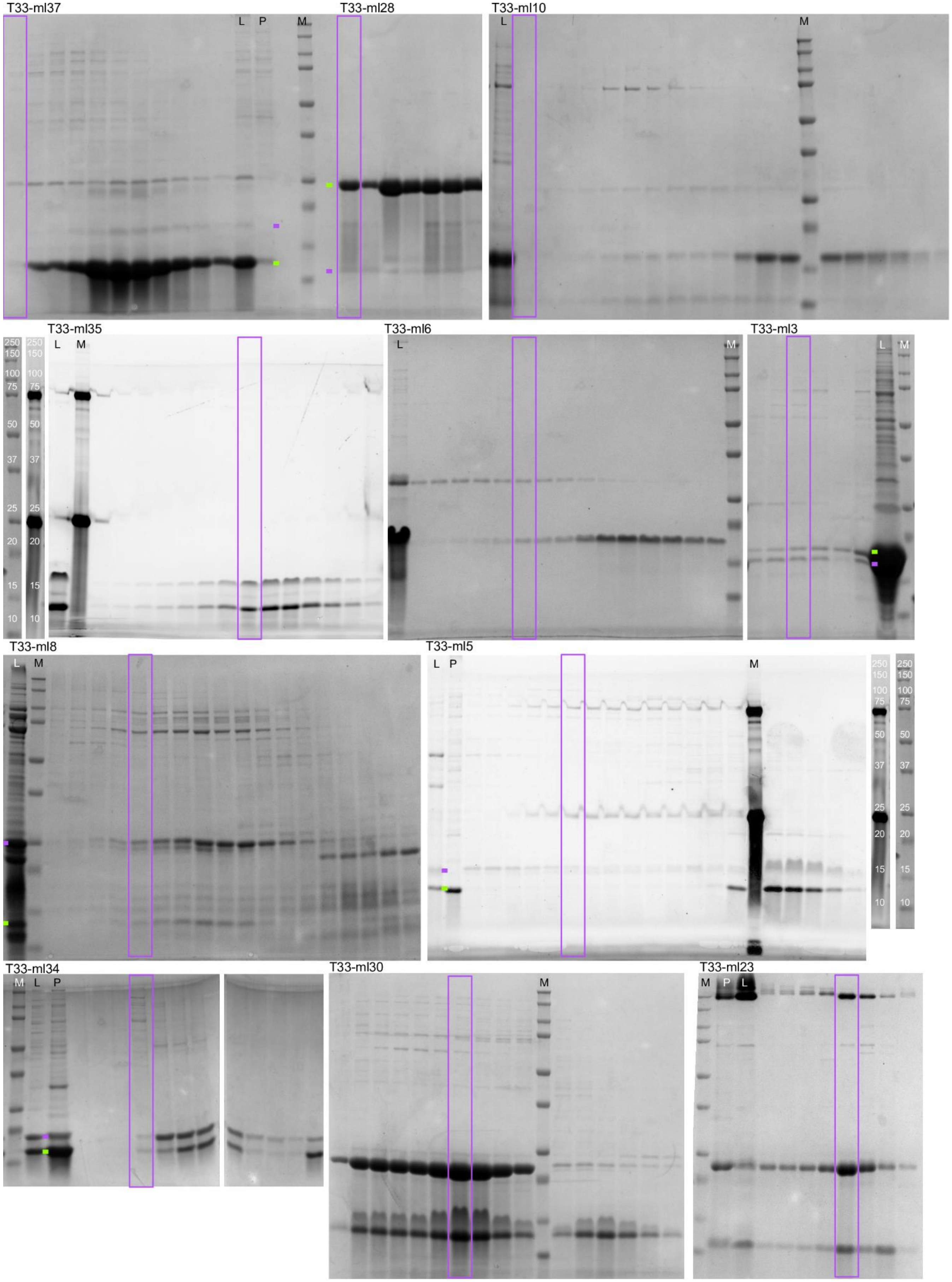

Separation of T33-ml assemblies using size exclusion chromatography.
**a)** SDS-PAGE gels corresponding to SEC runs presented in Figure 4a and Figure S3b. The expected assembly fraction (~11-12 ml) is overlaid in purple on top of each gel and lanes are indicated as: M - with a molecular weight marker, L - protein loaded to SEC, P - precipitation during protein concentration. The gel of T33-ml34 has been stitched together to reflect the ordering of the lanes in the SEC experiment. Despite robust co-elution, SEC results in many species, from assembled cages, trimeric species bound together and even monomers which are unbound. **b)** The designs T33-ml3, T33-ml5, T33-ml8, T33-ml10, and T33-ml37 mostly elute from the size exclusion at the incorrect fraction, which is indicative of *assembly errors* that prevent full assembled cages from forming, while resulting in larger assemblies as a result of protein complexation.

#### Supplementary figure 5

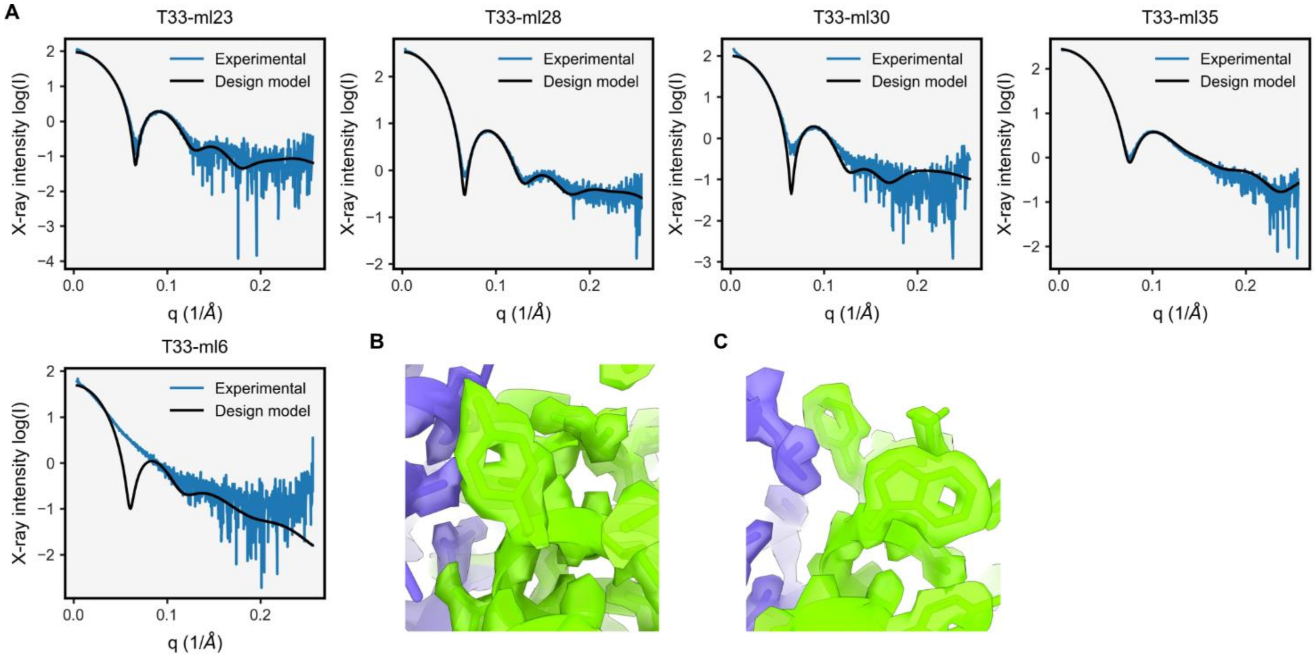


Validation of design models using small angle x-ray scattering and cryo-EM density.
**a)** Averaged experimental small angle x-ray scattering (SAXS) profiles for images acquired from SEC peak fractions corresponding to assembled cages. The SAXS profiles (blue) correspond to images 238-258, 239-255, 229-249, 239-256, and 235-241 for T33-ml23, T33-ml28, T33-ml30, T33-ml35, and T33-ml6, respectively. Each scattering profile is plotted alongside the theoretical scattering calculated from design models (black). **b-c)** For the design T33-ml23, the 2.0 Å resolution allows aromatic side chains to be resolved. Example residues include component A Y85 (panel b) and W112 and F115 (panel c).

**Supplementary figure 6**

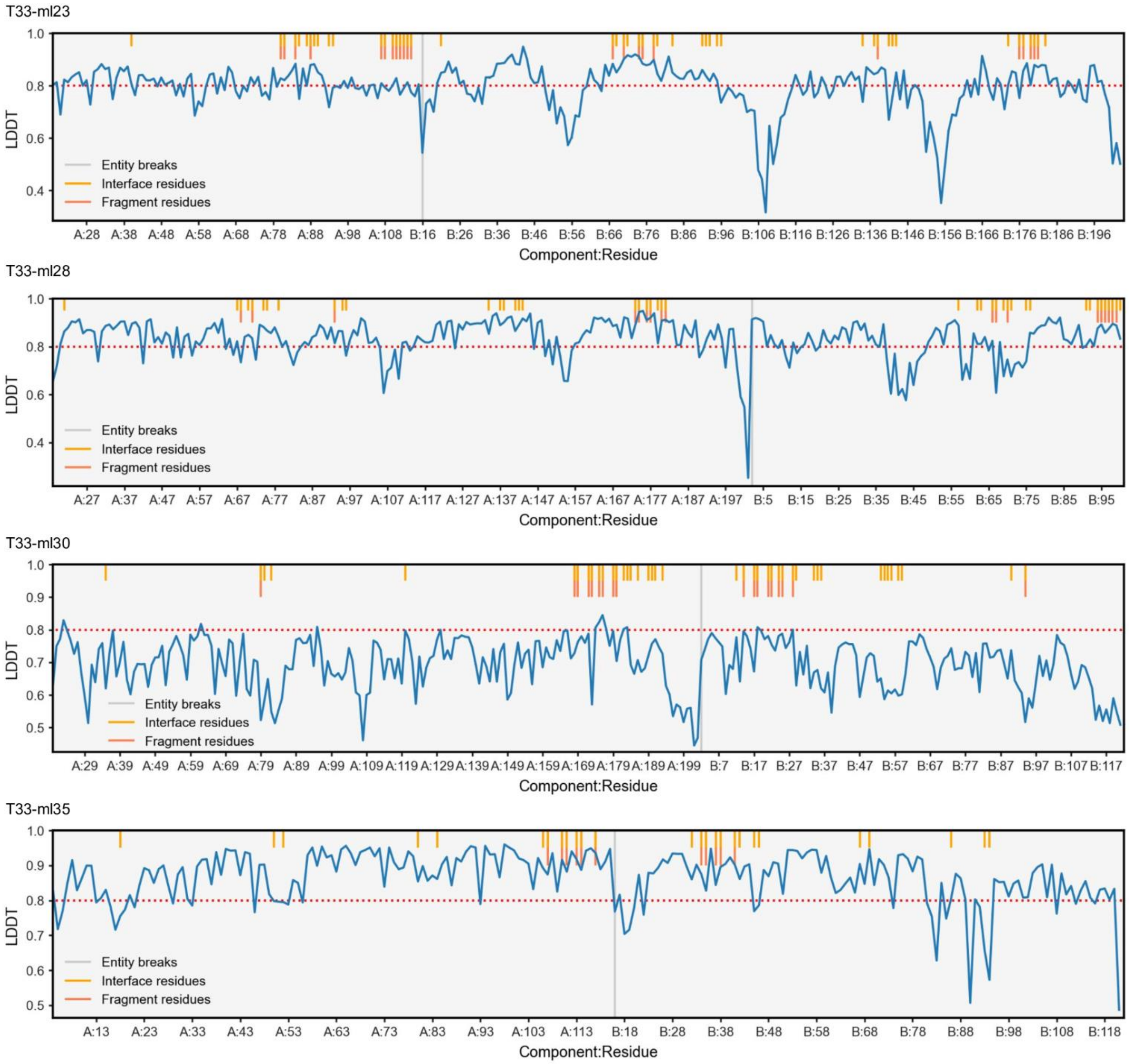


Per-residue local density difference test for interface and interface fragment residues.

For the indicated designs, the local distance difference test (LDDT) was measured between the full cryo-EM assembly structure and the predicted design model. Values with greater than 0.8 (horizontal red line) indicate strong agreement. The location of interface residues (orange bar, upper segment; identified by 8 Å C-beta C-beta distances) and fragment residues (red bar, lower segment) are highlighted to indicate the extent of agreement in the modeled *de novo* interface. Entity breaks separate the A and B chains in each assembly.

#### Supplementary figure 7

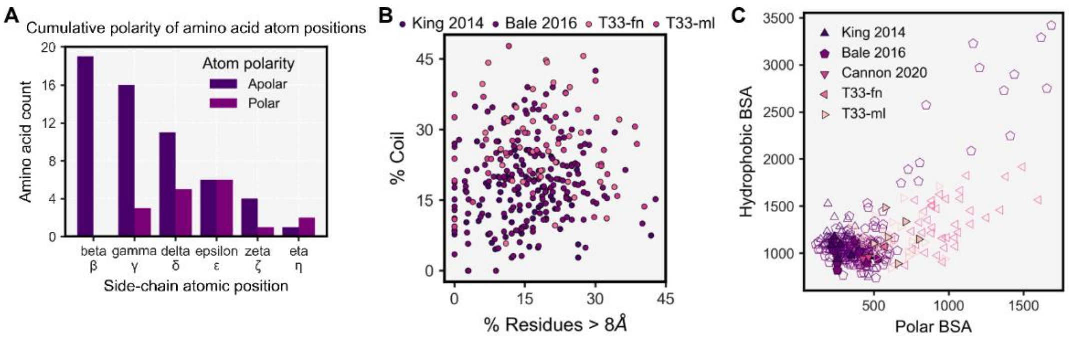


Retrospective analysis of polar interactions and interface areas for two-component designed protein cages. **a)** Histogram of atomic polarity by side-chain position. For each canonical amino acid (n=20), the polarity of each atom in the side-chain (polar: N, O, S; apolar: C, H) is plotted at the side-chain atomic position it occupies. Polar atoms more frequently occupy atomic positions distal to the alpha carbon (Ca). **b)** The distribution of polar conformational features for design models. The fraction of coiled residues is independent of the fraction of residues greater than 8 Å apart as measured by Cb-Cb distance. **c)** Comparison of the interface area contributions from hydrophobic and polar buried surface area (BSA) interactions for 2-component designed cages considered in this analysis. The markers denote the symmetry of the assembly (triangle - tetrahedral, pentagon - icosahedral) and are filled with color if the assembly is reported as successful.

**Supplementary sequences**

T33-fn1-A
MKLIVAIVRPEKLYDVLRFLFHAGVRGLTLSRVQGHGGETERVETYRGTTVKMEFAEKVRLEIGVSEPFVEATV
IAILIAARTGEVGDGKIFVLPVEKVYRIRTGEEDEAAVTPVQ
T33-fn1-B
MDAQSAAKCLTAVRRHSPLVHSITNNVVTNFTANGLLALGASPVMAYAKSEVADMAKIAGALVLNIGTLSKESL
LAMTAAGLSANEHGVPVILDPVGAGATPTRTLAARFIIHAVRLAAIRGNAAEIAHTVGVTDWLIKGVDAGEGGG
DIIRLAQQAAQKLNTVIAITGEVDVIADTSHVYTLHNGHKLLTKVTGAGCLLTSVVGAFCAVEENPLFAAIAAISS
YGVAAQLAAQQTADKGPGSFQIELLNKLSTVTEQDVQEWATIERVTVSGHHHHHH
T33-fn2-A
MSLDYTTQQIIEKLRELKIVPVIALDNADDIIPLALTLAAAGLSVAEITFRSEAAADAIRLLRKISPDFLIAAGTVLTA
EQVHRAKRSGADFVVTPGLNPKIVKLCQDLNFPITPGVNNPMAIEIALEMGISAVKFFPAEASGGVKMIKALLG
PYAQLQIMPTGGIGLHNIRDYLAIPNIVACGGSWFVEKKLIQSNNWEEIATLVKEVIDIIK
T33-fn2-B
MHHHHHHSGAYRYRDIVVRKQDGFTHILLSTKSSENNSLNPEVMREVQSALSTAAADDSKLVLLSAVGSVFC
CGLDFIYFIRRLTDDRYRESTLMALAIAYFVNTFIQFKKPIIVAVNGPAIGLGASILPLCDVVWANEKAWFQTPYT
TFGQSPDGCSTVMFPKIMGGASANEMLLSGRKLTAQEACGKGLVSQVFWPGTFTQEVMVRIKELASCNPVV
LEESKALVRCNMKDELIQANVREALVLTKIWGSAQGMDSMLKYLQRKIDEF
T33-fn3-A
MSLSYTTQQIIEKLKRLGIVPVIALDNADDILPLADTLAKNGLSVAEITFRSEAAADAIRLLRANRPDFLIAAGTVLT
AEQVVLAKSSGADFVVTPGLNPKIVKLCQDLNFPITPGVNNPMAIEIALEMGISAVKFFPAEASGGVKMIKALLG
PYAQLQIMPTGGIGLHNIRDYLAIPNIVACGGSWFVEKKLIQSNNWKEIAYLVLEVLIIIGE
T33-fn3-B
MHHHHHHSGMMTTSNAGAQQPNVEGRRFSPDQVRSVAPALEQYTQQRLYGDVWQRPGLNRRDRSLVTIA
ALIARGEAPALTYYADQALENGVKPSEISETITHLAYYSGWGKAMATVGPVSEAFAKRGIGQDQLAAVESTPL
PLDEEAEQQREDRVTRQFGSVAPGLVQYTTDYLFRDLWLRPDLAPRDRSLVTIAALISVGQVEQITFHLNKAL
DNGLSLDQAAEVITHLAFYAGWPNAMSALPVALAVKFKRHS
T33-fn4-A
MSLSYTTQQIIEKLRELKIVPVIALDDYRDILGLAMVLAANGLSVAEITFRSEAAADAIRALRIFHPDFLIAAGTVLT
AEQVVLAKSSGADFVVTPGLNPKIVKLCQDLNFPITPGVNNPMAIEIALEMGISAVKFFPAEASGGVKMIKALLG
PYAQLQIMPTGGIGLHNIRDYLAIPNIVACGGSWFVEKKLIKRNNGDEIARLVREVIDIIKE
T33-fn4-B
MHHHHHHSGDALVDYAGPAATGGNVARLTLNSPHNRNALSSALVSQLHQGLRDASSDPAVRVVVLAHTGGT
FCAGADLSEAGSGGSPSSAYDMAVERAREMAALMRAIVESRLPVIAAIDGHVRAGGFGLVGACDIAVAGPRS
SFALNEAAIGVAPAIISLTLLPKLSARAAARYYLTGSQFDAREAEEIGLITAAALIVLAQVLMLAGAVISGSPQGLA
ASKALTTAAVLERFDRDAERLAEESARLFVSDEAREGMLAFLENRAPNWFS
T33-fn5-A
METVETSAAPKPDGPYSQAIKVGNTLYVSGQIPIDEQTNTIVDGDIATQTAQVLLNIMAIVLAAGFSLSDVAMAF
VFLKDMNMFEDFNQTYALAFTDKPPARVTVEVSRLPKDALIEIAVICSKGSHHHHHH
T33-fn5-B
MAYRYRDIVVRKQDGFTHILLSTKSSENNSLNPEVMREVQSALSTAAKDSSKLVLLSAVGSVFCCGLDFIYFIR
RLTDDRLTESRKMAEAIRNFVNTFIKFQKPIIVAVNGPAIGLGASILPLCDVVWANEKAWFQTPYTTFGQTPDG
CSTVMFPKIMGGASANEMLLSGRKLTAQEACGKGLVSQVFWPGTFTQEVMVRIKELASCNPIVLKQSKILVRS
NMERELEKANDLEAYVLSKIWASAQGMDSMLKYLQRKIDEF
T33-fn6-A
MAYRYRDIVVRKQDGFTHILLSTKSSENNSLNPEVMREVQSALRKAQADESKLVLLSAVGSVFCCGLDFIYFIR
RLTDDRLDEAKKMAEAIRNFVNTFIQFTKPIIVAVNGPAIGLGASILPLCDVVWANEKAWFQTPYTTFGQSPDG

CSTVMFPKIMGGASANEMLLSGRKLTAQEACGKGLVSQVFWPGTFTQEVMVRIKELASCNPGVLIVSKALVR
SNMEMELEKANKLEAAVLLAIWALDDGMDSMLKYLQRKIDEF
T33-fn6-B
MHHHHHHGSLVRRIIFTDKAPDAIGAYSQAVLVDRTIYISGQLGMDPASGKLVPGGVIAETAQALLNILEILRAA
GCRMTNVVKATVLLADINDFLDVAIVIAGFHTRSFPARAAYQVAALPKGGRVEIEAIAVQGPLTTASL
T33-fn7-A
MHHHHHHSGAYRYRDIVVRKQDGFTHILLSTKSSENNSLNPEVMREVQSALSTAAHDDSKLVLLSAVGSVFC
CGLDFIYFIRRLTDDRSRESLKMAEAIRNFVNTFIQFDKPIIVAVNGPAIGLGASILPLCDVVWANEKAWFQTPYT
TFGQSPDGCSTVMFPKIMGGASANEMLLSGRKLTAQEACGKGLVSQVFWPGTFTQEVMVRIKELASCNPVV
LEISKALVRTNMEEELEQANADECSSLAYIWGLAQGMDSMLKYLQRKIDEF
T33-fn7-B
MPMFIVNTNVPRASVREGFLRRLTNALAAYTGKHRQYIAVHVVPDQLMTFSGTNDPCALCSLHSIGKIGGEQN
AALSAYLCILLSDDLKISPDRVYINYYDMNAANVGWNGDTFA
T33-fn8-A
MHHHHHHSGAYRYRDIVVRKQDGFTHILLSTKSSENNSLNPEVMREVQSALRTALYDDSKLVLLSAVGSVFC
CGLDFIYFIRRLTDDRKRESTKMAEAIKKFVITFIFFAKPIIVAVNGPAIGLGASILPLCDVVWANEKAWFQTPYTT
FGQSPDGCSTVMFPKIMGGASANEMLLSGRKLTAQEACGKGLVSQVFWPGTFTQEVMVRIKELASCNPVVL
KESKHLVRLNMLEELYKANERECEVLKKIWGSAQGMDSMLKYLQRKIDEF
T33-fn8-B
MQWQTKLPLIAILRGITPDEAALHVAAVIKAGFDAVEIPLNSPQWEQSIPMIVAIYGEVALIGAGTVLKPEQVDAL
ARMGCQLIVTPNIHSEVIRRAVGYGMTVCPGCATATEAFTALEAGAQALKIFPSSAFGPQYIKALKAVLPSDIAV
FAVGGVTPENLAQWIDAGCAGAGLGSDLYHQSLSLDATAKQALEFVEAYKRAVAL
T33-fn9-A
MHHHHHHSGMKVVVQIKDFDKVPQALRSVANLYADIKDAEIEVVLHQSAIKALLQDSPTRDITSLLIKANILIVGC
ENSIRSQNLSHDQLQPGIKIVTSGVGEIVRKQSEGWIYLAL
T33-fn9-B
MPGMLPWTEQQFQLLGEIEEVELGRIQKRSGANLSRNWVMIPHVTHFDKTDITELEAFRKQQNEEAAKRKLD
VKITPVVFIMKAVAAALEQMPRFNSSLSEDGQRLTLKKYINIGVAVDTPNGLVVPVFDDVNKKGIIELSRELMTIS
KKARDGKLDTSEMVGGCFTISSIGGLGTTHFAPIVNAPEVAILGVSKSAMEPVWNGKEFVPRLMLPISLSFDHR
VIDGADGARFITIINNTLSDIRRLVM
T33-fn10-A
MKVVVQIKDFDKVPQALQSVLNLFLDLGNAEIEVVLHQSAIKALLLNSPTRSIIEELIKLNILIVGCEHSIRSQNLD
HRQLIDGIKIVRSGVGEIVRKQSEGWIYLAL
T33-fn10-B
MHHHHHHSGMSLRLERDGAVARLLIDRADRRNAFSLDMWQRLPELLAEASGDDALRVLVVKSANGGAFCAG
ADIAELLANKDDAAFHLANQQAINRAQYELARFRLPTVAMVEGDCIGGGCGIALACDMRIAAPAARFGITPAKL
GLVYPLHDVKLLVDLVGPGQARRLMFTGGLIDANEAHRIGLVELLGESEDALVGQLATVSSFSTQAIKSFVRRV
LDGQVADDTLSLCVFASATLGADFREGTGAFLEKRPPVF
T33-fn11-A
MHHHHHHSGTDITANVVVSNPRPIFTESRSFKAVANGKIYIGQIDTDPVNPANQIPVYIENEDGSHVQITQPLIIN
AAGKIVYNGQLVKIVTVQGHSMAIYDANGSQVDYIANVLKYDPDQYSIEADKKFKLIKQIEDKIQLILAAIASILRD
LARIWKLIGE
T33-fn11-B
MSIDKLKHKLDDYAKDIKLNLSSITRSSVLDQEQLWGTLLASAAATRNKQVLADIKLDSTLYLDQREQHAALGA
AAIMGMNNVFYRGRGFLEGRYDDLRPGLRMNIIANPGIPKANFELWSFAVSAINGCSHCLVAHEHTLRTVGVD
REAIFEALKAAAIVSGVAQALATIE
T33-fn12-A

MTDITANVVVSNPRPIFTESRSFKAVANGKIYIGQIDTDPVNPANQIPVYIENEDGSHVQITQPLIINAAGKIVYNG
QLVKIVTVQGHSMAIYDANGSQVDYIANVLKYDPDQYSIEADKKFKLIKQIEDKIEKILAAIAHIEIDIALIKALIGE
T33-fn12-B
MHHHHHHSGDDPRLLSLFSAQREEDADIVIIGFPYDEGCVRNGGRAGAKKGPAAFRFFLQRLGSVENRELNV
NASHLKLYDAGDITASTLEEAHEKLESKVFTVLARGAFPFVIGGGNDQSAPNGRAMLRAFPGDVGVINVDSHL
DVRPPLSDGRVHSGTPFRQLLEESSFDGSRFVEFACQGSQCGALHAAYVQANQGHLMWLSEVRKKGAVRA
LAEAFKITGKNTFFSFDVDSLKSSDMPGVSCPAAVGLSAQEAFDMCFLAGSISTVMMMDMSELNPLVEEYRS
PRVAVYMFYHFVLGFATRP
T33-fn13-A
MTDITANVVVSNPRPIFTESRSFKAVANGKIYIGQIDTDPVNPANQIPVYIENEDGSHVQITQPLIINAAGKIVYNG
QLVKIVTVQGHSMAIYDANGSQVDYIANVLKYDPDQYSIEADKKFKLIAQIEGHLEAIASVLQSIINEIARIKKLIGE
T33-fn13-B
MHHHHHHSGDDPRLLSLFSAQREEDADIVIIGFPYDEGCVRNGGRAGAKKGPAAFRFFLQRLGSVNNLELNV
DASHLKLYDAGDITASTLEEAHEKLESKVFTVLARGAFPFVIGGGNDQSAPNGRAMLRAFPGAVGVINVDSHL
DVRPPLSDGRVHSGTPFRQLLEESSFDGRFFVEFACQGSQCGALHAQYVRDHQGVLMWLSEVRALGAVKA
LRLAFTLTGANTFFSFDVDSLKSSDMPGVSCPAAVGLSAQEAFDMCFLAGKTPEVMMMDMSELNPLVEEYR
SPRVAVYMFYHFVLGFATRSKPKAEN
T33-fn14-A
MTDITANVVVSNPRPIFTESRSFKAVANGKIYIGQIDTDPVNPANQIPVYIENEDGSHVQITQPLIINAAGKIVYNG
QLVKIVTVQGHSMAIYDANGSQVDYIANVLKYDPDQYSIEADKKFKLIKQIEDKIQQILEKIAIIFRQLARIAKYIGE
T33-fn14-B
MHHHHHHSGMSLRLERDGAVARLLIDRADRRNAFSLDMWQDLPKLLAEARSDSALRVLVVKSANGGAFCAG
ADIAELLANKDDALFHEENQEAINRAQYELARFRLPTVAMVEGDCIGGGCGIALACDMRIAAPAARFGITPAKL
GLVYPLHDVKLLVDLVGPGQARRLMFTGGLIDANEAHRIGLVELLGESEDALVGQLATVSSFSTQAIKSFVRRV
LDGQVADDAHSLNVFHLAFMTDDFREGTGAFLEKRPPVF
T33-fn15-A
MPFLQTIVSVSLDDQKRARLSLFYGMLCRKTLGIPGDQVMTAFSDKTPISFNGSTAPAAYVRVESWGEYAPSK
PKEMTAAIAAAIYAECGIPPERIYVFYYSTKHCGWNGHNF
T33-fn15-B
MHHHHHHSGAYRYRDIVVRKQDGFTHILLSTKSSENNSLNPEVMREVQSALSTAAADSASLVLLSAVGSVFC
CGLDFIYFIRRLTDDTKRESFKMAEAIRNFVNTFIQFQKPIIVAVNGPAIGLGASILPLCDVVWANEKAWFQTPYT
TFGQSPDGCSTVMFPKIMGGASANEMLLSGRKLTAQEACGKGLVSQVFWPGTFTQEVMVRIKELASCNPAV
LRESKFLVRCNMKMELEQANEREAAHLKFIHAHAQGMDSMLKYLQRKIDEF
T33-fn16-A
MPFLQTIVSVSLDDRKRALLSTAYLYICREELGLALDSVMTAFSDKTPISFDGSTEPAAYVRVESWGEYAPSKP
KMMTPRIAAAITKECGIPKARIYVFYYSTKHCGWNGTNF
T33-fn16-B
MHHHHHHSGDALVDYAGPAATGGPVARLTLNSPHNRNALSSALVSQLHQGLRDASSDPAVRVVVLAHTGGT
FCAGADLSEAGSGGSPSSAYDMAVERAREMAALMRAIVESRLPVIAAIDGHVRAGGFGLVGACDIAVAGLES
SFALYEARIGVAPAIISLTLLPKLSARAAARYYLTHEKFDARRAEEIGLITMAAEDVDLLVALLVLAVGSGSPQGL
AASKALTTAAVLERFDRDAERLAEESARLFVSDEAREGMLAFLEKRLPNWFS
T33-fn17-A
MHHHHHHSGMVLKERQDGVLVLTLNRPEKLNAITGELLDALYAALKEGEEDREVRALLLTGAGRAFSAGQDL
TEFGDHIPQYEDHLRRYNRVVEALSGLNKPLVVAVNGVAAGAGMSLALWGDLRLAAVGASFTTAFVRIGLVP
DSGLSFLLPRLVGLAKAQELLLLSPRLSAEEALALGLVHRVVPAEKLMEEALSLAKELAQGPTRAYALTKILLLE
TYRLSLTEALALEAILQGFAGLTKDHEEGVRAFREKRPPRFQGS
T33-fn17-B
MIVQQQNNLLRAIEAQEALLQLTVIGIKRLQARSGGRGGWETLERLIKKYTSTIASLIAESQNQQEK

T33-fn18-A
MHHHHHHSGMVLKERQDGVLVLTLNRPEKLNAITGELLDALYAALKEGEEDREVRALLLTGAGRAFSAGQDL
TEFGDHHPRYGSHLQRYNRVVEALSGLEKPLVVAVNGVAAGAGMSLALWGDLRLAAVGASFTTAFVRIGLVP
DSGLSFLLPRLVGLAKAQELLLLSPRLSAEEALALGLVHRVVPAEKLMEEALSLAKELAQGPTRAYALTKKLLL
ETYRLSLTEALALERLAQAIAGMSQDHEEGVRAFREKRPPRFQGR
T33-fn18-B
MPLIRIDLTSDRSREQRRAIADAVHDALVEVLAIPARDRFQILTAHDPSDIIAEDAGLGSDKSPSVVIIHVFTQAG
RTIETKQRVFKAITLSLLPIGVMDADVFIAITENAPHDWSFAGGQNQYVQGELAIPATGAA
T33-fn19-A
MVLKERQDGVLVLTLNRPEKLNAITGELLDALYAALKEGEEDREVRALLLTGAGRAFSAGQDLTEFGDRKPDY
EAHLRRYNRVVEALSGLEKPLVVAVNGVAAGAGMSLALWGDLRLAAKGASFTTAFVRIGLVPDSGLSFLLPRL
VGLAKAQKLLLDSLKLSAEQALKLGLVHGVVKAHALMLEALLIARRLAQGPTRAYALTKKLLLETYRLSLTEALA
LEAVLQGQAGQTQDHEEGVRAFREKREPRFQGR
T33-fn19-B
MHHHHHHSGMTQTAPAAVAYSVNHAGVAAIVLDRPDASNALDEHMKTELLQALLAAGGDPAVRAVVMSAAG
KNFCVGQDLEEHVERLDDDPAHAMDTVREHYNPVLEALDAIKVPVVVAINGACVGAGLGLALGADIRIAGQRA
KFGTAFTGIGLAADSALSASLPRLIGASRATAMFLLGDTIDAPTAHTWGLVHEVVDEGSPADVANSVAGRLAG
GPTAAFSEVKELLRRNAVAPLGDVLEREASAQQRLGASVDHSAAVLAFEAKDKPIFYGD
T33-fn20-A
MAVSDQRLSEATKRELQDELQRAGHPQAPVIPDGWRMDFELGVTHFTMRKSHGDEEIILQLTGEDRSNEEIT
RTLDVLVVNGGKALVFGMSVEDGEFVINNVCFRHDGKLALDTSAEAQFQKSQLYMGPDLADLEDHLVDSFTS
YLSARGVNDTLANFIDQASLVFEQQNYLAWLLAINLFVS
T33-fn20-B
MLSVNEIAAEIVEDMLDYEEELRIESKKLSTGAIVVDCGVNVPGSYDAGIMYTQVCMGGLADVDIVVDTINDVP
FAFVTEYTDHPAIACLGSQKAGWQIKVGKYFAMGSGPARALALKPLETMARIEYMDDARVAVIALEANQLPDD
RHMTYMAIECFVRLENVYALVAPTASIVGSVQISGRIVQTAIFKMNEIGYDPKLIVSGAGRCPISPILENDLKAMG
STNDSMMYYGSVFLTVKKYDEILKNVPSCTSRDYGKPFYEIFKAANYDFYKIDPNLFAPAQIAVNDLETGKTYV
HGKLNAEVLFQSYQIVLEEGSHHHHHH
T33-fn21-A
MADADQVLSAATQLELAIERTRAGLPEKPEIPPGWEIDRKPGVTHFTMRKSHGSETIILQLTGEDRSNEEITRTL
DVLVVNGGKALVFGMSVEDGEFVINNVCFRKDGKLALDTSAEAQFQKSQLYMGPDLADLEEYLVDSFTSYLS
ARGVNDTLANFIDQFSLWSEQADYEEWLESINQFMS
T33-fn21-B
MHHHHHHSGMSLRLERDGAVARLLIDRADRRNAFSLDMWQRLPELLKEASGDDALRVLVVKSANGGAFCAG
ADIAELLANKDDAAFHAANTAAILLAMAELASFRLPTVAMVEGDCIGGGCGIALACDMRIAAPAARFGITPAKLG
LVYPLHDVKLLVDLVGPGQARRLMFTGGLIDANEAHRIGLVELLGESEDALVGQLATVSSFSTQAIKSFVRAVL
DGQIIDTAKSFAVFASAFEGADFREGTGAFLEKRPPVF
T33-fn22-A
MSLSQDGTLEQRQGDGVLTLTLGRAPAHPLSKETLARLKAALEWAMGDDSVHVLVIHGPGRIFCAGHDLKEI
GRHRADPDEGREEVTILFEECSALMLDLAHCPKPTIALVEGIATAAGLQLMAACDLAYASPAARFCLPGVQNG
GFCTTPAVAVSRVIGRRAVTEMALTGATYDADWALAAGLINRILPEAALATHVADLAGALAARNQAPLRRGLE
TLNRHLELPLEQAYALATPVMVEHFMDPGRRHLDWID
T33-fn22-B
MHHHHHHSGSAVQPSFIRTNIGSTLRIIEEPQSDVYWIHMHADLAINPGRACFSTRLVDDITGYQTNLGQRLNT
AGVLAPHVVLASDSDVFNLGGDLALFCQLIREGDRARLLDYAQRCVRGVHAFHVGLGARAHSIALVQGNALG
GGFEAALSCHTIIAEEGVMMGLPEVRFDLFPGMGAYSFMCQRISAHLAQKIMLEGNLYSAEQLLGMGLVDRV
VPPGFGELEIAKHIQKSQLTPHAWAAMQQVREMTTAVPLEEMMRITEIWVDTAMQLGEESLRTMDQIVRKQS
RRSGLDAG

T33-fn23-A
MKKIHTKNAPAARGPYVQGKIVGNLLFASGQVPLSPLSGKVIGTTIEEQTRQVLANIAAILGAAGTDFDHVVKTT
CFLSDIADFLPFNEVYADQFKSDFPARSAVEVARLPKNVKIEIEVIAELI
T33-fn23-B
MHHHHHHSGMMTTSNAGAQQPNVEGRRFSPDQVRSVAPALEQYTQQRLYGDVWQRPGLNRRDRSLVTIA
ALIARGEAPALTYYADQALENGVKPSEISETITHLAYYSGWGKAMATVGPVSEAFAKRGIGQDQLAAVESTPL
PLDEEDERAFATRVRNMFGDVAPGLVQYTTDYLFRDLWLRPDLAPRDRSLVTIAALISVGQVEQIYFHLNKAL
DNGLSEEQAAEVITHLAFYAGWPNAMSALPVAKDVFKARRK
T33-fn24-A
MTTTVQNLIADINSLTSHLHEKDFLLTWEQTPDELKQVLDVAAALKALRADNISTKVFNSGLGISVFRDNSTRTR
FSYASALNLLGLAQQDLDEGKSQIAHGETVRETANMISFCADAIGIRDDMYLGAGNAYMREVGAALDDGYKQ
GVLPQRPALVNLQCDIDHPTQSMADLAWLREHFGSLENLKGKKIAMTWAYSPSYGKPLSVPQGIIGLMTRFG
MDVTLAHPEGYDLIEEVVLIAQAAAGHSDGHYKQVTSMEEAFKDADIVYPKSWAPYKVMEERTELLRANDHE
GLKALEKQCLAQNAQHKDWHCTEEMMELTRDGEALYMHCLPADISGVSCKEGEVTEGVFEKYRIATYKEAS
WKPYIIAAMILSRKYADPGALLEQLLKENQPRVK
T33-fn24-B
MHHHHHHSGAYRYRDIVVRKQDGFTHILLSTKSSENNSLNAEVMKEVLDALHTADKDDSKLVLLSAVGSVFC
CGLDFIYFIRRLTDDRKRESSLMALAIRATAAFYAAFSKPIIVAVNGPAIGLGASILPLCDVVWANEKAWFQTPYT
TFGQSPDGCSTVMFPKIMGGASANEMLLSGRKLTAQEACGKGLVSQVFWPGTFTQEVMVRIKELASCNPVV
LEESKALVRCNMKMELNQAIERECEVLKKIWGSAQGMDSMLKYLQRKIDEF
T33-fn25-A
MTGAGRSNDTGIVLQQAMLLMAIEAQQHLLQLTVWGIKQLQTRVLGGGGRWMQWDKEISNYTNTVYQILKG
AQSQQRKNKFDTLSL
T33-fn25-B
MHHHHHHSGSIEVLKAALSEYAKDIKLNLSSITRSSVLDQEQLWGTLLASAAATRNEQVLAMIAYQALDHLSG
GQFAAALGAAAIMGMNNVFYRGRGFLEGRYDDLRPGLRMNIIANPSIPKANFELWSFAVSAINGCSHCLVAHE
HTLRTVGVDREAIFEALKAAAIVSGVAQALATIEALSPS
T33-fn26-A
MAGAGQSNDSGIVQQQSNLLRAIEAQQHLLQLTVWGIKQLQTRVLGSGGAWGLWRGFIYKYTATVKRLLEES
QNQQERNEKDLLALA
T33-fn26-B
MHHHHHHSGAYRTRDIVVRKQDGFTHILLSTKSSENNSLNPEVMRLVISSLSKAATGDSKLVLLSAVGSVFCC
GLDFIYFIRRLTHSRQLTSEQMARAIRIFVNTFIQFSKPIIVAVNGPAIGLGASILPLCDVVWANEKAWFQTPYTT
FGQTPDGCSAVMFPKIMGGASANEMLLSGRKLTAQEACGKGLVSQVFWPGTFTQEVMVRIKELASCEPAAL
MATKHAVWANMKMELEQANEIECEALKVRWGSAQGMDSMLKYLQRKIDEF
T33-fn27-A
MTDAGRKLDSIIVQQQANLLRAIEAQQHLLQLTVWGIKQLQTRVLGGGGRWMQWDKEISNYTNTVYRLLEDS
RYKQAALKLALKALA
T33-fn27-B
MHHHHHHSGAYRYRDIVVRKQDGFTHILLSTKSSENNSLNAEVQLEVQSALSTAAQDDSKLVLLSAVGSVFC
CGLDFIFFIRLLTRDREETSTTQAGADAGFVATFIQFKKPIIVAVNGPAIGLGASILPLCDVVWANEKAWFQTPYT
TFGQSPDGCSTVMFPKIMGGASANEMLLSGRKLTAQEACGKGLVSQVFWPGTFTQEVMVRIKELASCNPVV
LEESKALVRCNMKMELEQANIREAQVLEKIWGKAQGMDSMLKYLQRKIDEF
T33-fn28-A
MTGAGFLNDWGIVQQQSNLLRAIEAQQHLLQLTVWGIKQLQTRVLGGGGRWMQWDKEISNYTNTVYRLLEE
SQNQQEINEHLLRHLA
T33-fn28-B

MHHHHHHSGAYRYRDIVVRKQDGFTHILLSTKSSENNSLNEQVMLYVRSALKKAATDDSKLVLLSAVGSVFC
CGLDFIYFIRRLTDDRDRESRFMAIAIREFVNTFIQFKKPIIVAVNGPAIGLGASILPLCDVVWANEKAWFQTPYT
TFGQSPDGCSTVMFPKIMGGASANEMLLSGRKLTAQEACGKGLVSQVFWPGTFTQEVMVRIKELASCNPVV
LEESKALVRCNMKMELEQANERECEVLKKIWRSKQGMDSMLKYLQRKIDEF
T33-fn29-A
MPHLTLEYTDNLPEPQIRHLLFLLNGALLSRPEIFPVGGIRARAYRLSEYALADGSEPSDAFVHLRLQIGAGRS
DEQKKKTGDILFLILVAHFRAEFSQRGLMLSAEISEFSDKGTWKKNNIHARYRK
T33-fn29-B
MHHHHHHSGDDPRLLSLFSAQREEDADIVIIGFPYDEGCVRNGGRAGAKKGPAAFRFFLQRLGSVNNLELNV
DASHLKLYDAGDITASTLEEAHEKLESKVFTVLARGAFPFVIGGGNDQSAPNGRAMLRAFPGDVGVINVDSHL
DVRPPLSDGRVHSGTPFRQLLEESSFDGQRFVEFACQGSQCGALHAQYVRDHQGILMWLSEVRKLGAYQAL
LIAFALTGSNTFFSFDVDSLKSSDMPGVSCPAAVGLSAQEAFDMCFLAGSDEQVMMMDMSELNPLVEEYRSP
RVAVYMFYHFVLGFALRSKPKAEN
T33-fn30-A
MHHHHHHSGEQLPQCETLILEKQGPTLVITINRPDVRNAMSLQMVAELSTIFSEIENDISIRAAVLRGAGGHFCA
GGDIEDMLEARAQKAGEGRDDPFYKLNRAFGQMIQQVNESSKVVIAITEGAVMGGGFGLACVSDLAIAGPTA
KFGMPETTLGVIPAQIAPFVVERIGLTQARRLALLGLRIDATEACKLGIVHQVAESEEQLSDMLNQALERVRLCA
PDATAETKALLHRVGHEAMAGLLDDAAEKFAAAIRGPEGAEGRMASLQDREPKWAELPNQ
T33-fn30-B
MTQTAPAAVAYSVNHAGVAAIVLDRPEASNALDRTMKTELLQALLAAGGDPAVRAVVMSAAGKNFCVGQDLA
EHVEALRDDPANAMKTVEEHYNKVLEALDAIKVPVVVAINGACVGAGLGLALGADIRIAGQRAKFGTAFTGIGL
AADSALSASLPRLIGASRATAMFLLGDTIDAPTAHTWGLVHEVVDEGSPADVANSVAGRLAGGPTAAFSEVKE
LLRRNAVAPLGTVLLKETIAQLRLGSSRDHSAAVEAFLAKDKPVFVGR
T33-fn31-A
MHHHHHHGSNDVLFSNHGRVAVITLNRGDRLNAWTTPMRETIIDALERFNRDPEVAAIIMTGKGREAFSAGQ
DLSEAHDFDGERAVAWVKEWQRYYTALRSLSKPLVMALNGTAAGSAFQVALLGDIRVGHPGVRMGQPEINA
GIASTTGPWIMNAMLGMSRTIELTLTGRLMPADHCHRIGLIHVLTSEDLVFDEALLIATELAAKPPVAMRLDKQR
FREMTEPGFIDCIEAGERIQREAYDSGEPARMMEEFFSKRAK
T33-fn31-B
MSGIDTKQQNNLLSAIIAQQHLLQLTVWGIKQLQARSGGRGGWMAWDRHINNLTSIIHSHIKESLDQQEK
T33-fn32-A
MHHHHHHGSNDVLFSNHGRVAVITLNRPDRGNAWTTPMRETIIDALERFNRDPEVAAIIMTGAGNDIFSKGQD
LSEAHDFDGERAVAWVKEWQRYYTALRSLSKPLVMALNGTAAGSAFQVALLGDIRVGHEATFMGQPEINAGI
ASTTGPWIMNAMLGMSRTIELTLTGRLMEAEECHRIGLIHLLVHESQVFDMALIIATNLAAKPPVAMRLDKQRF
REMTEPGFIDCIEAGERIQREAYDSGEPARMMEEFFSKRAK
T33-fn32-B
MILVYSTFPNEEKALEIGRKLLEMRLIACFNAFEIRSGYWKDGRIVQDKEWAAIFKTTEEKEHDLYEALRLLHPY
EFPAIFTLKVENVLEEYMALLRASVS
T33-fn33-A
MHHHHHHSGFQSMSNDVLFSNHGRVAVITLNRPERLNAWTTPMRETIIDALERFNRDPEVAAIIMTGAGQDAF
SAGQDLSEAHDFDGERAVAWVKEWQRYYTALRSLSKPLVMALNGTAAGSAFQVALLGDIRVGHKHVRMGQ
PEINAGIASTTGPWIMNAMLGMSRTIELTLTGRIMPAKECHRIGLIHYLTHESTVFDVALLIAEILARKPPVAMRL
DKQRFREMTEPGFIDCIEAGERIQREAYDSGEPARMMEEFFSKRAK
T33-fn33-B
MPHIRVRGAEKEKVRDFTAGLADILGRAASDTASAFTFEYVETTFFFDGKEDDGLVFIEVLWFDRDSETRATIA
LLFTLKWKRITDKIVTIVFNPLIENMYYVDGKRF
T33-fn34-A

MHHHHHHSGPSSAIATLAPVAGLDVTLSDGVFSVTINRPDSLNSLTVPVITGIADAMEYASTDPEVKVVRIGGA
GRGFSSGAGISADDVSDGGGVPPDTIILEIERLVRAIAALPHPVVAVVQGPAAGVGVSIALACDVVLASENAFF
MLAFTKIGLMPDGGASALVAAAVGRIRAMQMALLPERLPAAEALAWGLVTAVYPADEFEAEVDKVIARLLSGP
AVAFAKTKLAINAATLTELSPALQRESLGQSVLLKSPDFVEGATAFQQRRTPNFTDR
T33-fn34-B
MIIVYTTFPDWESAEFVVKQLLLARMIACANLREHRAFYWWSNSIEEDKEVGAILKTRESLWRDLKEAIKQLHP
YDVPAIIRIDVDDVNNGYEEWLIEETQK
T33-fn35-A
MHHHHHHSGTQTAPAAVAYSVNHAGVAAIVLDRPEASNALDRTMKTELAAALKKAAGDESVRAVVMSAAGK
NFCVGQDFTEHAVALARDPRHAMDTVREHYNPVLEALDAIKVPVVVAINGACVGAGLGLALGADIRIAGQRAK
FGTAFTGIGLAADSALSASLPRLIGASRATAMFLLGDTIDAPTAHTWGLVHEVVDEGSPADVANSVAGRLAGG
PTAAFSEVKELLRRNAVAPLGDVLEREASAQQRLGASRDHSAAVKAFLAKDKPVFVGR
T33-fn35-B
MQWQTKLPLIAILRGITPDEALAHVGAVIDAGFDAVEIPLNSPQWEQSIPAIVDAYGDKALIGAGTVLKPEQVKA
LADMGCQLIVTPNIHKEVIVAAVAFFMTVCPGCATATEAFTALEAGAQALKIFPSSAFGPQYIKALKAVLPSDIAV
FAVGGVTPENLAQWIDAGCAGAGLGSDLYRAGQSVERTAQQAAAFVKAYREAVQL
T33-fn36-A
MIVDYSVNHAGVAAIVLRDAKNSNALDDGAKTELLHALLKAGGDPAVRAVVMSAAGKNFCVGQDLREHWIAT
AKDPAHAMDTVREHYNPVLEALDAIKVPVVVAINGACVGAGLGLALGADIRIAGQRAKFGTAFTGIGLAADSAL
SASLPRLIGASRATAMFLLGDTIDAPTAHTWGLVHEVVDEGSPADVANSVAGRLAGGPTAAFSEVKELLRRNA
VAPLGDVLEREASAQQRLGASKDHRAALLAFMNKDKPVFVGR
T33-fn36-B
MHHHHHHSGSAVQPFIRTNIGSTLRIIEEPQRDVYWIHMHADLAINPGRACFSTRLVDDITGYQTNLGQRLNTA
GVLAPHVVLASDSEVFNLGGDLALFCQLIREGDRARLLDYAQRCVRGVHAFHVGLGARAHSIALVQGAALGG
GFEAALSCHTIIAELAGSYGLPEVQNDLFPGMGAYSFMCQRISAHLAQKIMLWGNLFSALQLLGMGLVDAVVS
EGSGVDMVEFVIFISKRTPHAWAAMQQVREMTTAVPLEEMMRITEIWVDTAMQLGEKSLRRMDELVKADSRR
SGLDAG
T33-fn37-A
MHHHHHHSGMTQTAPAAVAYSVNHAGVAAIVLDRPEASNALDRTMKTELLQALLAAGGDPAVRAVVMSAAG
KNFCVGQDLAEHVEALREDPSNAMATVREHYNPVLEALDAISVPVVVAINGACVGAGLGLALGADIRIAGQRA
KFGTAFTGIGLAADSALSASLPRLIGASRATAMFLLGDTIDAPTAHTWGLVHEVVDEGSPADVANSVAGRLAG
GPTAAFTFVKYALRMNAVAPLGVVLDIEATFQQFLGASRDHSAAVEAFLAKDKPVFVGR
T33-fn37-B
MAGAGQSNDSGIVQQQSNLLQAIQRQLHLLELTVKGIKQLQTRVLGGGGLWTAIDLQISFMTEAVKRLLREAQ
EQQDRNEKDLLALA
T33-fn38-A
MHHHHHHSGTQTAPAAVAYSVNHAGVAAIVLDRPEASNALDRTMKYELLKALLLAAANLDVRAVVMSAAGKN
FCVGQDRDEHIEALRDDPKNAMDTVREHYNPVLEALDAIKVPVVVAINGACVGAGLGLALGADIRIAGQRAKF
GTAFTGIGLAADSALSASLPRLIGASRATAMFLLGDTIDAPTAHTWGLVHEVVDEGSPADVANSVAGRLAGGP
TAAFSEVKELLRRNAVAPLGDVLEREASAQQRLGASRDHSLAVKAFMADAKPIFVGR
T33-fn38-B
MTSTAVEITVKNAADIAIIGGSGLYQMQALTNKRSVRIATPYALPSDDIVLGELNGVTVAFLTRHGQGHRLTPSE
VPYRANIYALKSLGVRYIVSVSAVGSLQETLKPLDMVIPDQMIDMTKQRVSTFFGDGAVAHVSMADPLCPEVA
DILIRAYDNADIADGQCHAKATYVCIEGPQFSTRAESHWYRQMQADIIGMTNMPEAKLAREASIAYATLALVTD
FDCWHPNEQAVSADYAIQNLMKNADNAQQVIKQAVALIASEQPKSIAHTALTQALVTPVEAMSEETKLRLFALL
P
T33-fn39-A

MTQTAPAAVAYSVNHAGVAAIVLDSPRNSNALDDEMKTELLQALLAAGGDPAVRAVVMSAAGKNFCVGQDL
FAHFAELRRDPAHAMDTVREHYNPVLEALDAIKVPVVVAINGACVGAGLGLALGADIRIAGQRAKFGTAFTGIG
LAADSALSASLPRLIGASRATAMFLLGDTIDAPTAHTWGLVHEVVDEGSPADVANSVAGRLAGGPTAAFSEVK
ELLRRNAVAPLGDVLEREASAQQRLGASRDHSAAVEAFLAKDKPVFVGR
T33-fn39-B
MHHHHHHSGMSLRLERDGAVARLLIDRADRRNAFSLDMWLRLPELLAEASGDDALRVLVVKSANGGAFCAG
ADIAELLANKDDGAFHDANQMAILRAQLELARFRLPTVAMVEGDCIGGGCGIALACDMRIAAPAARFGITPAKL
GLVYPLHDVKLLVDLVGPGQARRLMFTGGLIDANEAHRIGLVELLGESEDALVGQLATVSSFSTQAIKSFVRRV
LDGQVAMDADAAHVLASAYEGADFREGTGAFLEKRPPVF
T33-fn40-A
MSDLRLERDGAVARLLIDRPQNNNAFDTQMWQNLPVLLADASGDDALRVLVVKSANGGAFCAGADEHLLLT
RMLDDDWHAENQQAINRAQYELARFRLPTVAMVEGDCIGGGCGIALACDMRIAAPAARFGITPAKLGLVYPLH
DVKLLVDLVGPGQARRLMFTGGLIDANEAHRIGLVELLGESEDALVGQLATVSSFSTQAIKSFVRRVLDGQVA
DDADSLRVFASAFKQKDFMEGQLAFAQNRPPVF
T33-fn40-B
MHHHHHHSGMYETIRYEVKGQVAWLTLNRPDQLNAFTEQMNAEVTKALKQAGADPNVRCVVITGAGEAFCA
GEDLSGVTEEMDHGDVLRSRYAPMMKALHHLEKPVVAAVNGKAAGAGMSLALACDFRLLSETASFAPAFISV
GLVPDAGHLYYLPRLVGRAKALELAVLGVRVTALQAAKLGLATAVIPKHLWELAVKAYASALSNMPTKAIGLIK
RLLRESEETTFDRYLEREAECQRIAGLTSDHREGVKARNESRKPLFQGN
T33-fn41-A
MHHHHHHSGMSLRLERDGAVARLLIDRADRRNAFSLDMWQRLPELLKEASGDDALRVLVVKSANGGAFCAG
ADIAELLANKDDAAFHAANQDAINYAQYELARFRLPTVAMVEGDCIGGGCGIALACDMRIAAPAARFGITPAKL
GLVYPLHDVKLLVDLVGPGQARRLMFTGGLIDANEAHRIGLVELLGESEDALVGQLATVSSFSTQAIKSFVRRV
LDGQVADDKQSLLYFAAAYHHADFREGTGAFLEKRPPVF
T33-fn41-B
MNDFLNSTSTVPEFVGASKIGDTIGMVIPNVDQQLLDKLHVTKQYETLGILSDRTGAGPQIMAMDEGIKATNME
CIDVEWPRDTKGGGGHGCLIIIGGDDEKDARQAIRVALDNAARTFGDVYNAKAGHLELQFTARAAGAAHLGLG
AVEGKAFGLICGCPSGIGVVMGDKALKTDGVEPLNFTSPSHGTSFSNEGCLTITSAAAAVLTAVLAGRRVGLK
LLSQFGEEPKNDFQSYAK

T33-ml1-A
MYPVDLHMHTIKNDASFSTLSDYIARAKEKGIKLIAITDHGPSHPIAPHPEYYVRMKELPDVVDGVGVLRGVEA
NILDTNGNIDVTPEMEKSLDLILAGLYESVYPPQSRAENTKALINAIASGKVHVISHPADPRYPVDYRALAGAAA
VAGVALEITEHAFGEEFPGAEPRARELARAVKEAGGYVALGSDAHHAWHLGRFEHAERVLREVGFPEERVLN
RSPEKLLAFLESRGVPKKPAFADLGGSHHWGGHHHHHH
T33-ml1-B
MSISYRKLDIALSADGEEVLVEGFVLPTKFFENVIVTTMLNAAGTDEENINALLADVHAAGLDVSNYGKASEIYA
KGDPEKRAEAEARRAEAEARRAELAAELSTPEAQAEAKKEKVLEAAELAARFGPAGVKAGL
T33-ml2-A
MHHHHHHGGSHHWGGKLLLAATGSPAARFFGELAKQFVPHFEVRAVLTEGALEFVDLSSLPAEVPVYTDED
MRAAFKKEGDVILHIELADWADVLLIAPASINTIAEIASGLAPNLLLRIFAGWDLSKPVFIAPAMSQREYDNPATK
ENLKKLEERGVIIIPPVKGRGADGSVGNGVMAPPEKIADAVLGYLEARKALKKVVTS
T33-ml2-B
MNMAETYYEIGKKFEKTGAYDAAIHAYLAALAEDPNNAEAWYNLGKAYEKLGKYKEAIEAYKKALELDPKNAE
AWYNLGKAYEKLGDYKKALEEYLKSLELDPFNEEAKKNAKEAGKKGVLE
T33-ml3-A

MKLPNKVSLVAGTAEGATPENALHGARLNAGIGDVNLVPVSGIAPAGAEIVPLPELPPGALLPTAEASIVSDVP
GKTIAAAVAVGIPKDPSLPGIIATYAGEMSAEEARRRVELIVIEQFLQRGWELESIHSVAVEHTVKRLGAALAAA
VLWYKGGSHHWGGHHHHHH
T33-ml3-B
MTAEGETAVAIALLGDEDPHVRAEAAKKLGKIGDKEAVKPLIEALGDEDPAVRAAAALALGKIGDKEAVPPLIGA
LLDEDPAVRVAAALALGKIGDKEAVPALILALLDEDEAVRVAAAVALGKIGDKEAVEPLVKALEKEEGLVRKAAA
IALEKIGGEEVKKAAEELAKKGEGEARKAAEEYLKKHKLE
T33-ml4-A
MLDEIAFADARILTPFTEADIERLLDALELEPGTRVVDLGCGTGYFLVLGAERKGITATGIDISELAIEKARELARE
RGVEDRVEFIHGDVSTYVAEEKVDVAACIGAEAYFGGIEGALKALEKSLKEGGIILLGVPYWRTRPATEAEARA
CGFDSIDDLDTLAETVAKLEALGYRVIQIVLADEHGMEELYYGRLVQLDRWLREHPDHPFAPALEAELATLAAR
YEKYVRRHLGYGVFALRKRLEGGSHHWGGHHHHHH
T33-ml4-B
MIVVLITVPSEEVARKIARAAVEGGLAAEVLIIPALTLYYRENGKVVERPVYLLLVLSTESKFPALLALVKALHPEK
VPLIVALPVVDGNPEYLRWVKLNTG
T33-ml5-A
MPMVIFECSDNIREEAKFEELFARLNPALASTGLFPLEEIVGRVHWVDTWQFADGQHDYAFVHVTIDVPAGLS
EEDRLLVLNGVFALLLGHLEPLMKEHLLYLSLELRVLPATLSRRWNNAIELFKGGSHHWGGHHHHHH
T33-ml5-B
MSVKIDVIRVEIPEGTWVIIGQSHSRSIVFDLSQTLQSASGRLRFGIAYCEASGKRLILHDGNDPALVELAKETAL
KIGAGHTFVIYIRNGRPEDILNRIKNIESVVRIFAATANPLQVLVAETDQGRGVIGVVDGYTPLGVATAEDRAALA
AALRAEGYKR
T33-ml6-A
MTAVFAAIVGFLDKKIEELKKIQKHKTLPKMSGGWELELNGTEAKLVREVDGYKVTVTFNINNSIPPTFDGEEE
PSQGQPVEEQPPELTSTPNFVVEIVKASQPDTALVFDCYYPEDEVGQEEEEEKPLFEIKEVSFQSTGESEWK
DTNYTLNTDNLPEDLLLAFYALLAALGVDNDFAKELIELSTALEHQERITFFEQLRDFIA
T33-ml6-B
MNLAEKMYEAGKYFAAQGNYELAIIAYTLALLKDPNNAEAWYNLGKAYAALGKYEEAIEAYKKALALDPNNAE
AWYNLGGAYAGLGKYEEAIEYLEKALALDPNNELAKLMLKFAKLQLELEGGSHHWGGHHHHHH
T33-ml7-A
MHHHHHHGGSHHWGGMLFSFLHEEKKLGKIIVVDKGSGPEHVRSQLKTCGDYIDYVRFAGKTAAQMPAEVV
KEKIAIYHEKGIKVFPGASLFEEAVKKGKEDEFLAECKAAGFDAVEIGNLNINLSDEELEALIKKAKAAGFEVFTV
VGRADPKVDKLLSVSDIVRRIRRFLEAGADYVIIYGGSTGKGKGLYDENGNLKEEDLDYIKENVPMEKIIFEAPL
EKQQKQLIEKFGPSVNIAEISFSDVITVANLRSGLRGDTFGKV
T33-ml7-B
MKERQISLLETLLSLYIDLLEVMADMAGKSGKYVLLDVREDLKFVEKNKIPGAIWLPVSLLEERIDELDPSKTYV
VYDYKGHSTNSYRALLILLKAGFEAYILSGDPKLLLG
T33-ml8-A
MLAFEFLHEEKELGKIIVVDTGTSPHHLKGQLETVGDFIDYVKFAGMTAAVMPKKVVCEKILIYHKHGIKVMPG
GTLFEKAVSKGKEKEFLLECRELGFDAIEISDLNIDLSDEELKKLIKMAKEEGFEVFTKVGRADKERDAKLTVED
IIAKINFYLEAGADYVIIYGGASGKGIGLYDENGKLKKEWLDEIKKNVDMSKIIFEAPLPEQQKELLDKFGPSANL
AEISLHDVARLAEMRYGLRGDTFGKV
T33-ml8-B
MHHHHHHGGSHHWGGGMAQAQTQGQEEEQKKKIVIIIRHGPEEPIYCVTPLRLAVVAAEQGYETTIVFTELGP
ELLNKLYWIEEMAKGGNPVTKYLLKAREKGVKIYVCEWSLEEICKLKKEDIIPGVEIIDDKDIIKLMLEADVVIFF
T33-ml9-A
MHHHHHHGGSHHWGGMKAFEFLYEDFQRGLTVVLDKGLPPKFVEDYLKVCGDYIDFVKFGWGTSAVIDRDV
VKEKINYYKDWGIKVYPGGTLTEYAYSKGKFDEFLNECEKLGFEAVEISDASIDFSMKEMIDMIRKAKANGFMV

LTLVGRKDPAKDAELTIVERVIRITAYLDAGADYVIIYGRESGKGKGLFDKEGKLKLDDLDILASSVDMSKVIFEA
PQKSQQVALILKFGSSVNLANIAFDEVISLETLRRGLRGDTFGKV
T33-ml9-B
MSKTTVIYPGSFDPIHKGHVDLIERASKMFPRVVVAVVKGHHKKHTFSIVERLLLVEAAVGHLPNVEVRAVDGL
LVNVFKELKATAVLRGLRAVSDFEYEFQLANMNRQLDPHFEAVFLTPSEQYSFISSTLIQKLAANGGDISQFVP
PVVVAAFKALKGKGW
T33-ml10-A
MPLVVLVTVPSEEEARRIARALVERRLAAQVNIVPGLTSIYRRDGEVVEDQELLLLVFTTELRFPLLRELVRSLH
PEATPMIVALPVVDGNTDYLLWLLENTG
T33-ml10-B
MPFEKALYFLTYLSYTIDIAELSILIKKGDKSIIIVDVRDAEAYKECHIPTAISIPGNKINEESTKDLPKDKTIITYCWG
PACNGATRASRKFAELGFDVKRLIGGIEYWRKENGEVEGTLGAKADLFWNMKKESLEGGSHHWGGHHHHH
H
T33-ml11-A
MFSNKRVLVEKEGEAGIAVMKFKNPPVNSLSLEFLTEFVISLEKLENDKSIRGVILTSERPGIFSAGLDLMEMYG
RNPAHYAEYWKAVQELWLRLYLSNLTLISAINGASPAGGCLMALTCDYRIMADDDGYTIGLNESLLGIVAPFWL
KDNYVNTIGHRAAERALQLGTLFPPAEALKVGLVDAVVPPEAVLAAAKGTLAEWFQIPDHSRQLTKSMMRKAT
ADNLIKQREADIQNFTSFISRDSIQKSLHVYLEKLKQKKG
T33-ml11-B
MHHHHHHGGSHHWGGSSGLVPAGSHMRLTPHEQERLLLSYAAELARRRRARGLRLNHPEAIAVIADHILEGA
RDGRSPGELAAAGQTVLGRDDVVGGVPEMLAEVQVEATFPRGTMTVTVERPIA
T33-ml12-A
MMSGWFPVKTTEELEVIDITPLVEAALKGAGLKNGLVLVYVPDVDAAIIVNTADPELLEDIVRHLRTLCDPEGD
WAYNKVEPNAHAYLGTALVGNSVVIPVRNGKLDLGKEQKVLFIDMDGPDTYTVKLMALEE
T33-ml12-B
MIKKPEFGLMQPPKKRVRQELSSVAEETIEAAFDFFDVDGDGKINKEELKKALHALGFAVNDMQIEALMAAYD
KDGDGYINKEEFKEIVELLRKNGGSHHWGGHHHHHH
T33-ml13-A
MNAVVQRTELGNGVVQITMKDESSRNGFSPSIVAGLKAALDAVIDDSSVKVVILTGYGNYFSSGASKEYLLALT
KGEVGVLNLVPLILDCPVPVIAAMQGHSFGGGLLLGLACDFVVFSQESVYATNESKYGFTPYAAARLILRRKLG
SELAQEMLYTGENYRGKELAERGIPFPVVSRQDVLNYAQQLGQKIAKAPRLTLILLNIDARADLRAAYPAALRR
ELGLFSLTFSQPEIPERIQQEF
T33-ml13-B
MRRGLLPNDVWQADICEYKYKKKYKYCLHIVVDTFSGAMSVSCKKKKTPLETIEALLQAISLLGRPKKIISDHDPA
FRHGLTKAFCLSSGIELESYTPGDPSSSALVDAACKELKALLDRYLTENPELPLDNAINLALWEHNQLKVVEPY
GKTPWQLHHSGGSHHWGGHHHHHH
T33-ml14-A
MHHHHHHGGSHHWGGSELTVNVINGPNLRMLGFREPAVYGGTTFSELVELIEREAAELGLKAVVRQSDSEA
QLLKWIHLAALMAEPVILNAGGLTHTSVALRDACAELSAPLIEVHISNVHAREEFRRHSYLSPIATGVIVGLGIQG
YLLALRYLAEHVGT
T33-ml14-B
MPMFIVNTNVPRASVPDGFLSLLTRLLALLTGKPEKYIAVHVVPDQLMAFGGSSEPCALCSLHSIGKIGHEQNR
SYSKLLCTVLAQRLRISPDRVYINYYDMNPENVGWNNSTF
T33-ml15-A
MSLKDKKILIVEDSLEQAITIGLILVKYGYEVIIAGTGEQAVEYVSGGEYPDLILMDIELGEGMDGVQTALAIQQIS
ELPVVFLTAHTEPAVVEKIRSVTAYGYVMKSATEQVLITIVEMALRLYEANVHANEG
T33-ml15-B

MHHHHHHGGSHHWGGGRSLVVIVNDRTAHGDQDKSGPLVVGLLRAAGFVVDGVVVVENDLSEIQNAVNTA
VIGGVDLVVTVGGTGVTPRDVAPEATQPLLDRELLGIAEAIRSSGLAAGVTEAGLSRGVAGISGSTLVVNIAGS
QEAVLVGLKTLLPMAIQIIEQLSSLEI
T33-ml16-A
MAEVSIKTKIKAKHRLYSKNLSEEENKMLFGSAAKKGGHEHNYTITVTVKGEIDPTTGLVINGTDLRIWIEKAIILP
LDNKNLNEDVPYFKTNVPTTENIAKYIKENLEKVLPKGLLSKVVVEETEEHKVTIKGEGGSHHWGGHHHHHH
T33-ml16-B
MPAIILTTTPTEADARALAEGLLEKRLIAEAIITPNVTRIYLENGEIKSEKVVRMELYTVEEKVEAAMTYIEAHHPD
PIPPIIVIKPDKVSPKYKKWILEQTAL
T33-ml17-A
MAPTMTEFVGTAGGDTVGLVIANVDSLLHKHLGLDNTCRSIGIISARVGAPAQMMAADVAVQTTNTEVATIELP
RDTKGGAGHGIFIVLKAADVSDARRAVEIALAMTDEYLGDVYLCDAGHLEVQFTARASLIFEKAFGAPSGQAF
GIMHAAPAGVGMIVADTALKTADVKLITYGSPTNGVLSYTNEILITISGDERAVLKSLDAARKAGLSILKDMGEK
PVSMSEPTF
T33-ml17-B
MPLIRIDLTSTRSRLQRQLIAQAVHDALVEVLAIPARDRFQILTAHPISDIIAEDAGLGFTRSPDVVIIHVFTQAGR
TIETKQRVFAAITESLAPIGVAGSDVFIAITENAPHDWSFGFGSAQYVTGELAIPATGAAGGSHHWGGHHHHH
H
T33-ml18-A
MKILIVVTHGPEDLDRTYAPLFLAVVAAERGYKTSVFFMIKGPLLLNRDYIAKVALEGGNPYLEYLYKAKQLGVE
IYVCVQSLRDMCHLKEEDIIGGVKLVGGSTLIDLTLEADRTLFF
T33-ml18-B
MNLAEKMYKAGNAMYRKGQYTIAIIAYSLALLLDPKNAEAWYNLGNAYYAKGEYDDAIKAYEKALMLDPNNAE
AWYNLGNAYYAKGDYESAILAYQLALKLDPNNAEAKQNLANAKQKLALEGGSHHWGGHHHHHH
T33-ml19-A
MPMVTIRTNLPASEVPADFAAELTALLSKTLGVPADRIAVEVLPGVDLTFGGSREPVALITVESIGNLTPEQTNL
LTLQLTLLLQLRLGLPEDRVLILFHDLPASQVGRDGRTEAAA
T33-ml19-B
MHHHHHHGGSHHWGGSPSDPPRPALLMLELRSYALGLAVADAALRAAPVRLLLARPVEPGKALILLTGEEEA
CRAALEAALRVAREGSGNLLDSVFIPAIHPQLLPFLLEEVAAPPLADPDEAVLVAEVRTPAAAIRAANAALEAAP
VRLTRMRLAEHIGGKAYFTLTGRREDVLRAAQVIAEVAGEDLIDLRLIPRPHAALRGREFF
T33-ml20-A
MHHHHHHGGSHHWGGGMKVTFLGAAVVLIEGKKNIIIDPFISGNPVCPVKLEGLPKIDYILVTHGHGDHLGDAV
EIAKKNDATVISNYEICHYLGKKGVKTHAMHIGGSYLFDFGRVKMTPAVHGSGILDGDSMIYGGNPSGFLIEID
GKKIYHAGDTGLTREMELLAEENVDVAFLPIGGNFVMDVKDAIKAAKMIKPKKVVPMHYGTWELIFADVEAFKA
GVEAIGVECVILEPGESLEL
T33-ml20-B
MPVITVNTNVAEKSIPVFFQAALTNMMSKLLDVGKERMFVDLRSGANIMMGGDRNPCVFATVECIGRLNPGS
CALMAQEMEKMFIEHLNVRRERIVIRFIPVPAEFCSFNGKLHDVKEERDEYLE
T33-ml21-A
MTVPEFVGASEIGDTIGMVIPRVDQQLLDKLHVTKQYKTLGILSDRTGAGPQIMAMDEGIKATNMECIDVEWPR
DTKGGGGHGCLIIIGGDDPADARQAIRVALENLPRTFAGVFNAKAGHLEFQWTPRAAGAAHLGLGAVEGKAF
GLICGCPSGIGVVMGDKALKVAGVEPLNFTSPSHGTSFSNEGCLTITGDPRAVLAAVMAGAEVGLKLLSQFGE
EPVD
T33-ml21-B
MPLIRIDLTSDRSRFQRLAIAEAVHLALVEVLAIPERDRFQILTAHDPLDIIAEDAGLGFTRSPSVVIIHVFTQAGR
TIETKQRVFAAITEALAGIGVAGSDVFIAITENAPHDWSFGFGSAQYVTGELAIPATGAAGGSHHWGGHHHHH
H

T33-ml22-A
MHHHHHHGGSHHWGTVPEFVGASEIGDTIGLVIPRVDQQLLDKLHVTKQYKTLGIISDRTGAGPQIMAADEGI
KATNVECIDVEWPRDTKGGGGHGCLIILGGEDPEDMRHAVRVALAELPRTFARVFYNKAGYAVFQYTDAAAG
AAHLGLGAVEGKPFGLIAGCPSGIGVVAADEALKVEGVEPLNFTSPSHGTSFSNEGCLTITGDPEAVRLAVER
GEAVAIRLLKTFGEEPKNDFPSYIK
T33-ml22-B
MTDPMKVILYIAMLELEKYIMRAAAAYALGKLGDLRAVPPLIKALKDEDAIVRAAAADALGKIGDLKAVPPLIKAL
KDEDGAVRVSAAVALGKIGDLKAVEPLIKALKDEDAVVRVAAAIALGKIGDLRAVEPLIKALIDEKGKVQEAAAL
ALGAIGGERVREAMEKLAEEGKGRARLYAVKYLGEHDAE
T33-ml23-A
MAIIETTTPTEEEAKAIAKKLLENRLIAEAIITPALTKIYRENGEIKSETVTRVTLYTEEENVPKAVTYIKAIHPDPIPP
IIVITPTDANPAYKGWVAFET
T33-ml23-B
MHHHHHHGGSHHWGGDPERPALGILELSSYARGVKVADAALKAAPVKLLKCEPVEPGRALIMLLGEPEDVAK
AMIAALDVAGLGSGNLIDYALIPEIHPQLLPFLKEYKKSEPIKDPNKAIIVAEVSTVAAAIEAADVALRLANVELTS
MRLAEHIGGRASFTLIGDKEDVEKAARAIRGVAGERLLDLEIIEKPVEALIGNEFF
T33-ml24-A
MALIYTTTPTYEDAMNIAKKLLENKLIAYALIFSNITSVYVEEDEIHNNTECAVIMATVEEKVLKATLYIEAIHPKDM
PPIIVIVPADVSPRFQGWVYAKT
T33-ml24-B
MHHHHHHGGSHHWGGPARPALGVLLLTSIARGITVTDAALKAAPVRLLMSRPVCSGKHLNIFTGRPEEVLTA
MFAALETAGLGSGKLLDYAFIPALHPQLLRFLDAPVVADAWEEDTEAVAVVETTTPCAAIESADVALKLAPVRL
RDLRLAIGIAGKAYFTLAGREEDVRRAAKAVKGTAGDKLIELEFIARPVDELRGRLFF
T33-ml25-A
MDGEDALAAATAAEVAALLAILEAGLAALKALGFPLPDETGLDNRFLKALADWLRTEKALLTLREEALLRLLRLL
VERSA
T33-ml25-B
MHHHHHHGGSHHWGGEPDRPALGVLLLASIALGRAVADAALKAAPSLLLMSRPVCPGKHLIMMRGQVAEVE
TAMAAALATAGAGSGNLLDSAELPYAHEQLWRFLDAPVVADAWEEDTLAVLVVETATPCAAIRAADAALKTAP
VTLRDMRLAIGIAGKAWFTLAGDPLAVLRAAVTVVAVAGDRLLRLEFIERPVDELRGRLFF
T33-ml26-A
MPVLTIATNVPSEAVPEGALLGLTLMLSELLGVPPEEIAVQITPDQRMVFGGSSEPCAICELKSIGKINAEKNKE
LSAALTEFLERALGIPPERVLILFHNVKKENWGRNGGVFAGGSHHWGGHHHHHH
T33-ml26-B
MRMKYKVVIVTGVPGVGKSTVLKELEKIAKEKGIKIAVFDFFDYMLELAKKDGLVTKKDDIPFLPLDVLKKLMKE
AAKKIVEEAEKLLDEDGILLIDTQAVIKTNHGYVPGLPKFVMDVLKPDIIAVVEASPLDITRRMLADTSRRLAYMG
GGPGVAELMETERAAAIAAAIHTGAAVLFVRNAPGMERRAAERLLKAILNL
T33-ml27-A
MHHHHHHGGSHHWGPAGEPDRPALGVLELASIALGVAVADAALKAAPVLLLMARPVCSGKFLLVLRGEPEAV
RAAMEAALRTAGVGSGNLLDFLFLPAVHEQLLRFLDAPVVADALEDPDLALLVAETATPCAAIAAADAALKTAP
VRLVDLRLAIGIAGKAWFVLAGAEEDVLRAALVVARVAGDRLLDLRFLPAPHDELRGRLFF
T33-ml27-B
MIKLSADKETVLVHGQELSTKFFLEVVVQTQLLAAGTNTALATQILALVLAAGLPVDDYGAYSRAFATGDPALR
AAAERVRAKAEAEREAMAAIHATPEEIAKAVAERKAREEALIKRFGNKGAAFGL
T33-ml28-A
MHHHHHHGGSHHWGGDPARPALGVLELKSYALGVAVADAALRAAPVELLKCEPVEPGKALIMIRGEPEAVA
RAMAAALETAKAGSGNLIDHAFIGRIHPALLPFLLEETAAPPIEDPDEAVLVVETKTVAAAIEAADAALDVAPVRL
LRMRLSEHIGGKAYFVLAGDEEAVRKAARAVRAVAGEKLIDLRIIPRPHEALRGRLFF

T33-ml28-B
MPIALTVVPPEEAEPLARELVEAGLAAEVLLVPVRRIYREKGKVREEEVTLLLILVSREGVPALRAWIEARHPDD
IPLFIVLAVDEEASNKRYLGYIAAETHLYSA
T33-ml29-A
MAPARPALLVLELSSYALGVEVADAALKAAPVELLLCRPIEPGKALIMLTGEPEAVEAAMKAALETAQEGSGNL
IASLFIPAIHPALLPFLLEPVRAPPLADPDEALLVAETSTVPAAIRAADEALRAAPVTLVRMDLAEHIGGKASFVLT
GELEDVVRAARVVVEVAGEDLIDLRIIPRPVAALRGRLFF
T33-ml29-B
MPMLIVYVPEGFSKAQKRQLLLLLHLAVVEALGVPLENVSIILTTVEPEDVLLGGKIGRPLAVVLVYILEGLSPEQ
KAALIKALTEAVAKALGMDPENVSVIIVEVKPENFGVGNGKSAKEAGGSHHWGGHHHHHH
T33-ml30-A
MGPDEPERPALLILELKSYARGVRVADAALKAAPVRLLKCKIVEPGKALIMLTGRPEDVEKAYKAALTVANKGS
GNLIDSVFIPAIHPALLPFLLEETPAPPLEDPDRALLFVEVKTVAAAIRAADAALRAAPVELVRMRLSEHIGGKAV
FALVGDPADVLRAAAVVAEVAGDQLLDIAIIPRPHPALLGREFF
T33-ml30-B
MPMLVVYVPEGYSEAQKRALLFRLAAAVVEATGTPLENVRIILTTYAPADVLLGGAIGVPLVVILVYLLEGLSPE
QKAALVKALTAAAAEALGVDPENIRVILVPVPPENFGVGNGKTAAEAGGSHHWGGHHHHHH
T33-ml31-A
MHHHHHHGGSHHWGGPAGEPDRPALGVLLLKSYARGVAVADAALKAAPSLLLMNRPVCPGKHLLMMRGQV
AEVEEAMRAALEEAGEGSGQLLASAFIPYAHEQLWRFLDAPVVADAIEEPDLAVAVVETKTPCAAIRAADAAL
KAAPVVLRDMRLAIGINGKARFTLEGKLVDVLEAAAVVIEVAGDDLISLSIIPRPHDELRGRWFF
T33-ml31-B
MADFHEQMATMFKNLAKILKAKNAAEVKDALKEMRKAALAAHKEVPPSLKDKPLNSQEMIEFHDEMLELAWAI
HDAAHLAKEGKIEEAKKKAEEILKMVSRLVSLY
T33-ml32-A
MHHHHHHGGSHHWGGMSISYRKLDIALSADKKTVLVFGQELSTKYFTEIVVTTMLNSTGSDMANSNRILNDIH
AAGLDAGDYGKYSRWWAQSNAQERQEAERRRKEAKAHQERLRAEKATVAAQLAAAAARLAEMRRLRERFG
EAGIAAGL
T33-ml32-B
MNLAEKMYNAGQAMYRKGQYTIAIIAYTLSLLKDPKNAEAWYNLGQAYYKKGQYLDAIESYLKALTLDSSNAE
AWYNLGQAYYKLGHYEEAIEAYEKALALDPNNAEAKQNLGNAKQKLGLE
T33-ml33-A
MHHHHHHGGSHHWGGMSISYRKLDIALSADGREVLVFGQVLKTTFFKNIVVTTMLNSTGSDMANSNRILNDIH
AAGLDAGDYGKYSRWWAQSNAQERQEAERRRKEAKAHRAARRAALSTPEALAAATAEIEAERAALGARFGP
AGLDAGL
T33-ml33-B
MRMFFKVVVVTGVPGVGKTTVIKELQGLAEKEGIKLYVVDFEDVMLEEAVARGLVEDRDKIRTLPLDILRELQK
LAALRIRREALLALGASGILVVDTHALVKTVAGYYPGLPKFVMDILKPDMIAVVEASPEEVAARQARDTTRYRV
DIGGVEGVKRLMENARAASIASAIQYASTVAIVENREGEAAKAAEELLRLIKNL
T33-ml34-A
MHHHHHHGGSHHWGGMSISYRKLDIALSADGEEVLVDGQVLPTRFFLDTVVTTMLNACGTDEENINEILADV
HAAGLDVSNYGWASEVYKKGDPEKRAEAEARRAEAEARRAERRRRLASPEARRERRREEAERRLRLYERF
GEAGLEAGL
T33-ml34-B
MTTEEEVVLAIAELFLPDPHARAEAAKKLGKIGDPEAVPALIRALFDPDPAVRAAAAKALGKIGDKEAVPALIVAL
FDPDPAVRVAAAKALGKIGDKEAVPALIEALFDPDPAVRVAAAIALGKIGDKEAVPALVRALKYEEGLVREAAAI
ALKKIGGEEVKKAMEELAKFGEGEAKEFAEEYLKEN
T33-ml35-A

MNLAEKMYKAGQIEFAKGNYETAIIAYTLALLKDPNNAEAWYNLGEAYLALGNYEEAIEAYQKALELDPNNAEA
WYNLGEAYLALGDYDNAIEAFTKALELDPNNKTAKAGLKLAKEKKALE
T33-ml35-B
MTDLSSLIETADLRLLLTTVPTETEALYLALAAVEKGLAAEVLITPVTRVRRENGKLVVEDVYRLSFKTTRERLD
ALVAWLQRRHPLALPECLVLTPIASSVAYRDWLRSSLQGGSHHWGGHHHHHH
T33-ml36-A
MNMARDFYRAGLIAYAKGEYETAIVAFQLALLLDPNNAEAWYNLGKAYYALGLYREAIEAYKKALELDPNNAE
AWYNLGKAYYALGDYESAIEAYKKALELDPNNVEAHANLHKAKKKLALE
T33-ml36-B
MPSYAVSSRAGLIDQERRAAVADLITALHSEILKIPRYLVQVIFNDLDAGALFLAGREAPEGHVWIHADIISGRTK
EQKKAFLQALTVEVARVLGLPEEQVWVYVNEIPGENMTLFGQILPAPGEEEAWFATLPEELQKRLADLRGGS
HHWGGHHHHHH
T33-ml37-A
MNLANDFYEAGKEEFAKGRYNLAIVCFSLALLKDPNNAEAWYNLGKAYFALGKYDKAIEAYQKALELDPNNAE
AWYNLGLAYFALGNYKEAIEYYKKALELDPNNELAKLALKLAKEKLELE
T33-ml37-B
MAMPAVKLVIVTEKILLKDITRIILESGAKGFTVMNTGGIGSRERAGEGEPDIDKIRANIKFEVLCESRELAELIAE
AIASKFFDKYAGIIYTCSAEVLYGHDFCGPEGSGSHHWGGHHHHHH
T33-ml38-A
MHHHHHHGGSHHWGGMNLRAAGPGWLFCPAHRPELFAKAAAAADVVILDLEDGVAESMKPGARENLRAHP
LDPERTVVRINAGGTADQARDLEALAGTAYTTVMLPKAESAAQVIELAPRDVIALVETARGAVCAAEIAAADPT
VGMMWGAEDLIATLGGSSSRRADGAYRDVARHVRSTILLAASAFGRLALDAVHLDILDVEGLQEEARDAAAV
GFDVTVCIHPSQIPVVRKAYRPSHEKLEWARLVLLNAQGKAGAFVFEGQMVDSPVLTHAETMLRRAGEATSE
T33-ml38-B
MPSYAVSSRAGLIDRLRRLEVARLLTTLHRDIAVAPRYLVQVIFNDLDAGALFVAGAEAPEGHVWIHADIRSGR
TAQQKTDLLEQITSKVADVLELPPEHVWVYVNEIPGENMTEYGKLLPEPGKEEEWFATLPPGLQTVLSA
